# Accurate Characterization of Conformational Ensembles and Binding Mechanisms of the SARS-CoV-2 Omicron BA.2 and BA.2.86 Spike Protein with the Host Receptor and Distinct Classes of Antibodies Using AlphaFold2-Augmented Integrative Computational Modeling

**DOI:** 10.1101/2023.11.18.567697

**Authors:** Nishank Raisinghani, Mohammed Alshahrani, Grace Gupta, Sian Xiao, Peng Tao, Gennady Verkhivker

**Affiliations:** Keck Center for Science and Engineering, Graduate Program in Computational and Data Sciences, Schmid College of Science and Technology, Chapman University, Orange, CA 92866, United States of America; Department of Biomedical and Pharmaceutical Sciences, Chapman University School of Pharmacy, Irvine, CA 92618, United States of America; Department of Chemistry, Center for Research Computing, Center for Drug Discovery, Design, and Delivery (CD4), Southern Methodist University, Dallas, Texas, 75275, United States of America

## Abstract

The latest wave SARS-CoV-2 Omicron variants displayed a growth advantage and the increased viral fitness through convergent evolution of functional hotspots that work synchronously to balance fitness requirements for productive receptor binding and efficient immune evasion. In this study, we combined AlphaFold2-based structural modeling approaches with all-atom MD simulations and mutational profiling of binding energetics and stability for prediction and comprehensive analysis of the structure, dynamics, and binding of the SARS-CoV-2 Omicron BA.2.86 spike variant with ACE2 host receptor and distinct classes of antibodies. We adapted several AlphaFold2 approaches to predict both structure and conformational ensembles of the Omicron BA.2.86 spike protein in the complex with the host receptor. The results showed that AlphaFold2-predicted conformational ensemble of the BA.2.86 spike protein complex can accurately capture the main dynamics signatures obtained from microscond molecular dynamics simulations. The ensemble-based dynamic mutational scanning of the receptor binding domain residues in the BA.2 and BA.2.86 spike complexes with ACE2 dissected the role of the BA.2 and BA.2.86 backgrounds in modulating binding free energy changes revealing a group of conserved hydrophobic hotspots and critical variant-specific contributions of the BA.2.86 mutational sites R403K, F486P and R493Q. To examine immune evasion properties of BA.2.86 in atomistic detail, we performed large scale structure-based mutational profiling of the S protein binding interfaces with distinct classes of antibodies that displayed significantly reduced neutralization against BA.2.86 variant. The results quantified specific function of the BA.2.86 mutations to ensure broad resistance against different classes of RBD antibodies. This study revealed the molecular basis of compensatory functional effects of the binding hotspots, showing that BA.2.86 lineage may have primarily evolved to improve immune escape while modulating binding affinity with ACE2 through cooperative effect of R403K, F486P and R493Q mutations. The study supports a hypothesis that the impact of the increased ACE2 binding affinity on viral fitness is more universal and is mediated through cross-talk between convergent mutational hotspots, while the effect of immune evasion could be more variant-dependent.

## Introduction

The wealth of structural and biochemical studies on the SARS-CoV-2 viral spike (S) glycoprotein has provided critical insights into mechanisms of virus transmission and immune resistance.^1-9^ The conformational changes within the SARS-CoV-2 S protein, transitioning between closed and open states, are primarily driven by the global movements of a flexible amino (N)-terminal S1 subunit of the S protein that includes an N-terminal domain (NTD), the receptor-binding domain (RBD), and two structurally conserved subdomains—SD1 and SD2. These structural domains coordinate their dynamic changes with a structurally rigid carboxyl (C)-terminal S2 subunit to facilitate diverse functional responses of the S protein through conformational transitions between the RBD-down closed and RBD-up open states. A complex and synchronized interplay of functional motions in the NTD, RBD, and SD1 and SD2 subdomains mediates essential interactions of the S protein with the host cell receptor ACE2 and a broad spectrum of antibodies, influencing the immune responses triggered by the host.^10-15^. Biophysical studies offered comprehensive insights into the thermodynamics and kinetics governing the behavior of the SARS-CoV-2 S protein trimer.^16-18^ These studies revealed how mutations and long-range interactions can orchestrate coordinated structural alterations in the dynamic S1 subunit and the more rigid S2 subunit, thereby modulating population shifts between the open and closed conformations of the RBD, which is integral to the S protein interaction with different binding partners, including the host cell receptor ACE2.^16-18^ The increasing availability of cryo-electron microscopy (cryo-EM) and X-ray structures of the SARS-CoV-2 S protein variants of concern (VOCs) in diverse functional states and in association with antibodies has offered a deeper understanding of the remarkable adaptability of molecular mechanisms. These studies have unveiled that VOCs can induce structural changes in the dynamic equilibrium of the S protein, affect population of functional states and create the diversity of binding epitopes that contribute to the varying binding affinities of S proteins when interacting with different classes of antibodies.^19-28^ Structural studies of the BA.2 and BA.3 RBD-ACE2 complexes suggested a stronger binding of BA.2 variant as compared to BA.3 and BA.1 variants.^29-31^ The cryo-EM structures and biochemical analysis of the S trimers for BA.1, BA.2, BA.3, and BA.4/BA.5 subvariants of Omicron demonstrated the decreased binding affinity for the BA.4/BA.5 and confirmed the higher binding affinities for BA.2 as compared to other Omicron variants.^32,33^ Structural and biophysical studies of the Omicron BA.2.75 variant reported thermal stabilities of the Omicron variants at neutral pH, showing that the BA.2.75 S-trimer was the most stable, followed by BA.1, BA.2.12.1, BA.5 and BA.2 variants, exhibiting a 9-fold enhancement of the binding affinity with ACE2 as compared to its parental BA.2 variant and significant antibody evasion.^34-36^ Additionally, surface plasmon resonance (SPR) for multiple Omicron subvariants—BA.1, BA.2, BA.3, BA.4/5, BA.2.12.1, and BA.2.75—revealed that the BA.2.75 subvariant exhibited a notably higher binding affinity, approximately 4-6 times greater than the other Omicron variants.^35^ Functional studies of the Omicron BA.1, BA2, BA.3, BA.4/BA.5 and BA.2.75 variants revealed a common trend wherein the acquisition of specific mutations that potentially promote immune evasion, possibly at the cost of reduced ACE2 affinity (and therefore potentially decreased infectivity), is often counterbalanced by the emergence of compensatory mutations aiming to restore or enhance ACE2 binding, thus potentially increasing the virus’s ability to bind and enter host cells.^37-44^ The observed intricate dance of mutations, where certain changes favor immune evasion at the potential expense of decreased ACE2 affinity, followed by compensatory mutations that bolster ACE2 binding, highlighted the adaptability of the virus in response to evolutionary pressures. These studies suggested that evolution of Omicron variants is a dynamic process where different mutations may arise and interact, potentially impacting the virus’s ability to evade the immune system while maintaining or enhancing its ability to infect host cells.

A further striking demonstration of Omicron evolution was the emergence of XBB.1 subvariant is a descendant of BA.2 and recombinant of BA.2.10.1 and BA.2.75 sublineages. XBB.1.5 is very similar to XBB.1 with a single RBD modification which is a notably rare two nucleotide substitution compared with the ancestral strain.^45^ The biophysical studies of the S trimer binding with ACE2 for BA.2, BA.4/5, BQ.1, BQ.1.1, XBB, and XBB.1, variants showed that the binding affinities of BQ.1 and BQ.1.1 were comparable to that of BA.4/5 spike, while binding XBB and XBB.1 was similar to that of BA.2 variant.^46^ This study reported an alarming antibody evasion by Omicron BQ.1, BQ.1.1, XBB, and XBB.1 variants showing that monoclonal antibodies capable of neutralizing the original Omicron variant were largely inactive against these new subvariants.^46^ XBB.1.5 is equally immune evasive as XBB.1 but may have growth advantage by virtue of the higher ACE2 binding affinity owing to a single S486P mutation as F486S substitution in XBB.1 variant.^47,48^ The biochemical studies examined the binding affinity of the XBB.1.5 RBD to ACE2 revealing the dissociation constant K_D_ = 3.4 nM which was similar to that of BA.2.75 (K_D_=1.8 nM) while significantly stronger than that of XBB.1 and BQ.1.1 variants.^47^ Cryo-EM analysis of both the XBB.1.5 S ectodomain confirmed that XBB.1.5 is equally immune evasive as XBB.1 but exhibits the higher ACE2 binding affinity resulting in the growth advantage and the increased transmissibility of the XBB.1.5 variant.^49^

The Omicron subvariant BA.2.86 was identified by global genomic surveillance in late August 2023 and exhibits significant genetic differences compared to its predecessors.^50-54^ BA.2.86 is derived from BA.2 variant but has 34 mutations relative to BA.2 (29 substitutions, 4 deletions and 1 insertion) including RBD mutations I332V, D339H, K356T, R403K, V445H, G446S, N450D, L452W, N460K, N481K, delV483, A484K, F486P and R493Q (Figure 1).^51-53^

**Figure 1.**
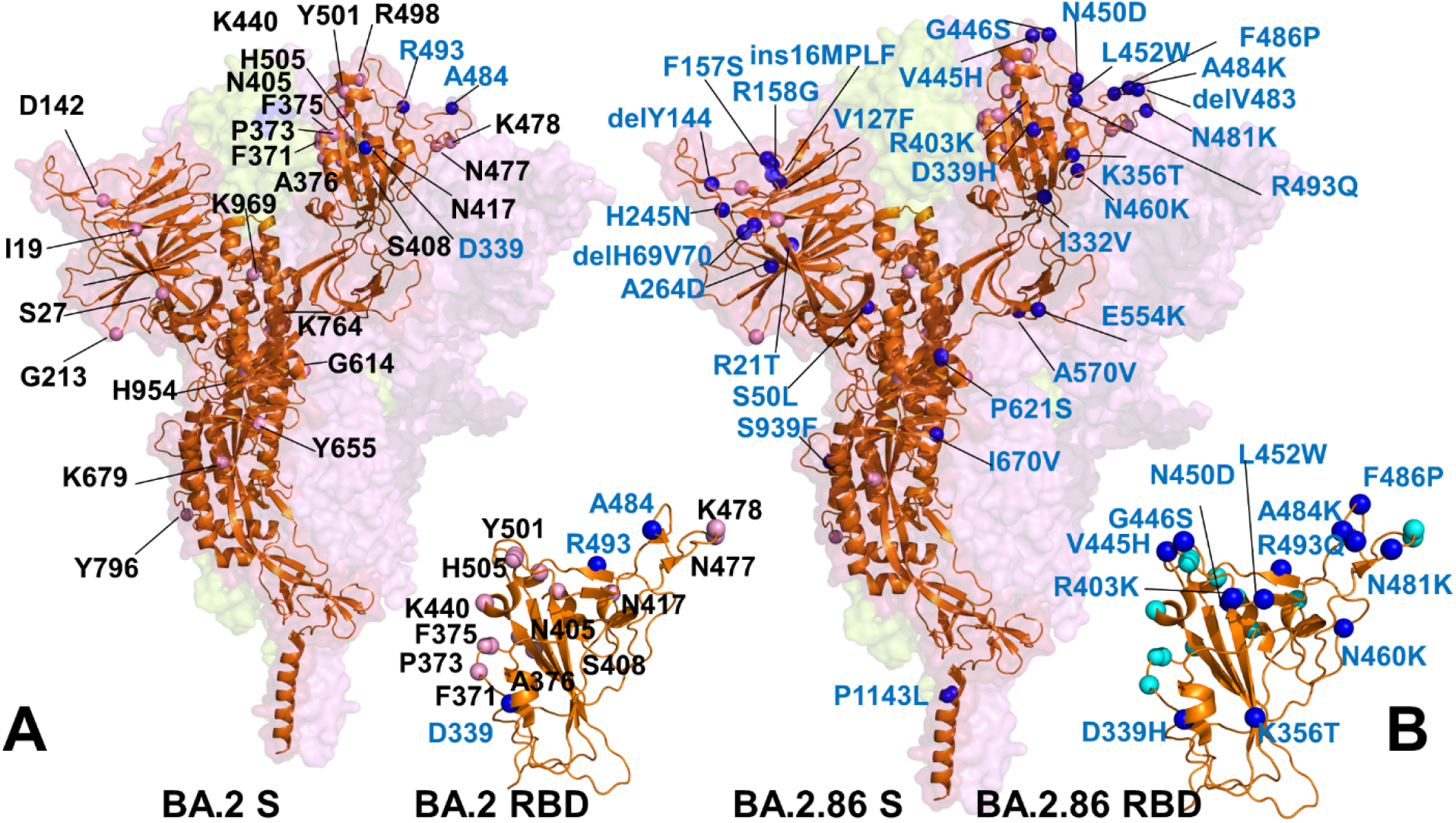
Structural overview of the SARS-CoV-2 S protein and S-RBD for Omicron BA.2 (A) and BA.2.86 variants (B). The S protein is shown on orange ribbons (single monomer) with the S protein trimer structure shown in surface with the reduced transparency. For BA.2 S protein all BA.2 mutational sites are shown in pink-colored spheres and annotated. The positions of BA.2 mutations D339, A484 and R493 mutations are shown in blue spheres. The BA.2 RBD mutations are also shown projected onto crystallographic RBD conformation (orange ribbons) in the BA.2 RBD-ACE2 complex, pdb id 7XB0 (A). The positions of unique BA.2.86 S mutations relative to its ancestral BA.2 variant are shown in blue-colored spheres and fully annotated. The AF2-generated BA.2.86 RBD model (in orange ribbons) and BA.2.86 RBD mutations in blue spheres are shown and annotated.

BA.2.86 is also quite different from XBB.1.5 with 36 unique mutations including 32 substitutions, 3 deletions and 1 insertion. Along with the shared mutations with XBB.1.5 (T19I, 24-26del, A27S, G142D, 144del, G339H, G446S, N460K, and F486P), additional mutations I332V, K356T, V445H, N450D, N481K, A484K and 483del on BA.2.86’s RBD are likely to enhance immune evasion.^51-53^ Some of these mutations such as K356T, R403K, V445H, N450D, L452W, delV483 and A484K differentiate BA.2.86 from both BA.2 and XBB.1.5 sublineage and are unique to this specific variant (Figure 1, Table 1).

**Table 1.** Mutational landscape of the Omicron BA.1, BA.2, XBB.1.5 and BA.2.86 variants.*

| Omicron Variant | Mutational landscape |
| --- | --- |
| BA.1 | A67, T95I, G339D, S371L, S373P, S375F, K417N, N440K, G446S, S477N, T478K, E484A, Q493R, G496S, Q498R, N501Y, Y505H, T547K, D614G, H655Y, N679K, P681H, N764K, D796Y, N856K, Q954H, N969K, L981F |
| BA.2 | T19I, G142D, V213G, G339D, S371F, S373P, S375F, T376A, D405N, R408S, K417N, N440K, S477N, T478K, E484A, Q493R, Q498R, N501Y, Y505H, D614G, H655Y, N679K, P681H, N764K, D796Y, Q954H, N969K |
| XBB.1.5 | T19I, V83A, G142D, Del144, H146Q, Q183E, V213E, G252V, G339H, R346T, L368I, S371F, S373P, S375F, T376A, D405N, R408S, K417N, N440K, V445P, G446S, N460K, S477N, T478K, E484A, F486P, F490S, R493Q reversal, Q498R, N501Y, Y505H, D614G, H655Y, N679K, P681H, N764K, D796Y, Q954H, N969K |
| BA.2.86 | Ins16MPLF, R21T, S50L, del69-70, V127F, delY144, F157S, R158G, delN211, L213I, L226F, H25N, A264D, I332V, D339H, K356T, R403K, V445H, G446, N450D, L452W, N460K, del V483, A484K, F486P, R493Q, E554K, A570V, P612S, I670V, H68R, D939F, P1143L |

\*For BA.2.86 variant only mutations that are different from its ancestral BA.2 are listed.

Structural mapping of BA.2 and BA.2.86 specific mutations onto the S protein showed the distribution of mutational sites (Figure 1C,D) highlighting the unique BA.2.86 positions particularly in the RBD region near the binding interface with the ACE2 host receptor. BA.2.86 also has as many as 58 mutations relative to the early Wu-Hu-1 variant (with 52 substitutions, 5 deletions and 1 inser1 insertion. A significant divergence of BA.2.86 subvariant is exemplified by the fact that the genetic distance between BA.2.86 and its predecessor, BA.2, is similar to the distance observed between the BA.1 variant and the Delta variant of the virus (Supporting Information, Figure S1).^50,51^ Overall, the number of different mutations in the BA.2.86 variant relative to BA.2 and XBB.1.5 is comparable to number of mutations in the initial Omicron strains relative to the original Wu-Hu-1 strain.

Biophysical studies used SPR techniques to measure ACE2 binding affinities showing that XBB.1.5 and EG.5.1 spikes exhibited comparable affinities to ACE2, with K_D_ values of 1.34 nM and 1.21 nM, respectively as compared to the K_D_ value of the BA.2 spike (1.68 nM). In contrast, both constructs of the BA.2.86 S protein showed a >2-fold increase in binding affinity, with similar K_D_ values of 0.54 nM and 0.60 nM.^51^ This study provided the first detailed account of the antibody evasion properties of BA.2.86 by evaluating thee susceptibility to neutralization by a panel of 25 monoclonal antibodies that retained activity against BA.2, XBB.1.5 and EG.5.1 variants, with 20 of these antibodies targeting the four different RBD epitope classes. In particular, this study established that BA.2.86 variant can be resistant to neutralization by monoclonal antibodies to NTD, SD1, and also RBD classes 1, 2 and class 3 epitopes, and the evasion potential from RBD-targeted antibodies is larger than the corresponding immune escape exhibited by XBB.1.5 and EG.5.1 variants.^51^

Antigenicity and immune evasion capability of the BA.2.86 subvariant was tested on a panel of XBB.1.5-effective neutralizing antibodies revealing that BA.2.86 is antigenically distinct from XBB.1.5 and previous Omicron variants and can escape XBB-induced neutralizing antibodies due to N450D, K356T, V445H, L452W, A484K, V483del, and A484K mutations.^52^ These findings suggest a complex interplay between the immune evasion capabilities of the BA.2.86 subvariant and the efficacy of certain vaccines. The neutralizing antibody responses against the BA.2.86 variant were considerably lower compared to the responses observed against the BA.2 variant but comparable to those observed against the XBB.1.5, XBB.1.16, EG.5, EG.5.1, and FL.1.5.1 variants.^53^ This study suggested that the BA.2.86 variant likely evolved directly from the less resistant BA.2 variant, rather than originating from the more highly resistant circulating recombinant variants. Functional studies examined whether BA.2.86 may exhibit growth advantages over other currently circulating Omicron variants, including EG.5.1 and the FLip variant, which contains the L455F and F456L mutation in the background of XBB.1.5 variant.^54^ Antigenic mapping showed that BA.2.86 may exhibit similar immune evasion as the XBB variants while it is more antigenically similar to early Omicron subvariants BA.1, BA.2, and BA.4/5 and antigenically distinct from the FLip variant.^54^ Functional analysis of transmissibility, infectivity, and immune resistance of the BA.2.86 variant suggested BA.2.86 potentially has greater fitness than current circulating XBB variants including EG.5.1 while the infectivity of BA.2.86 was significantly lower than that of B.1.1 and EG.5. ^55^ Neutralization assay using XBB breakthrough infection sera examined showed that BA.2.86 evades the antiviral effect of the humoral immunity induced by XBB subvariants, suggesting that BA.2.86 is among the most highly immune evasive variants ever.^55^

Comprehensive multiscale investigations of the virological characteristics of the BA.2.86 variant demonstrated that the ACE2 binding affinity of BA.2.86 S RBD was comparable to that of XBB.1.5 S RBD and significantly higher than those of the ancestral B.1.1, XBB.1, XBB.1.16, EG.5.1 and the parental BA.2 variants.^56^ This illuminating study showed that fusogenicity of the S protein of BA.2.86 and BA.2 were comparable, but the intrinsic pathogenicity of BA.2.86 was significantly lower than that of BA.2, suggesting that the attenuated pathogenicity of BA.2.86 may be due to its decreased replication capacity.^56^

Computer simulation studies provided important atomistic insights into understanding the dynamics of the SARS-CoV-2 S protein and the effects of Omicron mutations on conformational plasticity of the S protein states and their complexes with diverse binding partners. Molecular dynamics (MD) simulations of the full-length SARS-CoV-2 S glycoprotein with a complete glycosylation profile provided detailed characterization of the conformational landscapes of the S proteins in the physiological environment.^57-61^ Large-scale adaptive sampling simulations of the viral proteome captured the conformational heterogeneity of the S protein and predicted the existence of multiple cryptic epitopes and hidden allosteric pockets.^62^ Replica-exchange molecular dynamics (MD) simulations examined the conformational landscapes of full-length S protein trimers, discovering transition pathways via inter-domain interactions, hidden functional intermediates along open-closed transition pathways, and previously unknown cryptic pockets that were consistent with FRET experiments.^63,64^ Our recent studies demonstrated that Omicron mutational sites can be dynamically coupled forming an adaptive allosteric network that controls balance and tradeoffs between conformational plasticity, protein stability, and functional adaptability.^65--69^ We combined MD simulations and Markov state models to systematically characterize conformational landscapes and identify specific dynamic signatures of the early Omicron variants BA.1, BA.2, BA.3 and BA.4/BA.5 variants^70^ and recent highly transmissible XBB.1, XBB.1.5, BQ.1, and BQ.1.1 Omicron variants and their complexes.^71^ A significant number of computational studies emphasized the role of electrostatic interactions as a dominant thermodynamic force leading at binding of the S-protein with the ACE2 receptor and antibodies.^72-75^ Recent studies mapped the electrostatic potential surface of S protein and major variants to show accumulated positive charges at the ACE2-binding interface, revealing the critical role of complementary electrostatic interactions driving the enhanced affinity the Omicron S-ACE2 complexes.^73^ Extensive simulations investigated the electrostatic features of the RBD for the several S variants, their main physical–chemical properties and binding affinities at several pH regimes revealing that the virus evolution may primarily exploit the electrostatic forces to make RBD more positively charged and improve the RBD-ACE2 binding affinity and transmission.^75^

The recent structural and computational studies suggested that Omicron mutations have significant and often variant-specific effect on mediating conformational dynamics changes in the S protein including allosterically induced plasticity at the remote regions leading to the formation and evolution of druggable cryptic pockets.^76,77^ Evolutionary trajectories of Omicron lineages proceeded through diverse mechanisms including but not limited to simple and complex recombination, antigenic drift and convergent evolution that led to convergent immune-escape mutations and effective evasion of neutralizing antibodies.^56^ The recent proliferation of several lineages with similar replicative fitness and significant antigenic distance from each other was marked by the recent emergence of BA.2.86. Although several investigations examined antigenicity and receptor binding signatures of the BA.2.86 variant, there is a conspicuous lack of the atomistic level information regarding the structure, dynamics, and binding mechanisms of the BA.2.86 RBD binding with ACE2 receptor and a wide spectrum of experimentally studied antibodies. To our knowledge, there have been no computational investigations conducted to date focusing on this particular variant.

In this study, we employed and synergistically combined AlphaFold2 (AF2) methodology^78,79^ with all-atom MD simulations and in silico mutational scanning of binding energetics and stability for AI-augmented atomistic predictions of the structure, dynamics, and binding of the Omicron BA.2 and BA.2.86 RBD complexes with ACE2 receptor. The remarkable success of AF2 technology which uses multiple sequence alignment (MSA) as input that captures conserved contacts between evolutionarily related sequences marked a revolutionary change in structural biology.^78,79^ The latest breakthrough in AI-based protein structure predictions is also high accuracy end-to-end family of transformer protein language model ESMFold for atomic level structure prediction directly from the individual sequence of a protein.^80^ Several studies found that sub-sampling the input MSAs and increasing the number of predictions leads to structural ensembles of physiologically-relevant conformations from the same sequence.^81-83^ Another AF2 adaption used a simple MSA subsampling method of clustering sequences to predict alternate states of proteins.^84,85^ We adapted and applied AF2 methodology using shallow MSA to predict structure and conformational variability of the BA.2.86 RBD and the RBD-ACE2 complexes. Using several structural alignment metrics including AF2-based predicted local distance difference test (pLDDT) metric, we can evaluate and rank structural models leading to a robust prediction of the BA.2.86 RBD-ACE2 structure and conformational ensembles. Microsecond atomistic MD simulations confirmed the AF2 predictions of conformational states revealing similarities and differences in conformational dynamics of BA.2 and BA.2.86 variants. We also perform an ensemble-based mutational scanning of the RBD residues in the BA.2 and BA.2.86 RBD-ACE2 complexes to characterize conserved and variant-specific binding energy hotspots that explain the molecular basis of binding affinity mechanism for the BA.2.86 variant. Our results show that binding affinity differences can be determined by common hydrophobic binding hotspots and variant-specific BA.2.86 mutational sites R403K, F486P and R493Q, while other BA.2.86 RBD mutations may have evolved to modulate immune resistance against diverse classes of RBD antibodies.

To understand the energetics of antibody evasion for BA.2.86 and characterize specific role of BA.2.86 mutations in enabling immune escape, we conduct a large scale structure-based mutational scanning of the S RBD complexes with 20 antibodies for the four epitope classes. The results detail energetic mechanisms by which BA.2.86 mutational sites can elicit resistance to neutralization the RBD antibodies. The results suggest a mechanism in which convergent Omicron mutations can promote high transmissibility and antigenicity of the virus by controlling the interplay between the RBD stability and conformational adaptability, allowing for optimal fitness tradeoffs between binding to the host receptor and robust immune evasion profile.

## Materials and Methods

### AI-based structural modeling and statistical assessment of AF2 models

Structural prediction of the BA.2.86 RBD and BA.2.86 RBD-ACE2 complex were carried out using AF2 framework^78,79^ within the ColabFold implementation^86^ using a range of MSA depths and other parameters.^81-83^ The default MSAs are subsampled randomly to obtain shallow MSAs containing as few as five sequences. We generated structures using shallow MSA depth by adjusting AF2 parameters using ColabFold.^86^ We used *max_msa* field to set two AF2 parameters in the following format: *max_seqs:extra_seqs*. Both of these determine the number of sequences subsampled from the MSA (*max_seqs* sets the number of sequences passed to the row/column attention track and *extra_seqs* the number of sequences additionally processed by the main evoformer stack). The lower values encourage more diverse predictions but increase the number of misfolded models. Similar to previous studies showing how MSA depth adaptations may facilitate conformational sampling,^81-83^ MSA depth was modified by setting the AF2 config.py parameters *max_extra_msa* and *max_msa_clusters* to 32 and 16, respectively. We additionally manipulated the *num_seeds* and the *num_recycles* parameters to produce more diverse outputs. We use *max_msa*: 16:32, *num_seeds*: 4, and *num_recycles*: 12. AF2 makes predictions using 5 models pretrained with different parameters, and consequently with different weights. To generate more data, we set the number of recycles to 12, which produces 14 structures for each model starting from recycle 0 to recycle 12 and generating a final refined structure. Recycling is an iterative refinement process, with each recycled structure getting more precise. Each of the AF2 models generates 14 structures, amounting to 70 structures in total. In addition, we also predicted one more structure using AF2 with the default and ‘auto’ parameters serving as a baseline structure for prediction and variability analysis.

AF2 models were ranked by Local Distance Difference Test (pLDDT) scores (a per-residue estimate of the prediction confidence on a scale from 0 to 100), quantified by the fraction of predicted Cα distances that lie within their expected intervals. The values correspond to the model’s predicted scores based on the lDDT-Cα metric, a local superposition-free score to assess the atomic displacements of the residues in the model.^78,79^ Structural models were compared to the experimental structure of BA.2 RBD-ACE2 (pdb id 7XB0) using structural alignment as implemented in TM-align.^87^ An optimal superposition of the two structures is then built and TM-score is reported as the measure of overall accuracy of prediction for the models. Several other structural alignment metrics were used including the global distance test total score GDT_TS of similarity between protein structures and implemented in the Local-Global Alignment (LGA) program^88^ and the root mean square deviation (RMSD) superposition of backbone atoms (C, Cα, O, and N) calculated using ProFit (http://www.bioinf.org.uk/software/profit/).

### All-Atom Molecular Dynamics Simulations

The crystal structure of the BA.2 RBD-ACE2 (pdb id 7XB0) is obtained from the Protein Data Bank and structures of the BA.2.86 RBD-ACE2 complex are obtained from AF2 modeling.^78,79^ We selected BA.2.86 RBD-ACE2 models based on the pLDDT scores, particularly of the RBM loop residues as the highest scoring models have the lowest RMSD to the cryo-EM and crystal structures of the RBD-ACE2 complexes for Omicron BA.1, BA.2, BA.3, BA.4/BA.5 and XBB.1 variants. Hydrogen atoms and missing residues were initially added and assigned according to the WHATIF program web interface.^89^ The missing regions are reconstructed and optimized using template-based loop prediction approach ArchPRED.^90^ The side chain rotamers were refined and optimized by SCWRL4 tool.^91^ The protonation states for all the titratable residues of the ACE2 and RBD proteins were predicted at pH 7.0 using Propka 3.1 software and web server.^92,93^ The protein structures were then optimized using atomic-level energy minimization with composite physics and knowledge-based force fields implemented in the 3Drefine method.^94,95^ We considered glycans that were resolved in the structures. The structurally resolved 2-acetamido-2-deoxy-beta-D-glucopyranose, 2-acetamido-2-deoxy-beta-D- glucopyranose-(1-4)-2-acetamido-2-deoxy-beta-D-glucopyranose and chloride ions present in the RBD-ACE2 structures were included and optimized.

NAMD 2.13-multicore-CUDA package^96^ with CHARMM36 force field^97^ was employed to perform 1µs all-atom MD simulations for the Omicron RBD-ACE2 complexes. The structures of the SARS-CoV-2 S-RBD complexes were prepared in Visual Molecular Dynamics (VMD 1.9.3)^98^ and with the CHARMM-GUI web server^99,100^ using the Solutions Builder tool. Hydrogen atoms were modeled onto the structures prior to solvation with TIP3P water molecules^101^ in a periodic box that extended 10 Å beyond any protein atom in the system. To neutralize the biological system before the simulation, Na^+^ and Cl^−^ ions were added in physiological concentrations to achieve charge neutrality, and a salt concentration of 150 mM of NaCl was used to mimic a physiological concentration. All Na^+^ and Cl^−^ ions were placed at least 8 Å away from any protein atoms and from each other. MD simulations are typically performed in an aqueous environment in which the number of ions remains fixed for the duration of the simulation, with a minimally neutralizing ion environment or salt pairs to match the macroscopic salt concentration.^102^ All protein systems were subjected to a minimization protocol consisting of two stages. First, minimization was performed for 100,000 steps with all the hydrogen-containing bonds constrained and the protein atoms fixed. In the second stage, minimization was performed for 50,000 steps with all the protein backbone atoms fixed and for an additional 10,000 steps with no fixed atoms. After minimization, the protein systems were equilibrated in steps by gradually increasing the system temperature in steps of 20 K, increasing from 10 K to 310 K, and at each step, a 1ns equilibration was performed, maintaining a restraint of 10 Kcal mol^−1^ Å^−2^ on the protein C atoms. After the restraints on the protein atoms were removed, the system was equilibrated for an additional 10 ns. Long-range, non-bonded van der Waals interactions were computed using an atom-based cutoff of 12 Å, with the switching function beginning at 10 Å and reaching zero at 14 Å. The SHAKE method was used to constrain all the bonds associated with hydrogen atoms. The simulations were run using a leap-frog integrator with a 2 fs integration time step. The ShakeH algorithm in NAMD was applied for the water molecule constraints. The long-range electrostatic interactions were calculated using the particle mesh Ewald method^103^ with a cut-off of 1.0 nm and a fourth-order (cubic) interpolation. The simulations were performed under an NPT ensemble with a Langevin thermostat and a Nosé– Hoover Langevin piston at 310 K and 1 atm. The damping coefficient (gamma) of the Langevin thermostat was 1/ps. In NAMD, the Nosé–Hoover Langevin piston method is a combination of the Nosé–Hoover constant pressure method^104^ and piston fluctuation control implemented using Langevin dynamics.^105,106^ An NPT production simulation was run on equilibrated structures for 1µs keeping the temperature at 310 K and a constant pressure (1 atm).

### Distance Fluctuations Stability Analysis

Our approach involved employing distance fluctuation analysis on simulation trajectories to derive residue-based stability profiles. This analysis computed the variations in the average distance between pseudo-atoms representing a specific amino acid and those belonging to the remaining protein residues. These distance fluctuations for each residue, concerning all other residues in the ensemble, were then translated into distance fluctuation stability indexes. These indexes serve as measurements quantifying the energy expenses associated with the deformation of each residue throughout the simulations.^107,108^ The adaptation of this approach for the analysis of rigid and flexible residues in the SARS-CoV-2 S proteins was detailed in our previous studies.^66-68^ The high values of distance fluctuation stability indexes point to structurally rigid residues as they display small fluctuations in their distances to all other residues, while small values of this index would point to more flexible sites that display larger deviations of their inter-residue distances. The distance fluctuation stability index for each residue is calculated by averaging the distances between the residues over the simulation trajectory as follows:

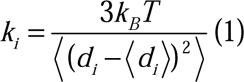

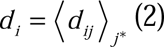

*^d^ ij* is the instantaneous distance between residue *i* and residue *j*, *k_B_* is the Boltzmann constant, *T* =300K. 〈 〉denotes an average taken over the MD simulation trajectory and 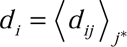 is the average distance from residue *i* to all other atoms *j* in the protein (the sum over *j*_*_ implies the exclusion of the atoms that belong to the residue *i*). The interactions between the *C_α_* atom of residue *i* and the *C_α_* atom of the neighboring residues and *i*+1 are excluded in the calculation since the corresponding distances are constant. The inverse of these fluctuations yields an effective force constant *k_i_* that describes the ease of moving an atom with respect to the protein structure.

### Mutational Scanning and Binding Free Energy Computations

The binding free energies were initially computed for the Omicron RBD-ACE2 complexes using the Molecular Mechanics/Generalized Born Surface Area (MM-GBSA) approach.^109,110^ We also conducted mutational scanning analysis of the binding epitope residues for the SARS-CoV-2 S RBD-ACE2 complexes. Each binding epitope residue was systematically mutated using all substitutions and corresponding protein stability and binding free energy changes were computed. BeAtMuSiC approach^111-113^ was employed that is based on statistical potentials describing the pairwise inter-residue distances, backbone torsion angles and solvent accessibilities, and considers the effect of the mutation on the strength of the interactions at the interface and on the overall stability of the complex. The binding free energy of protein-protein complex can be expressed as the difference in the folding free energy of the complex and folding free energies of the two protein binding partners:

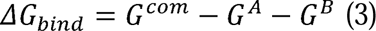

The change of the binding energy due to a mutation was calculated then as the following:

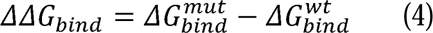

We compute the ensemble-averaged binding free energy changes using equilibrium samples from simulation trajectories. The binding free energy changes were computed by averaging the results over 1,000 equilibrium samples for each of the studied systems.

### Electrostatic Potential Calculations

In the framework of continuum electrostatics, the electrostatic potential for biological macromolecules can be obtained by solving the Poisson–Boltzmann equation (PBE)

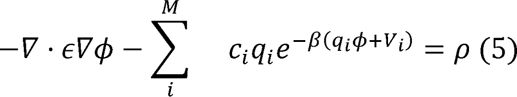

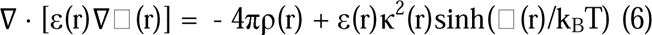

where LJ(r) is the electrostatic potential, ε(r) is the dielectric distribution, ρ(r) is the charge density based on the atomic structures, κ is the Debye–Huckel parameter, k_B_ is the Boltzmann constant, and T is the temperature. The electrostatic interaction potentials are computed for the averaged RBD-ACE2 conformations using the APBS-PDB2PQR software^114-116^ based on the Adaptive Poisson–Boltzmann Solver (APBS). These resources are available from the APBS/PDB2PQR website: http://www.poissonboltzmann.org/. The atomic charges and radii are assigned in this approach based on the CHARMM force field.

## Results

### AF2-Based Atomistic Modeling and Prediction of the BA.2.86 RBD-ACE2 Structure and Conformational Ensembles

Despite the established view that structural differences between SARS-CoV-2 S proteins for different variants are usually relatively moderate and the RBD-ACE2 complexes typically feature the same binding arrangement, it is possible that the considerable number of unique mutations in BA.2.86 may impose specific structural changes. To develop accurate and robust atomistic models of BA.2.86 RBD structure and dynamics, we performed comparative structural prediction of the BA.2 and BA.2.86 RBD-ACE2 complexes using AF2 default settings and AF2 methodology with shallow MSA depth^81-83^ within the ColabFold.^86^ We systematically tested the accuracy of predicting the BA.2 RBD-ACE2 structural ensembles by comparing the AF2- derived conformations with the structural and biophysical studies of the S Omicron protein dynamics and binding.^117-119^ First, we generated top five AF2 models of the BA.2 RBD-ACE2 complex with default settings (Supporting Information, Figure S2) showing an excellent structural alignment with the crystallographic BA.2 RBD conformation (pdb id 7XB0) with high confidence pLDDT values. All the top AF2 models of the BA.2 RBD displayed RMSD < 1.0 Å from the crystal structure demonstrating the ability of AF2 to accurately reproduce atomistic details of the BA.2 RBD (Supporting Information, Figure S2). Using this prediction as reference, we tested several combinations of parameters to employ AF2 with varied MSA depth. Te objective of this analysis was to go beyond prediction of the native state and characterize the RBD-ACE2 ensembles in which the crystallographic state is the most frequent prediction within the ensemble. Consistent with previous studies, we found changing *max_seq* and *extra_seq* allows to obtain predictions consistent with the crystallographic conformations with a *max_seq*:*extra_seq* ratio of 256:512 leading to the most diversity of the RBD loops and *max_seq*:*extra_seq* ratio sufficient to unequivocally reproduce the experimental structure using pLDDT metric (Figure 2). Strikingly, we found that the ensemble of AF2 models selected by the predicted local difference test (pLDDT) display the greatest structural similarity to the experimental structure. Using several structural similarity metrics such as AF2-based predicted pLDDT score^78,79^ TM-score,^87^ GDT_TS^88^ and RMSD we first examined and validated the prediction accuracy of AF2-MSA depth models for BA.2 RBD-ACE2 complex (Figure 2). The density distribution of the pLDDT values obtained for the AF2-MSA depth ensemble of BA.2 conformations showed a pronounced peak at pLDDT scores of ∼ 80-85 (Figure 2A).

**Figure 2.**
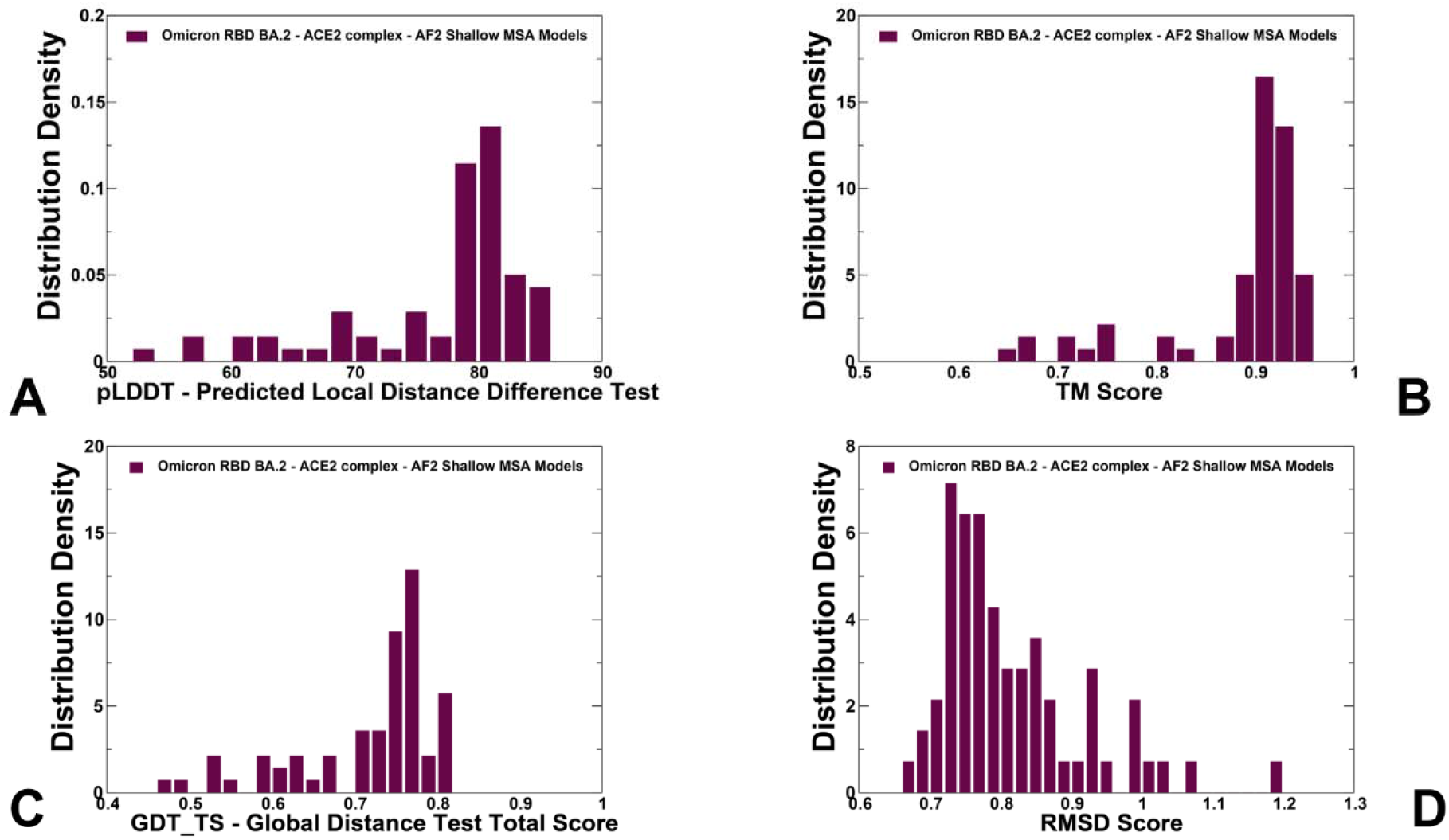
The distributions of structural model assessment and structural similarity metrics for the BA.2 RBD conformational ensemble obtained from AF2-MSA depth predictions. (A) The density distribution of the AF2-derived pLDDT structural model estimate of the prediction confidence on a scale from 0 to 100. (B) The density distribution of TM-score measuring structural similarity of the AF2-MSA depth predicted RBD conformations with respect to the crystal structure of the BA.2 RBD-ACE2 (pdb id 7XB0). (C) The density distribution of GDT_TS structural similarity metric between the AF2-predicted RBD conformations and the crystal structure of the BA.2 RBD-ACE2. (D) The density distribution of RMSD between the AF2-predicted RBD conformations and the crystal structure of the BA.2 RBD-ACE2.

The density distributions of other structural metrics TM-score (Figure 2B) and GDT_TS (Figure 2C) that measured similarity between the predicted conformations and the crystal structure similarly displayed sharp peaks signaling the consistent prediction of the crystallographic conformation, while also pointing to moderate variability reflecting fluctuations of the flexible RBD loops. In particular, the distribution featured a major peak for TM-score ∼ 0.9 confirming excellent predictions of the AF2-MSA depth model (Figure 2B). Several minor distribution peaks around TM-score ∼0.7-0.8 reaffirmed the quality of predictions that also reproduced the flexible RBD regions with high accuracy. The RMSD distribution showed that most of the predicted conformations are similar to the crystal structure with the main peak corresponding to RMSD ∼ 0.75 Å from the crystallographic state (Figure 2D). Overall, the predicted AF2-MSA depth models consistently produced highly accurate predictions of the BA.2 RBD-ACE2 structures.

Using the predictions of BA.2 RBD-ACE2 structure as a validation baseline, we also generated AF2 models for the BA.2.86 RBD and BA.2.86 RBD-ACE2 complex (Supporting Information, Figure S3). The alignment of top five BA.2.86 models with the crystallographic conformation o the BA.2 RBD showed only small deviations that are primarily associated with the functionally relevant plasticity of the RBM tip loop (residues 475-487) (Supporting Information, Figure S3). Structural mapping of the BA.2.86 mutational sites showed that their backbone positions remain unchanged and most variability may be expected in N481K and A484K sites. We also presented the statistical analysis and confidence assessment metrics of the BA.2.86 RBD predictions (Supporting Information, Figure S4). The multiple sequence alignment summarized as a heatmap indicating all sequences mapped to the input sequences. The relative coverage of the sequence with respect to the total number of aligned sequences is shown indicating reduced sequence identity to query for flexible RBD regions (residues 480-530) (Supporting Information, Figure S4A). The predicted LDDT per residue for the top five models showed that the RBD core and functionally important mobile regions featured pLDDT values within 70-90 range, signaling fairly high confidence of the predictions (Supporting Information, Figure S4B). The regions with pLDDT values ∼ 50-70 have lower confidence and must be treated with caution, while pLDDT < 50 may be a strong predictor of disorder (Supporting Information, Figure S4B). The heat maps of the predicted alignment error (PAE) between each residue in the model are shown for the best AF2 five models (Supporting Information, Figure S4C), showing differences between the high confidence regions and the low confidence regions. It is apparent from this analysis that flexible RBD loops 381-394 and especially 475-487 represent regions of lower confidence resulting from their inherent plasticity. Consistent with the ranking of the models, the lower confidence regions are moderately expanded in the flexible loop regions for models 4 and 5 (Supporting Information, Figure S4C).

We then proceeded to generate AF2 MSA depth models for the BA.2.86 RBD-ACE2 complex and analyzed structural similarities between the optimized AF2-default model of the BA.2.86 RBD-ACE2 complex and generated conformations (Figure 3). The distribution of pLDDT scores yielded a peak at pLDDT ∼ 82-85 reflecting the prediction convergence (Figure 3A). The distribution densities of structural similarity metrics obtained for the BA.2.86 RBD conformational ensemble showed similar trends to the ones seen for the BA.2 variant, featuring pronounced peaks for TM-score ∼0.9-0.95 (Figure 3B), GDT_TS ∼ 0.9 (Figure 3C) and for RMSD ∼ 0.65 Å from the reference structure (Figure 3D). Hence, the majority of predicted BA.2.86 conformations using AF2 with MSA depth are similar to the reference AF2-generated BA.2.86 RBD state and also crystallographic conformation of the BA.2 RBD-ACE2 complex.

**Figure 3.**
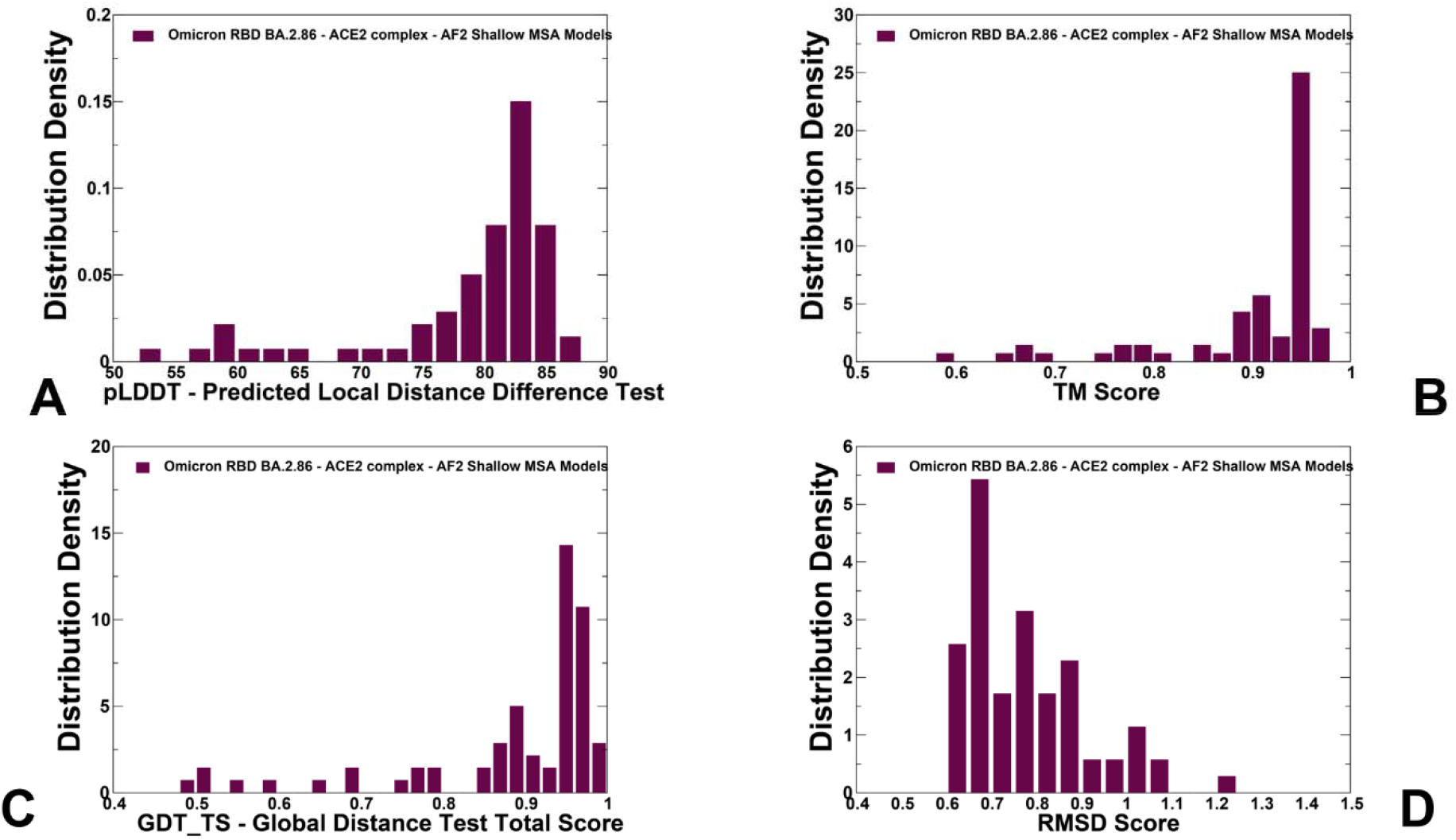
The distributions of structural model assessment and structural similarity metrics for the BA.2.86 RBD conformational ensemble obtained from AF2-MSA depth predictions. (A) The density distribution of the AF2-derived pLDDT structural model estimate of the prediction confidence. (B) The density distribution of TM-score measuring structural similarity of the AF2- MSA depth predicted RBD conformations with respect to the optimized AF2-generated reference state (C) The density distribution of GDT_TS structural similarity metric between the AF2-predicted RBD conformations and the optimized AF2-generated reference state. (D) The density distribution of RMSD between the AF2-predicted RBD conformations and the optimized AF2-generated reference state.

Structural alignment of the BA.2 RBD-ACE2 crystal structure and the refined AF2-default model of the BA.2.86 RBD-ACE2 complex (Figure 4A) illustrated a considerable similarity of the RBD conformations showing minor displacements in the intrinsically flexible RBM tip loop (residues 475-487) and in peripheral region (residues 520-527). These results are consistent with a significant body of cryo-EM and X-ray studies of Omicron S proteins and RBD-ACE2 complexes showing that VOC’s and Omicron lineages typically induce small structural changes but may affect the dynamics and binding energetics with the host receptor. Despite a substantial number of BA.2.86 unique RBD mutations relative to BA.2, the AF2-generated structural model of the BA.2.86 RBD is remarkably similar to the parental BA.2 RBD-ACE2 crystal structure, with only minor fluctuations in the RBM tip region (Figure 4A). Structural mapping of the unique BA.2.86 RBD mutations on the AF2-predicted conformations showed that these substitutions can induce only minor structural perturbations of the RBD backbone, including positions of A484K and F486P residues in the RBM region (Figure 4A). A more detailed inspection of the structural differences in the side chains of the RBD binding site residues depicted more appreciable variations of the positively charged side chains for N440K, N460K, N481K and also H505 (Figure 4B).

**Figure 4.**
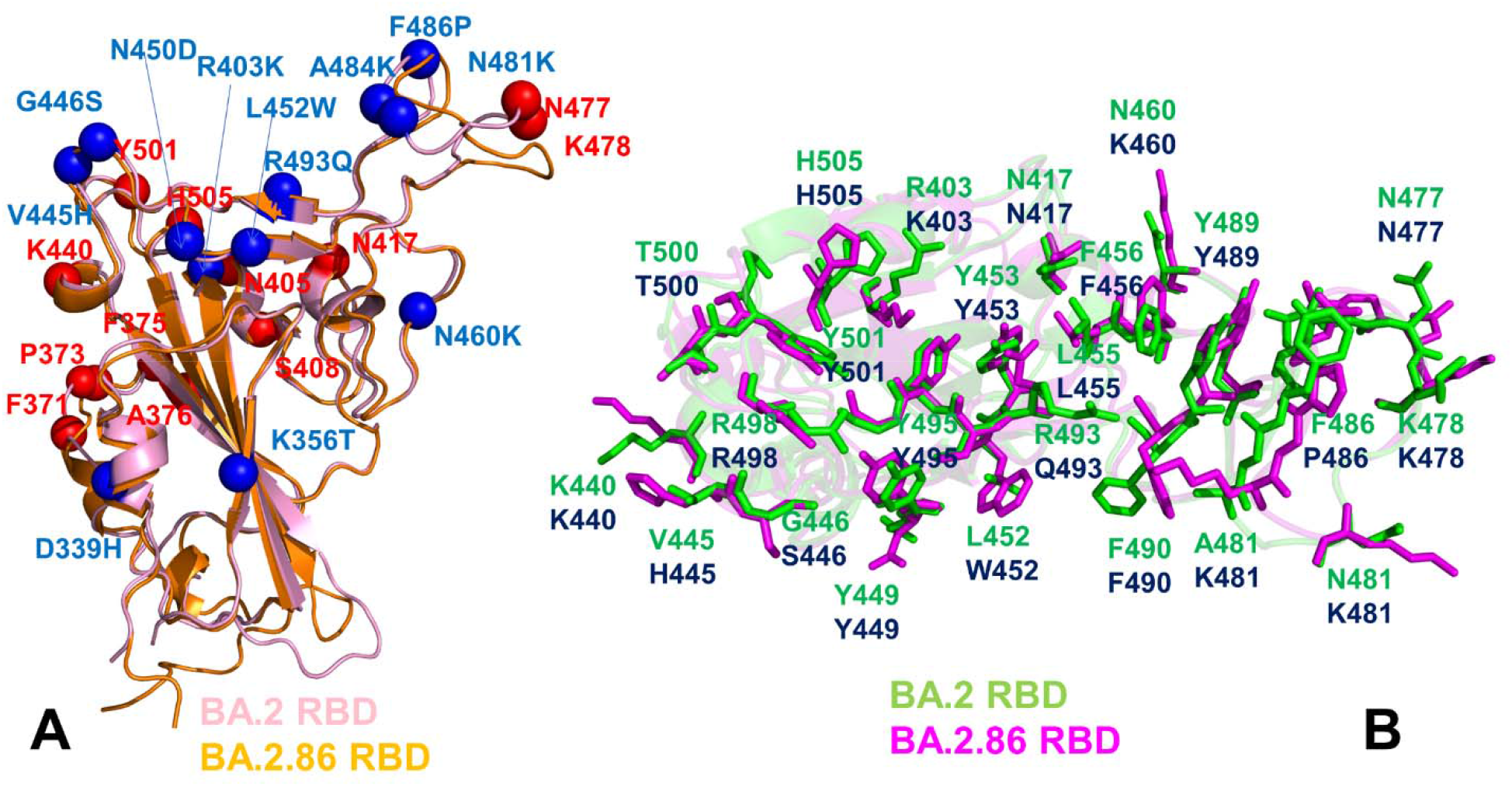
Structural alignment of the BA.2 and BA.2.86 RBD conformations from complexes with the ACE2 receptor. (A) Structural alignment of the crystallographic BA.2 RBD conformation (in light pink ribbons) and the refined AF2-predicted conformation of the BA.2.86 RBD (in orange ribbons). The BA.2 mutational sites are shown in red spheres and unique BA.2.86 mutations (D339H, R403K, V445H, G446S, N450D, L452W, N460K, N481K, A484K, F486P, R493Q). (B) Structural alignment of the RBD binding site residues making contacts with the ACE2 receptor. The structural positions of the BA.2 RBD binding site residues (in green sticks) are from the crystal structure of the BA.2 RBD-ACE2 complex. The structural arrangement of the BA.2.86 RBD binding sites residues are overlayed onto BA.2 RBD and are shown in magenta sticks. The RBD binding site residues for both BA.2 and BA.2.86 variants are annotated.

Importantly, more significant and functionally important for ACE2 binding displacements were observed for side chains of specific BA.2.86 mutational sits R403K, V445H, N450D, L452W, A484K reversal R493Q, and F486P (Figure 4B). These results showed that AF2-generated predictions of the BA.2 and BA.286 RBD-ACE2 complexes can accurately reproduce the experimental structures and capture conformational details of the RBD fold and variant-specific functional adjustments of the RBD binding site residues.

To illustrate the performance of the AF2-MSA depth approach in characterizing conformational ensembles we performed structural alignment of the predicted BA.2 RBD conformations with high pLDDT values and the crystal structure of the BA.2 RBD-ACE2 complex (Figure 5A). A high degree of similarity (RMSD < 1.0 Å) illustrated the ability of AF2-MSA models to accurately reproduce the experimental structure. Strikingly, we observed that selection of the BA.2 RBD models based on the high pLDDT scores not only can reproduce the RBD fold but also accurately predict conformations of the RBD flexible loops (444-452, 455-472) and capture moderate yet functionally relevant plasticity in the RBD loop 444-452, RBM tip (residues 475-487) that harbor important mutational sites and peripheral flexible region (residues 515-530). These regions are inherently flexible as revealed in hydrogen/deuterium-exchange mass spectrometry (HDX-MS) studies that informed of residue-specific changes in conformational dynamics induced by mutations and binding of SARS-CoV-2 S protein.^116-119^ HDX studies discovered that the RBM region of the S proteins could feature bimodal isotopic distributions between two distinct and slowly interconverting populations of ACE2-bound structurally stable and more flexible RBM conformations.^117^ Our predictions showed that the position of functionally important for ACE2 binding and immune escape F486 residue in BA.2 RBD (F486P in XBB.1.5 and BA.2.86 variants) is preserved despite functional displacements of the RBM loop (Figure 5A). The predicted conformations of other RBD loops (355-375, 381- 394, 455-471) are identical to the ones in the crystallographic conformation (Figure 5A). A similar structural alignment of the AF2-MSA depth ensemble was seen in the BA.2.86 RBD (Figure 5B), revealing appreciable but more homogeneous displacements of the flexible RBM tip. Interestingly, the deletion of V483 in BA.2.86 caused minor conformational changes in the local mobility of the RBM loop (Figure 5B).

**Figure 5.**
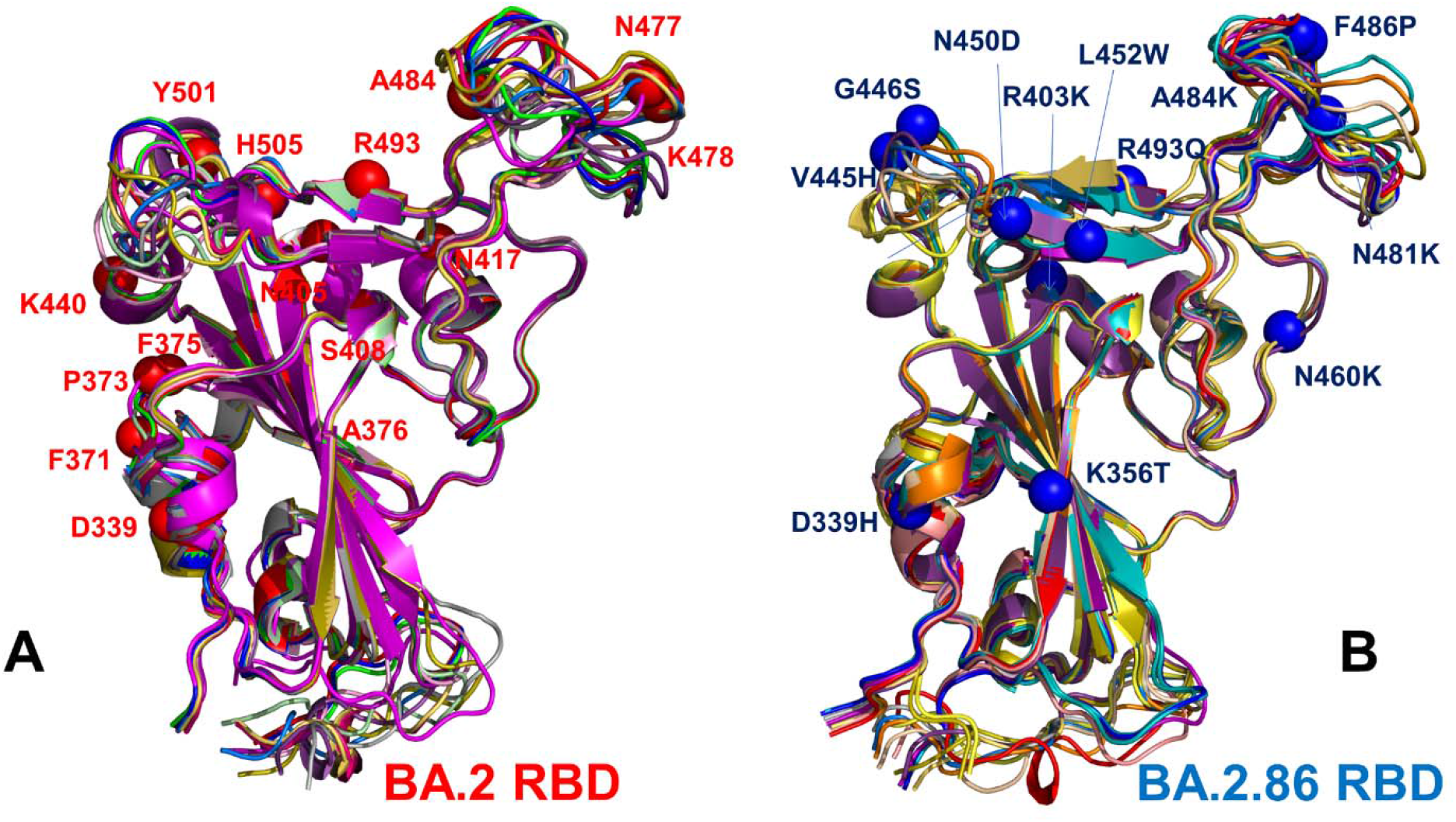
Structural alignment of the AF2-predicted BA.2 and BA.2.86 RBD conformational ensembles. (A) Structural alignment of the AF2-predicted BA.2 conformations with high pLDDT values and the crystal structure of the BA.2 RBD-ACE2 complex. The RBD conformations are shown in ribbons and the position BA.2 mutational sites D339, F371, P373, F375, A376, N405, S408, N417, N477, K478, A484, R493, Y501 and H505 are shown in red spheres. (B) Structural alignment of the AF2-predicted BA.2.86 RBD conformations with high pLDDT values. The sites of unique BA.2.86 mutations D339H, K356T, R403K, V445H, G446S, N450D, L452W, N460K, N481K, A484K, F486P, R493Q are shown in blue spheres.

The important feature of the AF2 predicted conformations for BA.2.86 RBD is that the deletion of V483 does not disrupt the C488-C480 disulfide bond, and this critical stabilization signature is shared in both BA.2 and BA.2.86 structures (Supporting Information, Figure S5). Despite the V483 deletion in the center of the functional RBM loop, which represents a fairly momentous change, and presence of several other mutations N481K, A484K and F486P the AF2-MSA depth ensemble unveiled a tolerant structural response of the RBM loop in BA.2.86 RBD (Figure 5B, Supporting Information, Figure S5A-C). The predicted BA.2 and BA.2.86 conformations have precisely overlapped C480-C488 positions and contacts (Supporting Information, Figure S5C), where N481K and A484K mutational sites in the neighboring positions incur no strain on the position and stability of these disulfide bonds that are critical for RBD folding and stability, thus preserving binding of the BA.2.86 RBD with ACE2. This is consistent with deep mutational scanning (DMS) investigations^120-122^ showing that delV483 only moderately affects RBD stability and ACE2 binding and has a smaller effect than F486S/V mutational changes in Omicron variants.

Importantly, the predicted BA.2.86 RBD ensemble showed small functionally relevant variations in the flexible RBD loops 444-452 and 475-487 (Figure 5B) that harbored unique BA.2.86 mutations V445H, G446S, N450D, L452W, delV483, A484K and F486P representing attractive hotspots of Omicron convergent mutations. These predictions suggested that BA.2.86 mutations would not exert large perturbing effects in the conformational ensemble. Small functional variations of the side chains in the mutated RBM residues indicated while these mutants may have an impact on the stability of the binding interface with ACE2, these substitutions do not perturb the RBD fold and preserve the backbone conformation (Figure 5B). Moreover, variations in the RBD loops 444-452 and 475-487 of the BA.2.86 RBD become more homogeneous than in the BA.2 RBD and represent small loop displacements around the dominant loop state without inducing elements of disorder and preserving ordered “hook-like” folded RBM tip required to minimize binding liabilities of F486P interactions with the ACE2 receptor (Figure 5). These observations echoed the results of MD simulations showing that the RBM loop has an inherent conformational flexibility that is not observed in the static structures of the RBD-ACE2 complexes, where ACE2 and antibody binding to this region may elicit specific distribution of conformations as compared to the unbound RBD form.^123^ The AF2- MSA predictions of the BA.2.86 ensemble also indicated that F486 side chain may experience some fluctuations which is consistent with the presence of many conserved mutations (F486V, F486I, F486S, F486P) seen in other variants making this position a convergent evolutionary hotspot shared by the recent wave of Omicron subvariants. F486 is also one of the major hotspots for escaping neutralization by antibodies. According to the DMS experiments, among the most common F486 mutations (F486V/I/S/L/A/P), F486P imposes the lowest cost in RBD affinity loss and has the largest increase in RBD expression.^120-122^ These predictions may be relevant for the BA.2.86 RBD dynamic responses to binding in which variant-specific mutations may increase adaptability of the RBD loop and exploit the induced plasticity to boost immune evasion. The AF2 predictions are also consistent with our previous MD simulation studies of the Omicron RBD-ACE2 complexes, showing that the major structural differences between Omicron BQ.1 and XBB.1/XBB.1.5 RBD-ACE2 complexes are mostly confined to the fluctuations of the flexible loop regions 444-452 and 475-487^70,71^.

The key finding of this analysis is that AF2 models with the shallow MSA depth are robust in accurately reproducing structural and biophysical experiments of the Omicron RBD-ACE2 complexes. Moreover, we also showed that pLDDT statistical assessment of the AF2 models can be reliably used to identify flexible RBD regions and predict conformational heterogeneity of these adaptable regions with atomistic accuracy.

### Atomistic MD Simulations Reveal Variant-Specific Signatures of Conformational Dynamics and Binding Contacts in the BA.2 and BA.2.86 RBD-ACE2 ACE2 Complexes

We performed comparative all-atom MD simulations of the BA.2 RBD-ACE2 and BA.2.86 RBD-ACE2 complexes. For the latter, the initial structure was taken as the AF2 predicted and refined conformation of the BA..86 RBD-ACE2 complex. Conformational dynamics profiles obtained from MD simulations were similar and revealed several important trends (Figure 6). The RMSF profiles showed local minima regions corresponding to the structured five-stranded antiparallel β-sheet core region that functions as a stable core scaffold (residues 350-360, 375- 380, 394-403) and the interfacial RBD positions involved in the contacts with the ACE2 receptor (residues 490-505 of the binding interface) (Figure 6A). The conformational dynamics profiles revealed marginally greater stability for the BA.2 RBD as compared to BA.2.86 but the differences in thermal fluctuations are relatively minor (Figure 6A). Consistent with AF2 predictions, the profiles displayed displacements in the flexible RBD regions (residues 355-375, 381-394, 444-452, 455-471, 475-487). Of notice are generally larger fluctuations in the BA.2.86 RBD for the flexible region 460-490, particularly the RBM loop featuring mutations N481K, delV483, A484K and F486P (Figure 6B). Despite a moderately increased RBM mobility, the RBM tip is maintained in a stable folded conformation that can be described as “hook-like” folded RBD tip and is similar to the crystallographic conformation of the BA.2 RBD-ACE2 complex. Our previous studies showed that well-ordered and stable “hook-like” conformation of the RBM tip is maintained in BA.2 and XBB.1.5 variants due to hydrophobic interactions provided by F486 (BA.2) and F486P (XBB.1.5).^70-71^

**Figure 6.**
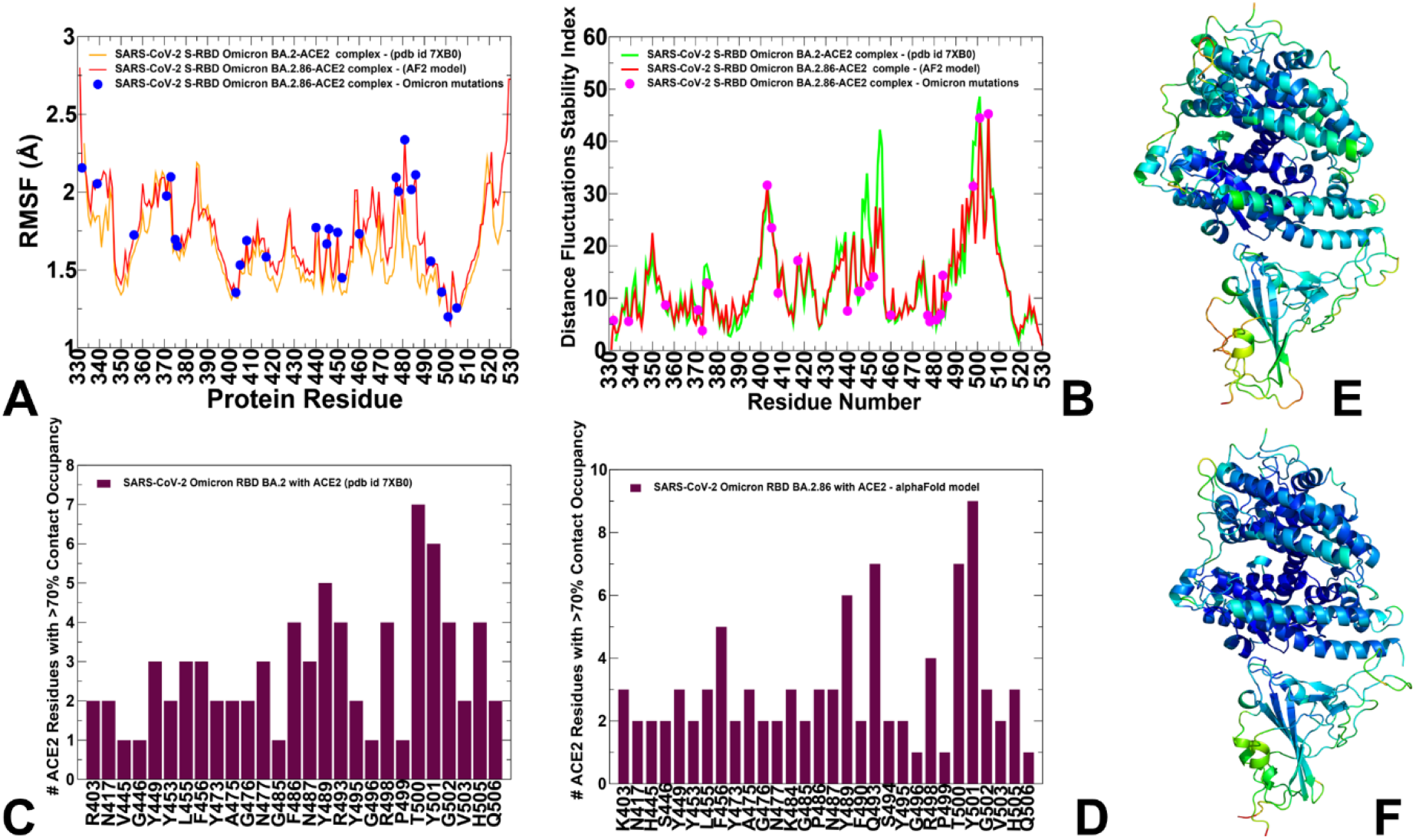
Conformational dynamics profiles obtained MD simulations of the Omicron RBD BA.2 and BA.2.86 RBD complexes with ACE2. (A) The RMSF profiles for the RBD residues obtained from MD simulations of the BA.2 RBD-ACE2 complex, pdb id 7XB0 (in green lines), and BA.2.86 RB-ACE2 complex (in red lines). (B) The distance fluctuation stability index profiles of the BA.2 RBD residues (in green lines) and BA.2.86 RBD residues (in red lines). The positions of Omicron mutational sites are highlighted in magenta-colored filled circles. The ensemble-average numbers of distinct ACE2 residues making stable intermolecular contacts (> 70% occupancy) with the BA.2 RBD residues (C) and BA.2.86 RBD residues (D). Structural mapping of the conformational dynamics profiles for the BA.2 RBD-ACE2 complex (E) and BA.2.86 RBD-ACE2 complex (F) The profiles are mapped onto crystal structure of the BA.2 RBD-ACE2 and AF2-predicted BA.2.86 RBD-ACE2 complex with the rigidity-flexibility sliding scale colored from blue (most rigid) to red (most flexible).

Interestingly, significant BA.2.86 mutational changes in this region, RBM tip showed only marginally elevated mobility preserving the folded hook-like conformation seen in the BA.2 RBD (Figure 6A). The RMSF analysis of the ACE2 residues showed similar profiles across all the examined variants (Supporting Information, Figure S6). The key ACE2 binding motifs correspond to an alpha-helix (residues 24-31) and a beta-sheet (residue 350-356) that display moderate RMSF values in both BA.2 and BA.2.86 complexes. The conformational ensembles of the S-RBD complexes with ACE2 were subjected to distance fluctuations stability analysis based on the dynamic residue correlations (Figure 6B). A comparative analysis of the residue-based distance fluctuation stability indexes revealed several dominant and common peaks, reflecting similarity of the topological and dynamical features of the RBD-ACE2 complexes. Despite a similar shape of the distributions for the BA.2 and BA.2.86 RBD variants, we noticed the reduced stabilit indexes for residues 445-455 in BA.2.86 which implies that this loop becomes somewhat more flexible in BA.2.86 owing to modifications V445H, G446S, N450D and L452W (Figure 6B). Instuctively, AF2-predicted conformations similarly revealed the variability of this RBD loop, showing that AF2-generated ensemble and model ranking using pLDDT metric can capture the dynamics signatures of the RBD-ACE2 complexes.

Using conformational ensembles of the Omicron RBD-ACE2 complexes, we performed a statistical analysis of the intermolecular contacts that revealed several fundamental commonalities and differences in the interaction profiles for the Omicron RBD subvariants (Figure 6C,D). The RBD-ACE2 contacts with the occupancy > 70% in the MD trajectories were considered as long-lived stable interactions and were recorded for this analysis. The RBD residues can be classified in several groups based on their conservation and respective roles in binding with the host receptor. One of these groups includes identical residues such as Y453, N487, Y489, T500, and G502 and homologous positions (Y449/F/H, F456/L, Y473/F, F486/L, and Y505H), while other group includes more diverse residues undergoing various modifications such as G446/S/T, L455/S/Y, A475/P/S, G476/D, G496/S, K417/V/N/R/T, E484/K/P/Q/V/A, Q493/N/E/R/Y, Q498/Y/H/R, and N501/Y/T/D/S. The conserved residues from the first group make consistent and similar interactions in all Omicron variants, indicating that these RBD positions can function as molecular determinants of the RBD stability and binding affinity (Figures 6C,D). The distribution of contacts in the BA.2 RBD-ACE2 complex indicated that stable interactions are established across the entire binding interface, particularly revealing the formation of salt bridges by R403 with E37, R493 with E35 and D38 (Figure 6C). A considerable number of ACE2 residues participate in interactions with F486, Y489, R493, R498, T500 and Y501 positions. In particular, R498 interacts with D38, Y41, Q42 and L45 ACE2 residues; T500 forms stable interactions with Y41, L45, G326, Q330, K353, G354, D355 and R357; and Y501 is engaged in the favorable stable contacts with D38, Y41, K353, G354 and D355 (Figure 6C). The analysis showed that the interfacial salt bridge interactions in the BA.2 RBD formed by R493 with E37, E35 and D38 residues in ACE2 are partially lost in the BA.2.86 RBD-ACE2 complex. To compensate for this loss the mutated R493Q forms a number of stable interfacial contacts with D30, K31, N33, H34 and E35 including hydrogen-bonding interaction with the K31 side chain and the carboxyl group of E35 from ACE2 (Figure 6D). Interestingly, the salt bridge formed by R493 of BA.2 with E35 of ACE2 is replaced by Q493 in BA.2.75, BF.7, XBB.1 and BA.2.86 in a manner reminiscent of the Wu-Hu-1 strain. Hence, diverse mutations in different Omicron variants including BA.2.86 can reassemble the interaction network while preserving a strikingly convergent binding pattern of interactions and major determinants of binding.

### Comparative Mutational Scanning of the BA.2 and BA.2.86 RBD Residues Identifies Universal and Variant-Specific Structural Stability and Binding Affinity Hotspots

Using conformational ensembles obtained from AF2 predictions and MD simulations we performed a systematic mutational scanning of the BA.2 and BA.2.86 RBD residues in the RBD-ACE2 complexes (Figure 7). In silico mutational scanning was done using BeAtMuSiC approach^111-113^ by averaging the binding free energy changes over the equilibrium ensembles and allows for predictions of the mutation-induced changes of the binding interactions and the stability of the complex. The resulting mutational scanning heatmaps are reported for the RBD binding interface residues that make stable contacts with ACE2 in the course of simulations. To provide a systematic comparison, we constructed mutational heatmaps for the RBD interface residues of BA.2 RBD-ACE2 (Figure 7A) and BA.2.86 RBD-ACE2 complexes (Figure 7B). Consistent with DMS experiments of SARS-CoV-2 S VOC’s^120-122^ the hydrophobic residues Y453, L455, F456, F486, Y489 and Y501 play a decisive role in stability and binding for both BA.2 (Figure 7A) and BA.2.86 complexes (Figure 7B). The large destabilization changes become more particularly pronounced for Y453, L455 and F456 while also revealing high sensitivity of F486, N487, Y489, R493, T500, Y501, and H505 residues (Figure 7). Mutational heatmaps clearly showed that all substitutions in these key interfacial positions can incur a consistent and considerable loss in the stability and binding affinity with ACE2. In addition, mutational scanning of the RBD residues F486, N487 and H505 showed appreciable and consistent destabilization changes, placing these residues as another important group of energetic centers (Figure 7). This analysis is consistent with previous studies, suggesting that these conserved hydrophobic RBD residues may be universally important for binding across all Omicron variants and act as stabilizing sites of the RBD stability and binding affinity.^70,71^

**Figure 7.**
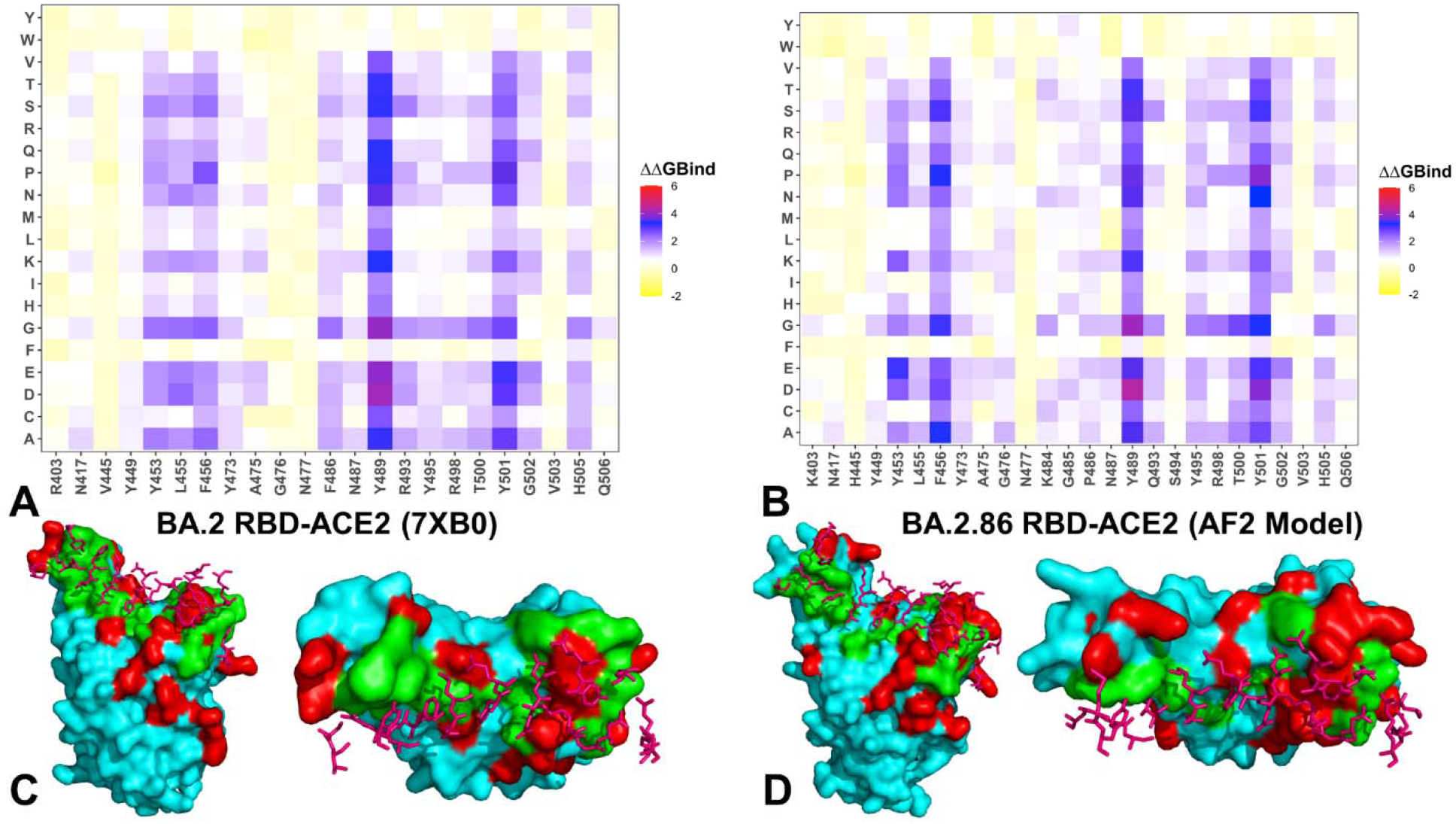
Ensemble-based dynamic mutational profiling of the RBD intermolecular interfaces in the Omicron RBD-ACE2 complexes. The mutational scanning heatmaps are shown for the interfacial RBD residues in the BA.2 RBD-ACE2 complex (A) and BA.2.86 RBD-ACE2 complex (B). Structural mapping of the RBD binding epitopes of the BA.2 RBD-ACE2 complex (C) and BA.2.86 RBD-ACE2 complex (D). The RBD binding epitope is in green-colored surface. The Omicron RBD mutational sites are shown in red surface. The ACE2 binding residues are in pink sticks. The standard errors of the mean for binding free energy changes and are within ∼ 0.05-0.12 kcal/mol using averages based on a total of 1,000 samples obtained from the three MD trajectories for each system.

The mutational heatmap for the BA.2.86 RBD-ACE2 was quite similar showing the major hotspots for Y453, L455, F456, Y489, Y501 and H505 positions that are shared between BA.2 and BA.2.86 (Figure 7B). The direct binding interface is broader for BA.2.86 revealing small variations and tolerance to mutational changes in positions Y473, A475, G476, N477, K484, G485, P486 (Figure 7B). Notably, unique BA.2.86 mutational positions N450D and L452W are distant from the immediate intermolecular interface and do not form direct contacts with the ACE2 receptor. To quantify the stability and binding free energy changes for the sites of BA.2.86 mutations, we reported detailed mutational scanning results for these positions (Figure 8). The results showed that binding free energy changes induced by scanning in the BA.2.86 positions H339 (Figure 8A), T356 (Figure 8B), H445 (Figure 8D), S446 (Figure 8E), D450 (Figure 8F), W452 (Figure 8G), K460 (Figure 8H), and K484 (Figure 8J) are small and mostly destabilizing with ΔΔG < 1.0 kcal/mol. Noticeably, mutational scanning of K481 position resulted in minor stabilizing changes reflecting favorable stability effects as K481 is not involved in direct intermolecular contacts with the ACE2 receptor (Figure 8I). The initial analysis of the BA.2.86 antigenicity and binding suggested that he higher receptor binding affinity of the BA.2.86 might be partly attributed to the additional positive charges associated with V445H, N460K, N481K and A484K sites.^51^ However, our AF2-enabled analysis of the RBD-ACE2 contacts derived from simulations of BA.2.86 RBD-ACE2 complex indicated that mutational sites V445H, G446S, N450D, L452W, A484K are not involved in formation of long-lived specific interactions (Figure 6D). Mutational scanning confirmed that these positions displayed a considerable degree of tolerance to modifications and are unlikely to significantly contribute to the binding affinity. These results are consistent with the biophysical analysis of the BA.2.86 mutations^51^ suggesting that D339H, K356T, V445H, G446S, N450D, L452W, N460K, N481K and A484K may have emerged primarily to strengthen immune evasion and mediate the enhanced resistance which implies that infectivity may be traded for higher immune evasion during long-term host-viral evolution.^51-53^

**Figure 8.**
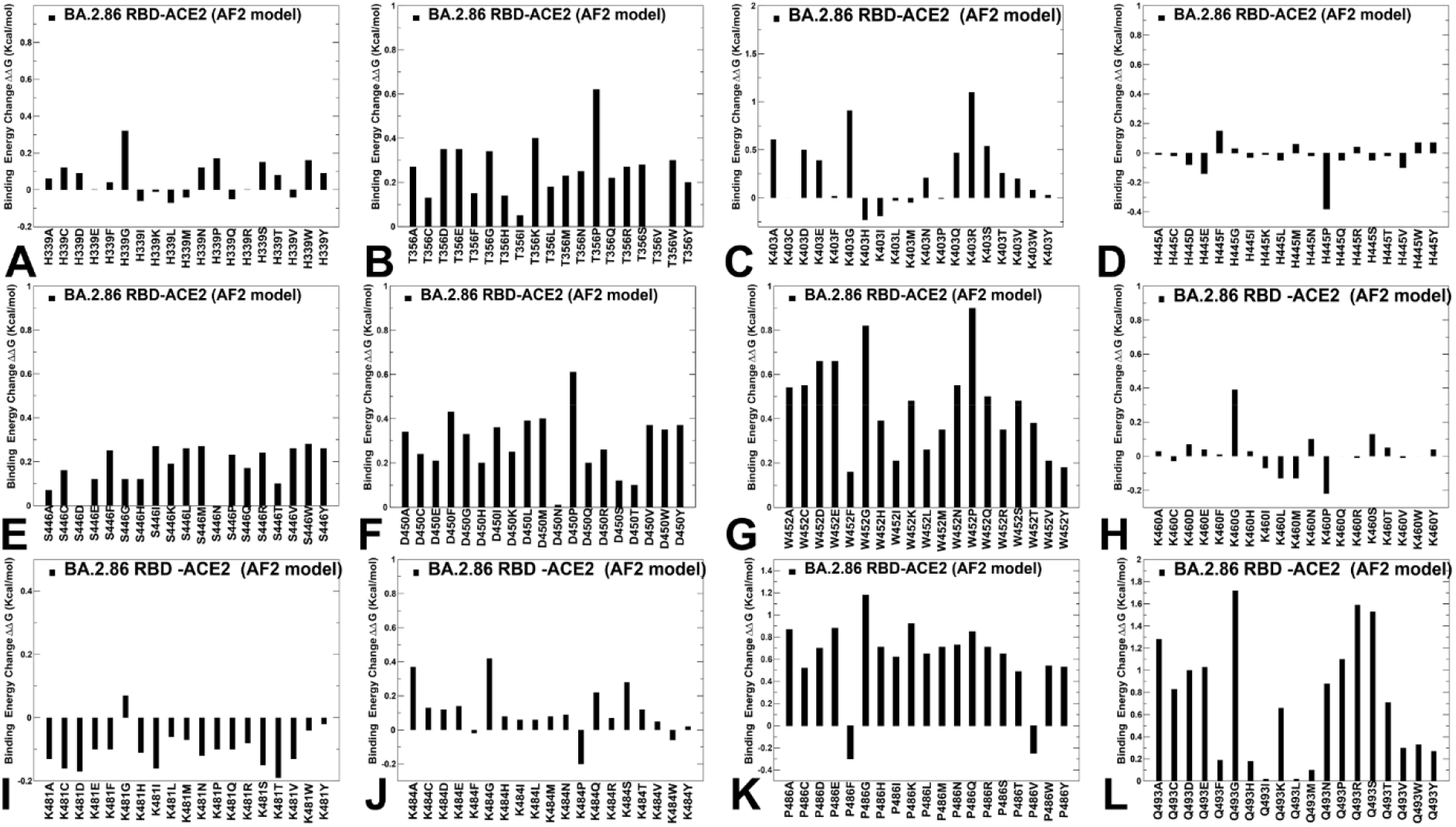
Ensemble-based mutational scanning of stability and binding for the individual BA.2.86 mutational sites in the BA.2.86 background. The profiles of computed binding free energy changes ΔΔG upon 19 single substitutions for the Omicron BA.2.86 mutational sites D339H (A), K356T (B), R403K (C) V445H (D), G446S (E), N450D (F), L452W (G), N460K (H), N481K (I), A484K (J), F486P (K), and R493Q (L). The respective binding free energy changes are shown in maroon-colored filled bars. The positive binding free energy values ΔΔG correspond to destabilizing changes and negative binding free energy changes are associated with stabilizing changes.

Of particular interest is the mutational scanning profile of R403K position where the reversed change K403R is highly unfavorable (ΔΔG = 1.2 kcal/mol) suggesting that R403K in BA.2.86 can lead to the increased binding affinity (Figure 8C). These results are in agreement with the recent experimental data which examined mutations in the BA.2.86 RBD by generating a set of reverse mutations and demonstrated that K403R reversal significantly increased the equilibrium dissociation constant signaling the reduced binding affinity upon reversed mutation.^56^ Our mutational scanning results are also consistent with these biochemical experiments suggesting that R403K mutation in BA.2.86 may contribute significantly to the binding affinity with the ACE2. Of special significance is the pattern of energetic changes in BA.2.86 positions P486 (Figure 8K) and Q493 (Figure 8L) that correspond to the critical RBD positions involved in both binding and immune escape. We observed that modifications of P486 with the exception of P486F and P486V are destabilizing, confirming the notion that F486P mutation may have evolved to secure productive binding at minimum loss while allowing a sufficient room for immune escape from antibodies targeting this region.^120-122^ F486 from the RBM loop is a critical site of convergent evolution is that is exploited by BA.2, BA.2.75, BF.7, XBB.1, XBB.1.5 and BA.2.86 variants as one of the hotspots of compensatory binding and immune evasion. Indeed, F486 in BA.2 and BA.2.75 is substituted with F486V in BF.7 and F486S in XBB.1 and F486P in XBB.1.5 and BA.2.86 variants.^124-126^

Our mutational scanning analysis is consistent with these studies suggesting that mutations in these hotspots allow for delicate manipulation of conflicting fitness requirements to properly balance ACE2 binding affinity and immune evasion. Another critical site of convergent evolution is reversed R493Q mutation in BA.2.86 variant. Mutational scanning results showed that modifications at Q493 positions are generally destabilizing, particularly the back-reversed Q493R modification in the background of BA.2.86 (Figure 8L). This suggested that R493Q may improve the binding affinity and compensate for partial binding loss incurred by F486P mutation. These findings are also in agreement with previous studies showing that R493Q reversal may induce the increased affinity of the RBD with ACE2 receptor.^127^

To further investigate and compare the impact of mutations on the binding affinity to ACE2 receptor, we computed the binding free energy changes that unique BA.2.86 mutations induce in the BA.2 RBD-ACE2 complex (Figure 9A) as well as the effect of the reversed mutations in the BA.2.86 RBD structure (Figure 9B). A particular objective of this comparison was to estimate the role of the BA.2 and BA.2.86 backgrounds on the mutational effects which may indicate synergistic epistatic effects of the BA.2.86 substitutions. It appeared that D339H, K356T, V445H, G446S, N450D, L452W, N481K and A484K substitutions cause small changes (ΔΔG < 0.5 kcal/mol) in the BA.2 structure, indicating that these substitutions play no significant role in modulating binding affinity (Figure 9A). Notably, A484K and R493Q mutations can induce more appreciable but still moderate stabilization changes that improve the binding affinity in the BA.2 RBD-ACE2 complex (Figure 9A). The assessment of binding free energies upon reversal mutations in the BA.2.86 RBD-ACE2 complex (the BA.2.86 background) not only reasserted the important contribution of K403 and Q493 to the ACE2 binding but also yielded significantly larger destabilization changes for reversed K403R (ΔΔG = 1.2 kcal/mol) and Q493R changes (ΔΔG = 1.6 kcal/mol) (Figure 9B). Hence, in the BA.2.86 background K403 and Q493 positions are far more significant for the ACE2 bindi. i.e., suggesting a potential epistatic effect of R403K and R493Q mutations in the BA.2.86 RBD-ACE2 complex. These results echo similar arguments proposed in the recent functional studies of binding and antigenicity of the BA.2.86 variant.^56^ Our findings are particularly intriguing in light of the recent studies showing that convergent evolution of the Omicron XBB lineages due to flip changes L455F and F456L between adjacent residues on the ACE2 binding interface can synergistically enhances antibody evasion and ACE2 binding suggesting epistatic effect of these mutations.^128^ Moreover, this study showed that the L455F/F456L flip mediates conformation changes of the binding interface that amplifies binding contribution of Q493 position which suggested a synergistic epistasis between these functional hotspots of the RBD-ACE2 binding.^128^ The presented data on forward and reversed mutational scanning of the BA.2.86 mutations in the BA.2 and BA.2.86 backgrounds suggested a potential epistatic effect of R493Q in the context of the BA.2.86 variant.

**Figure 9.**
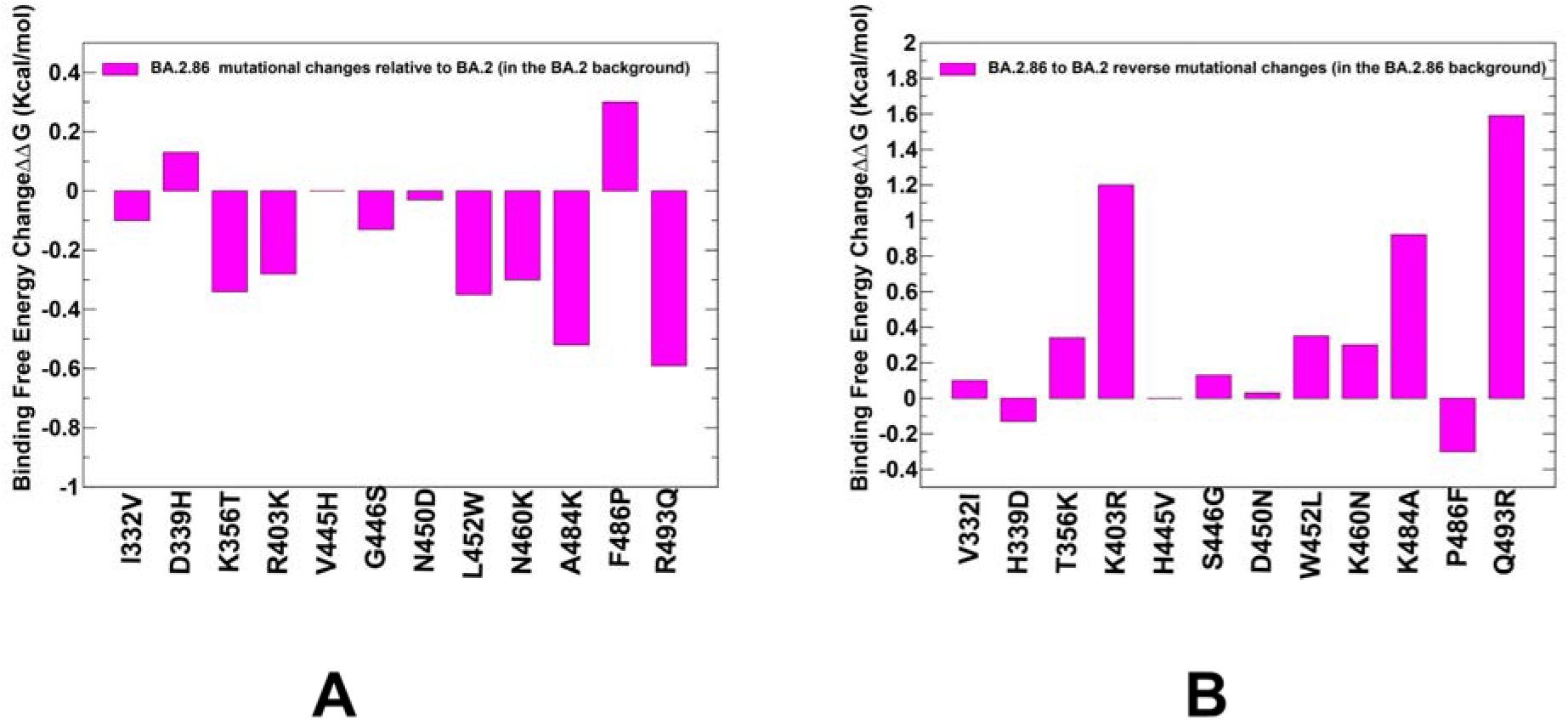
The computed binding free energy changes that unique BA.2.86 mutations induce in the BA.2 background in the BA.2 RBD-ACE2 complex (A) and the effect of the reversed to BA.2 mutations induce in the BA.2.86 background in the BA.2.86 RBD-ACE2 complex (B). The computed binding free energies are shown in magenta-colored filled bars. The positive binding free energy values ΔΔG correspond to destabilizing changes and negative binding free energy changes are associated with stabilizing changes.

Functional studies showed that Q493R markedly diminished ACE2 affinity while BA.2 with a single R493Q change improved binding affinity to the ACE2 receptor^34^ sensitized BA.2 to neutralization by several class 1 and 2 RBD monoclonal antibodies.^44^ According to our results, the main drivers of the binding free energy changes in BA.2.86 would result from balancing “dance” performed by hotspots of convergent evolution where a moderate ACE2 binding loss incurred by F486P mutation is compensated by the favorable binding effect of the R403K and R493Q mutations. Similar balancing effects were observed in Omicron BA.4/BA.5 variant where the R493Q reversion in the BA.4/5 contributed to evading immunity and improvements in the ACE2 binding affinity. ^44^ We argue that specific modulation of the BA.2.86 RBD binding affinity may be determined by a combined effect of R493Q and F486P mutations that represent key attractive hotspots of convergent evolution. These results are consistent with the notion that most of the BA.2.86 mutations may have primarily emerged to improve the evasion of acquired immunity by Omicron variants rather than affecting the ACE2 binding. These findings lend support the evolutionary mechanism according to which emerging RBD mutations in the BA.2.86 variant (particularly N440K, N460K, T478K, N481K, A484K, and F486P) may be primarily driven by the immune pressure-driven changes while the improved ACE2 binding affinity relative to BA.2 is mediated via R403K and R493Q mutations.^56^ In general, the results indicated that latest Omicron subvariants including BA.2.86 may evolve to accumulate convergent immune escape mutations while exploiting potential synergistic epistatic effects of selected group of hotspots R403K, R493Q as well as L455F/F456L to enable sufficient ACE2-binding capability. Several lines of evidence indicated that the observed coordination of evolution at different sites is due to epistatic, rather than random selection of mutations.^129,130^

Using the conformational equilibrium ensembles obtained from MD simulations, we computed the binding free energies for the Omicron RBD BA.2 and BA.2.86 RBD complexes with ACE2 using the MM-GBSA method^109,110^ A common strategy to reduce noise and cancel errors in MM-GBSA computations is to run MD simulations on the complex only, with snapshots taken from a single trajectory to calculate each free energy component. The total binding free energy changes showed a moderately more favorable binding affinity for the BA.2.86 RBD (Supporting Information, Table S1, Figure S7). The breakdown of the MM-GBSA binding energies showed that the electrostatic contribution is more favorable for the BA.2.86 RBD-ACE2 complex but is offset by the unfavorable solvation (Table S1). This reflected the presence of BA.2.86 mutational sites N460K, N481K, A484K that increased the electrostatic interactions but the net effect of these mutational sites to binding is minor due to balance of electrostatic and solvation contributions (Table S1). The experimentally measured ACE2 binding affinities for two versions of the BA.2.86 S proteins with K_D_ values of 0.54 nM and 0.60 nM, while compared to the K_D_ value of the BA.2 S protein of 1.68 nM.^51^ As expected, the introduction of R403K, N460K, N481K, and A484K mutations enhanced the contribution of electrostatic interactions (Supporting Information, Figure S7) but for N460K, N481K, an A484K these favorable interactions are largely offset by unfavorable solvation contributions resulting in the net moderately destabilizing interactions.

Of particular interest is the analysis of R403K, R493Q and F486P mutational effects in the BA.2.86 RBD-ACE2 complex (Supporting Information, Figure S7D-E). We found that the binding energy contribution for R403K is more favorable (ΔG = -3.46 kcal/mol) while in the BA.2 RBD-ACE2 complex this contribution ΔG = -1.03 kcal/mol which is due to a more favorable balance of electrostatics and solvation energies in the BA.2.86 complex (Supporting Information, Figure S7D-E). Importantly, MM-GBSA analysis showed that the R493Q mutation in BA.2.86 yielded ΔG = -3.86 kcal/mol while the respective contribution of R493 in BA.2 accounted for ΔG = -2.03 kcal/mol thus confirming a stronger favorable effect of reversal R493Q on binding affinity in the BA.2.86 variant (Supporting Information, Figure S7D-E). It appeared that R493 in BA.2 can enhance the electrostatic interactions, but this contribution is offset entirely by the unfavorable solvation, while R493Q in BA.2.86 produced a more favorably balanced total effect of favorable van der Waals and electrostatic interactions. Notably, the MM-GBSA decomposition analysis also confirmed a highly favorable contribution of F486 in BA.2 to the binding energy (ΔG = -4.87 kcal/mol with a dominant van der Waals component of -5.24 kcal/mol), while F486P in BA.2.86 yielded a significantly reduced binding contribution (ΔG = -1.66 kcal/mol) primarily due to a considerable loss of the packing van der Waals contacts (Supporting Information, Figure S7D-E). These findings provided a useful insight into the mechanism underlying stronger binding of the BA.2.86 variant relative to BA.2 suggesting that differences in the binding affinity are determined by cumulatively better balance of electrostatic and solvation contributions for several binding affinity hotspots including R403K, F486P, and R493Q, while universally important for binding Y489, R498 and Y501 maintain their favorable interactions in both complexes (Supporting Information, Figure S7).

The analysis of the BA.2.86 antigenicity and binding suggested that he higher receptor binding affinity of the BA.2.86 was initially proposed to result from the additional electrostatic interactions associated with V445H, N460K, N481K and A484K sites.^51^ At the same time, mutational scanning suggested only small contributions to the binding affinity. To further investigate this conjecture, we computed and compared the electrostatic potential for the BA.2 and BA.2.86 RBD-ACE2 complexes (Figure 10). The electrostatic potentials on the RBD surfaces for the Omicron BA.2 (Figure 10A,B) and BA.2.86 (Figure 10C,D) complexes are positive with variable charge distributions, showing relatively moderate changes in the overall surface distribution between subvariants. However, we noticed a stronger positively charged potential in the Omicron RBD BA.2.86 (Figure 10C,D) showing positive densities in the RBM-interacting regions and distal from binding RBD regions.

**Figure 10.**
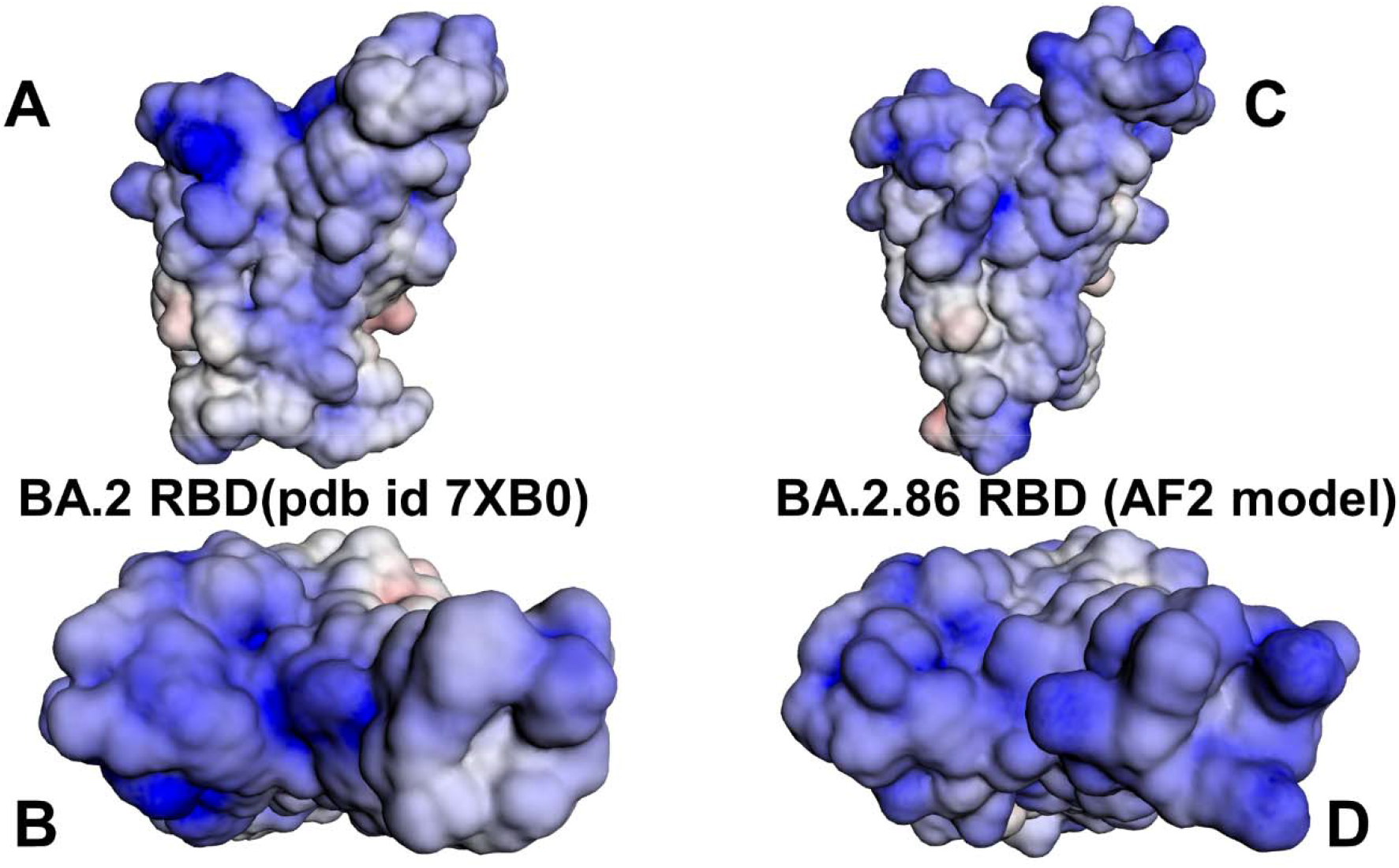
The distribution of the electrostatic potentials calculated on the molecular surface of the BA.2 RBD-ACE2 complex (A,B) and BA.2.86 RBD-ACE2 (C,D). The crystal structure of the BA.2 RBD-ACE2 complex and the best AF2-predicted model of the BA.2.86 RBD-ACE2 complex are used in computations of the electrostatic potentials. The color scale of the electrostatic potential surface is in units of kT/e at T = 37°C. Electro-positively and electronegatively charged areas are colored in blue and red, respectively. Neutral residues are shown in white.

Hence, computations of the electrostatic potential surfaces confirmed that the BA.2.86 RBD featured a stronger positive electrostatic potential due to substitutions to basic residues N440K, N460K, T478K, N481K, A484K, and Q498R. To facilitate a comparison, we also compared the electrostatic potential using the experimental structures of BA.1, BA.2, BA.2.75 BA.3, BA.4/BA.5 and XBB.1.5 RBD complexes with ACE2 (Supporting Information, Figure S8) revealing that BA.2.86 displayed the strongest positive potential in the binding interface and RBM regions. These results are consistent with recent studies showing that the evolution of the electrostatic surface in Omicron variants shows a gradual accumulation of positive surface charges in the RBD over time and that related clades have similar electrostatic surfaces.^72,73^ It was also noted that many antibodies have positively charged S RBD-recognition surfaces.^73^ Based on the mutational scanning data and electrostatic potential analysis, we suggested that evolution of the electrostatic RBD surface in latest Omicron variants, including further strengthening of the positive electrostatic potential on the BA.2.86 RBD, may have been primarily exploited to evade antibodies without compromising binding affinity with ACE2. We argue that BA.2.86 lineage may have evolved to outcompete other Omicron subvariants by boosting immune suppression while balancing binding affinity with ACE2 via through compensatory effect of R493Q and F486P mutations.

### Mutational Profiling of Protein Binding Interfaces with Distinct Classes of Antibodies: Quantifying Functional Role of BA.2.86 Mutations in Immune Evasion

We embarked on structure-based mutational analysis of the S protein binding with different classes of RBD-targeted antibodies, focusing specifically on the role of BA.2.86 mutations in mediating potential resistance to broad class of antibodies and eliciting robust immune escape. We specifically examined a panel of monoclonal antibodies that were reported to retain activity against BA.2. XBB.1.5 and EG.5.1 but displayed a markedly reduced or completely abolished neutralization potential against BA.2.86 variant.^51^ These biochemical studies highlighted the ability of the BA.2.86 variant to effectively evade the immune response triggered by a panel of antibodies targeting multiple epitopes on the S protein. To understand the antibody evasion properties of BA.2.86 in greater atomistic detail and clarify role of the BA.2.86 mutations in immune escape against distinct classes of RBD-targeted antibodies, we performed large scale structure-based mutational scanning of the S protein binding interfaces with a panel of these RBD antibodies featuring three distinct antibody classes targeting different RBD epitopes.^51^ Mutational profiling analysis allowed for direct comparison with the latest experiments that reported fold changes in IC_50_ of antibody binding with BA.2.86 relative to BA.2 variant^51^ and enabled to clarify specific role of unique BA.2.86 mutations in mediating antibody resistance (Figures 11-13).

**Figure 11.**
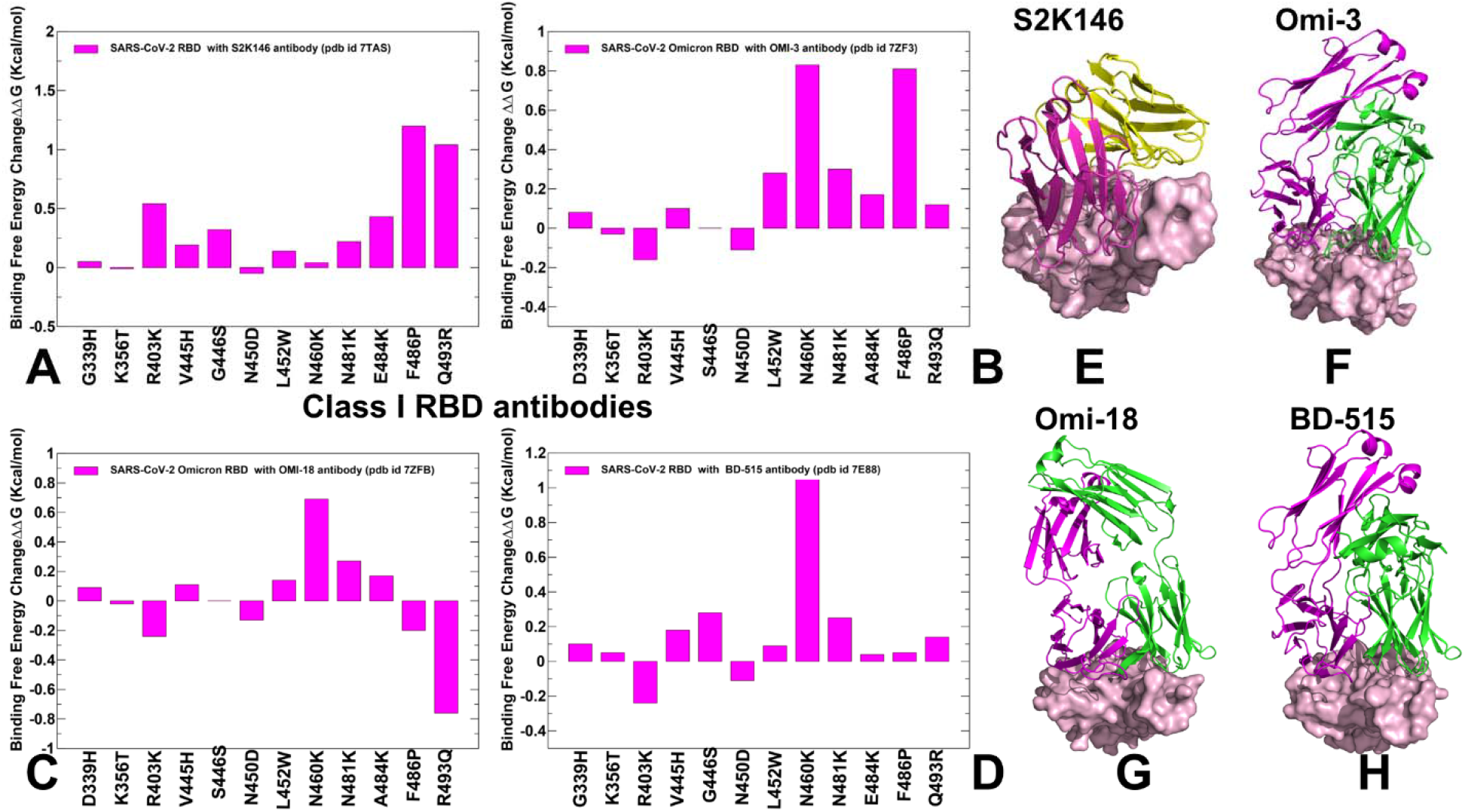
Structure-based mutational profiling of the S-RBD complexes with class 1 of RBD antibodies. The mutational screening evaluates binding energy changes induced by BA.2.86 mutations in the RBD-antibody complexes. Mutational profiling of the S-RBD complex with S2K146 (A), S-RBD Omicron complex with Omi-3 (B), S-RBD Omicron complex with Omi-18 (C), and S-RBD in complex with BD-515 (D). The binding free energy changes are shown in magenta-colored filled bars. The positive binding free energy values ΔΔG correspond to destabilizing changes and negative binding free energy changes are associated with stabilizing changes. The 3D structures of the RBD-antibody complexes are shown for RBD-S2K146 (E),

The examined structures for class1 RBD antibodies included S-RBD complex with S2K146 (pdb id 7TAS)^131^, S-RBD Omicron complex with Omi-3 (pdb id 7ZF3)^132^, S-RBD Omicron complex with Omi-18 (pdb id 7ZFB)^132^, S beta trimer complex with Omi-42 (pdb id 7ZR7)^132^ and S-RBD in complex with BD-515 (pdb id 7E88)^133^. In the analysis of mutational scanning, we specifically focused on the S-antibody binding energy changes induced by BA.2.86 mutations in the corresponding RBD positions (Figure 11). The binding free energy changes associated with BA.2.86 mutations in the complex with S2K146 (Figure 11A) showed an appreciable loss of binding upon F486P and Q493R mutations which implied that F486P in the BA.2.86 is highly deleterious for S2K146 binding while the reversed R493Q in BA.2.86 is favorable for the antibody binding. RBD-Omi-3 (F), RBD-Omi-18 (G), and RBD- BD-515 complexes (H). The RBD is shown in pink-colored surface representation and the antibodies are shown in ribbons (heavy chain in magenta and light chain in green-colored ribbons).

These observations agree with the experimental screening showing that F486P caused dramatic fold change (> 254) in IC_50_ of antibody binding with BA.2.86 relative to BA.2 variant.^51^ At the same time, according to these experiments^51^, R493Q mutation may lead to ∼ 11 fold favorable fold change in the antibody binding. For Omi-3 antibody, the two experimental hotspots that cause significant loss of binding with BA.2.86 are N460K and F486P mutations^51^ which is accurately captured in the mutational scanning revealing destabilizing ΔΔG = 0.62 kcal/mol for N460K mutation and ΔΔG = 0.81 kcal/mol for F486P (Figure 11B. Notably, our results also reproduced the experimentally observed R403K-induced favorable change in binding and moderate loss in the Omi-18 affinity caused by R493Q reversal (Figure 11B).

A similar pattern was seen in mutational scanning of the Omicron complex with Omi-18 antibody, revealing the same two energetic hotspots N460K and F486P that cause a significant reduction in the binding affinity (Figure 11C). In this case, both R403K and R493Q mutations appeared to be favorable for the antibody binding but the net effect may be offset by loss in affinity caused by N460K and F486P mutations. In agreement with the biochemical studies^51^ mutational profiling of the S-RBD with another class I RBD antibody BD-515 unveiled a single binding hotspot N460K that can induce a significant loss in binding, while both R403K and R493Q changes can be marginally favorable for the antibody binding (Figure 11D). These results showed that mutations N460K and F486P, also shared by XBB.1.5 and EG.5.1 variants, can mediate resistance to some RBD class 1 antibodies by disrupting key hydrogen bonding between the RBD and antibodies in case of N460K and reducing the hydrophobic packing with the ACE2-mimicking antibodies. Accordingly, these mutations can modulate the balance between ACE2 binding and antibody escape. Interestingly, R403K and R493Q mutations that are important drivers of the BA.2.86 binding with the ACE2 are also favorable for antibody binding, while reversed Q493R is moderately unfavorable (Figure 11). We argue that BA.2.86 variant may have accumulated other mutations to potentially balance a synergistic stabilizing effect of R403K/R493Q mutations on both ACE2 binding and antibody interactions.

We also examined the class 2 RBD antibodies including S-RBD complex with COV2-2196 (tixagevimab) (pdb id 8D8Q)^134^, Omicron S-RBD with XGV347 (pdb id 7WED)^135^, Omicron S-RBD with XGV051 (pdb id 7WTG)^136^ and Omicron S-RBD complex with ZCB11 (pdb id 7XH8).^137^ Consistently, mutational scanning of the BA.2.86 mutational sites demonstrated that F486 is the key binding hotspot as F486P modification resulted in significant binding affinity loss for this class of antibodies (Figure 12). These findings are in excellent agreement with the biochemical experiments^51^ revealing dramatic losses in antibody binding relative to BA.2 (> 81 for COV2-2196, > 7505 for XGV347, ∼ 298 for XGV051 and > 2293 for ZCB11) that are induced by F486P mutation. Structural studies showed that formation of a hydrophobic cage F486 by COV2-2196 is a primary driver of binding and F486A mutation can completely abrogate binding with S protein (Figure 12A).^134^ Consistent with these experiments, mutational profiling singled out F486P mutation as the dominant hotspot of antibody resistance (Figure 12A). The antibodies XGV347 and XGV051 form extensive hydrophobic interactions contributed by Y449, Y453, L455, A475, A484, F486 and Y489 positions from Omicron RBD (Figure 12B,C).^135,136^ For both XGV347 and XGV051 mutational profiling revealed the same trend showing that BA.2.86 mutation F486P induced a large loss in the antibody binding affinity (ΔΔG > 3.0 kcal/mol) and confirming a critical role of this convergent evolutionary hotspot in immune escape. Notably, the magnitude of the antibody affinity loss due to F486P mutation significantly exceeded a moderate detrimental effect on ACE2 binding.

**Figure 12.**
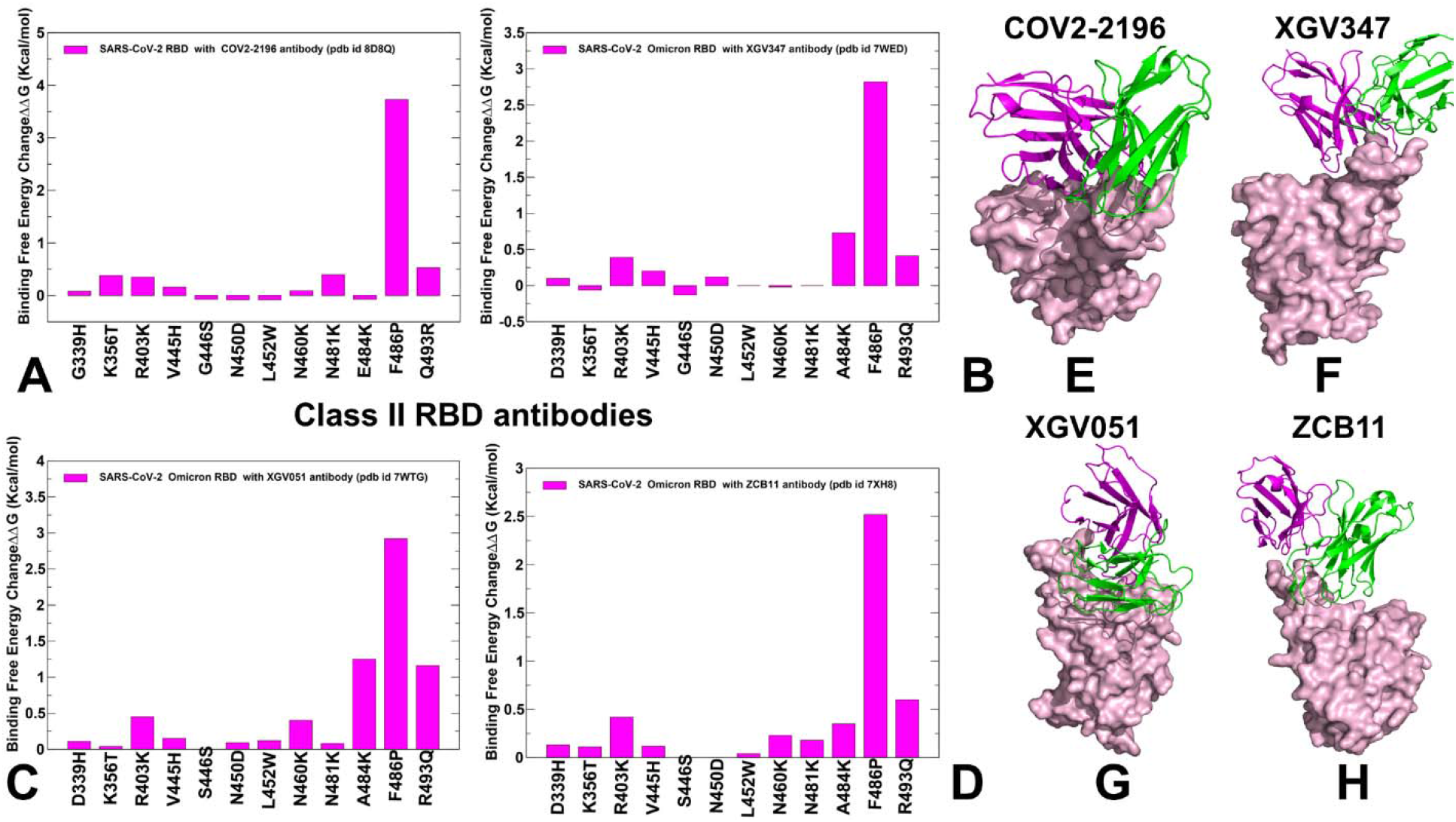
Structure-based mutational profiling of the S-RBD complexes with class 2 of RBD antibodies. The mutational screening evaluates binding energy changes induced by BA.2.86 mutations in the corresponding RBD positions of the RBD-antibody complexes. Mutational profiling of the S-RBD complex with COV2-2196 (A), S-RBD Omicron complex with XGV347 (B), S-RBD Omicron complex with XGV051 (C), and S-RBD in complex with ZCB11 (D). The respective binding free energy changes are shown in magenta-colored filled bars. The 3D structures are shown for the S-RBD complex with COV2-2196 (E), S-RBD Omicron complex with XGV347 (F), S-RBD Omicron complex with XGV051 (G), and S-RBD in complex with ZCB11 (H). The RBD is shown in pink-colored surface representation and the antibodies are shown in ribbons (heavy chain in magenta and light chain in green-colored ribbons).

**Figure 13.**
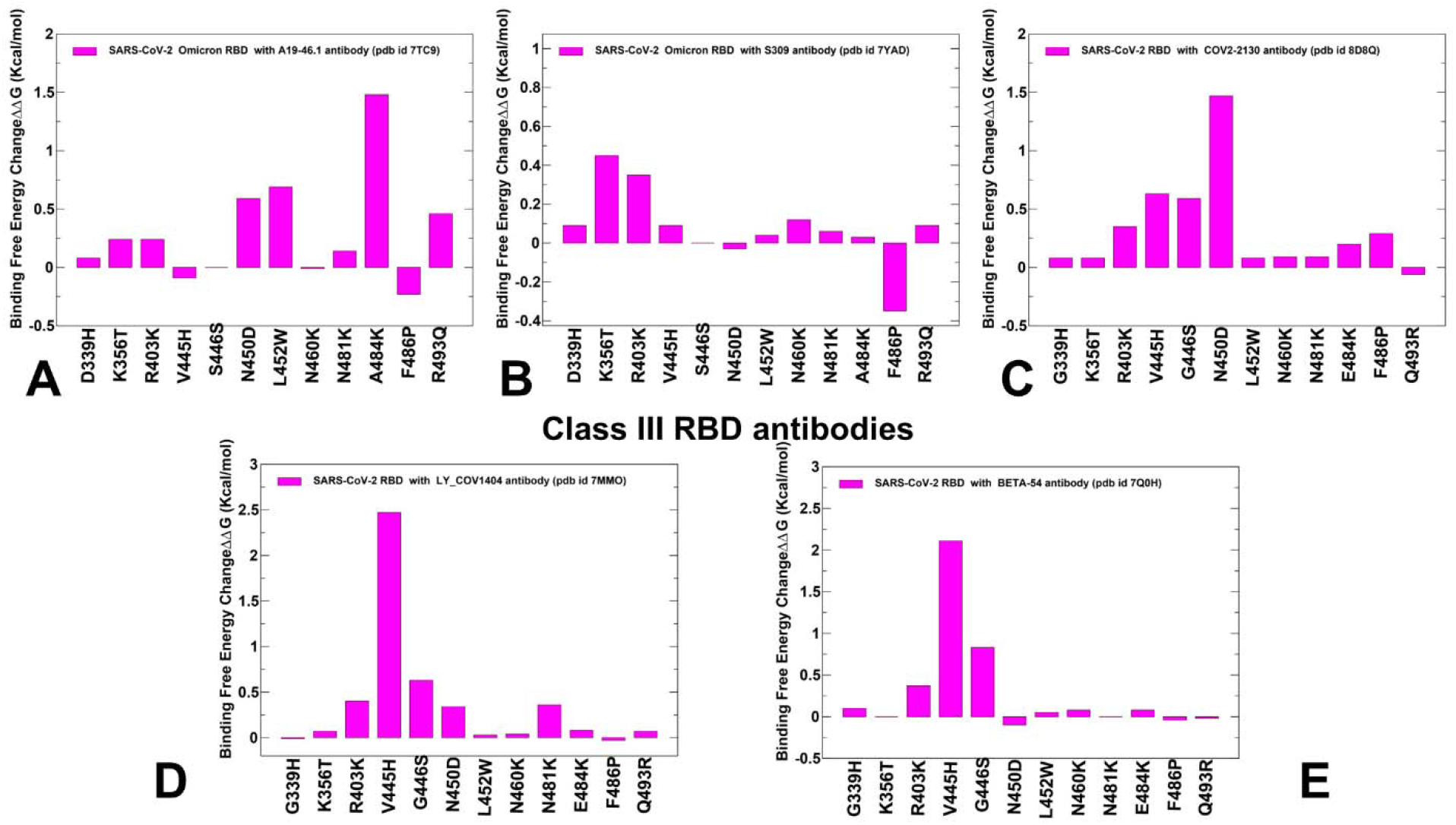
Structure-based mutational profiling of the S-RBD complexes with class 3 of RBD antibodies. The mutational screening evaluates binding energy changes induced by BA.2.86 mutations in the corresponding RBD positions of the RBD-antibody complexes. Mutational profiling of the S-RBD complex with A19-46.1 (A), Omicron S-RBD with S309 (B), S-RBD with COV2-2130 (C), S-RBD with LY-CoV1404 (D) and S-RBD beta variant complex with Beta-54 (E). The binding free energy changes are shown in magenta-colored filled bars.

By quantifying these differences, our results supported the notion that F486P mutation in BA.2.86 has a primary fitness effect on the immune escape rather than ACE2 binding. ZCB11 binding is determined by hydrogen bonds, including those formed between N477, K478, N487, N460 while the second interface between ZCB11 and RBM is stabilized by the hydrophobic interaction mediated by L455, F456, and F486 (Figure 12D).^137^ Here again, the mode of antibody binding is severely affected by F486P mutation leading to the large destabilizing change of ΔΔG > 2.5 kcal/mol which may explain the experimentally observed complete abrogation of ZCB11 binding to BA.2.86 variant.^51^

RBD antibodies targeting the Class 3 epitope bind outside of the ACE-2 binding region and often provide a potential for synergistic allosteric effects when combined with RBD class ½ antibodies that interfere with ACE2 binding. Mutational profiling for class 3 RBD antibodies was performed for the S-RBD complex with antibody A19-46.1 (pdb id 7TC9)^138^, Omicron RBD with S309 (sotrovimab) (pdb id 7YAD)^139^, S-RBD with COV2-2130 (cilgavimab) (pdb id 8D8Q)^133^, S-RBD with LY-CoV1404 (bebtelovimab) (pdb I 7MMO)^140^, and S-RBD beta variant complex with Beta-50 and Beta-54 (pdb id 7Q0H).^141^ The results of mutational scanning for this class of antibodies revealed a different group of energetic hotspots where BA.2.86 mutations result in substantial loss of binding affinity (Figure 13). For A19-46.1 antibody the large destabilizing free energy changes were observed for N450D, L452W and A484K mutations (Figure 13A) which is consistent with the experiments showing substantial detrimental losses in IC50 binding of ∼5 fold change for N450D, > 220 fold for L452W and ∼17 fold for A484K mutations.^51^ Structural mapping of the A19-46.1 binding to the RBD illustrated a different mode of interactions for class 3 RBD antibodies (Figure 14A).

**Figure 14.**
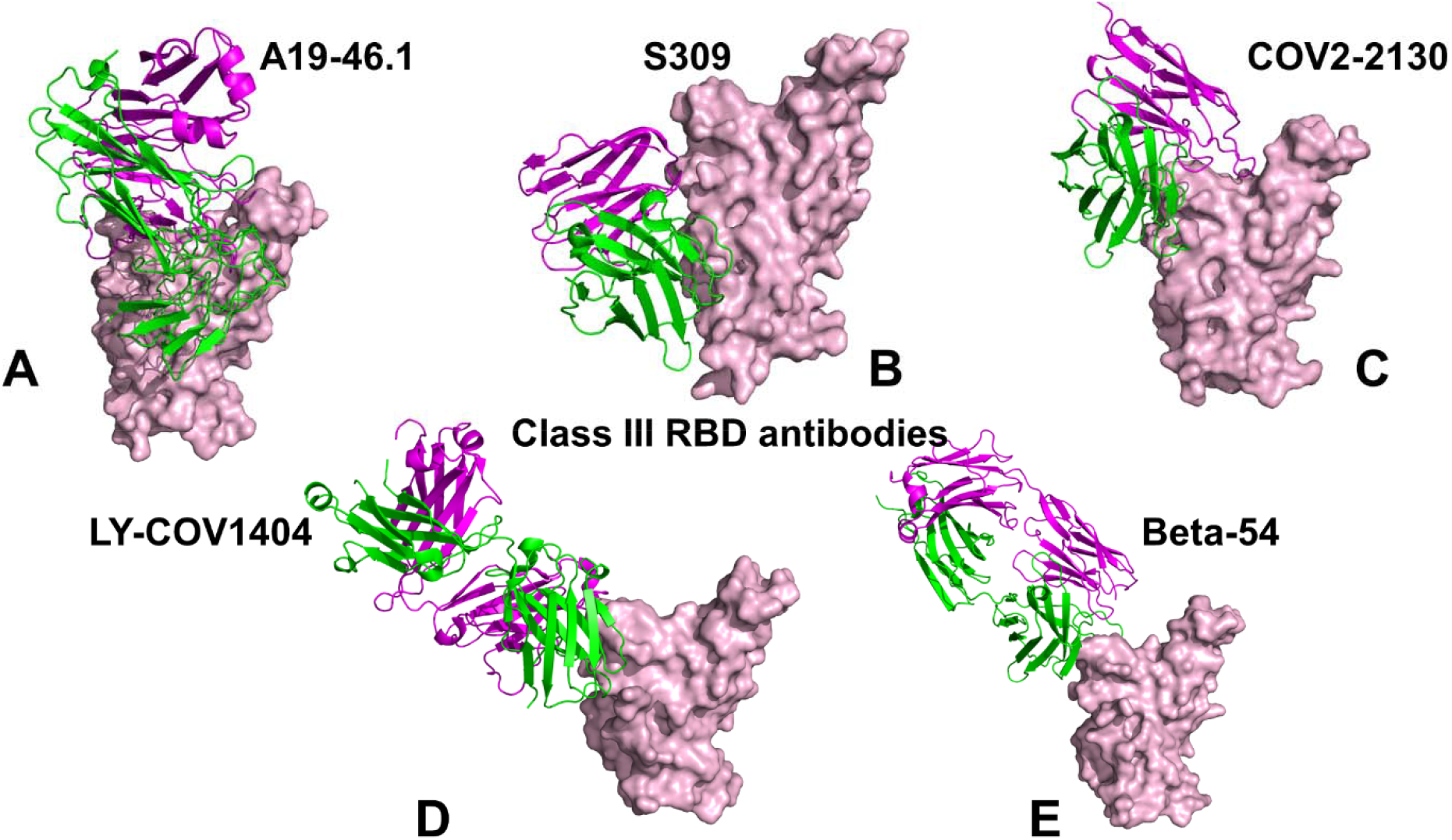
The structures of RBD-antibody complexes for class 3 RBD-targeted antibodies. The structure of the S-RBD complex with A19-46.1 (A), Omicron S-RBD with S309 (B), S-RBD with COV2-2130 (C), S-RBD with LY-CoV1404 (D) and S-RBD beta variant complex with Beta-54 (E). The RBD is shown in pink-colored surface representation and the antibodies are shown in ribbons (heavy chain in magenta and light chain in green-colored ribbons).

The analysis of S309 binding with Omicron RBD pointed to an appreciable loss of binding due to K356T mutation (Figure 13B). These observations reproduced the experimental effects of BA.2.86 mutations on S309 binding in the BA.2 background which demonstrated that K356T is a single dominant hotspot causing almost 20 fold loss in binding.^51^ Structural analysis of RBD- S309 complex (Figure 14B) showed that S309 forms hydrogen bonds with the RBD residues N334, E340, N343, T345, R346, and K356T of BA.2.86 can singlehandedly cause antibody resistance.^139^ Similarly, a single BA.2.86 mutational change N450D resulted in considerable ΔΔG = 1.5 kcal/mol binding loss while other BA.2.86 substitutions caused only moderate changes (Figure 13C, 14C). These results corroborate with the neutralization data showing > 240 fold change in IC50 of BA.2 RBD binding upon single N450D mutation.^51^ The mutational profiles for S-RBD with LY-CoV1404 (Figure 13D) and S-RBD beta variant complex with Beta-54 antibodies (Figure 13E) were similar disclosing the key role of BA.2.86 mutation V445H. This mutational change resulted in a dramatic binding loss of ΔΔG = 2.5 kcal/mol for LY-CoV1404 (Figure 13D) and ΔΔG = 2.5 kcal/mol for Beta-54 antibody (Figure 13E).

Structural analysis of the S-RBD complexes with LY-CoV1404 (Figure 14D) and Beta-54 (Figure 14E) highlighted a very similar binding mode where both antibodies recognize a RBD region that only barely overlaps with the ACE2 binding interface, particularly at positions V445H and G446S. Strikingly, our analysis suggested that a single BA.2.86 mutation V445H could elicit resistance to the entire class 3 of RBD antibodies. These findings are in accordance with the experimental observations^51^ that highlighted > 1000 fold loss in LY-CoV1404 binding upon V445H mutation. The results of mutational scanning of a broad range of RBD-antibodies complexes targeting class1, class 2 and class 3 epitopes demonstrated how different BA.2.86 mutations including N450D, L452W, N460K, N481K and F486P can incur substantial binding free energy losses and lead to immune evasion. These results are also in agreement with antigenicity characterization of BA.2.86 which demonstrated that N450D, K356T, L452W, A484K, V483del, and V445H are responsible for the enhanced immune evasion of BA.2.86 as compared to another highly evasive XBB.1.5 variant.^51^

At the same time, R403K and R493Q could often confer a moderate degree of sensitization to neutralization by certain antibodies but may not be the primary determinants of antibody binding. The results suggest that both R403K and R493Q modifications can play a significant role in both receptor affinity and modulation of antibody binding but have a stronger effect on the BA.2.86 RBD-ACE2 binding.

## Discussion

The latest biophysical studies propose possible scenarios underlying activity and binding of the BA.2.86 S protein by analyzing antigenic cartography and binding data on BA.2.86 neutralization by distinct classes of antibodies.^51-56^ These experimental studies suggested that the increased viral fitness of BA.2.86 through acquisition a wide range of unique RBD mutations may be largely due to immune pressure that promotes convergent evolution. There is however lack of molecular details on structure, dynamics and binding energetics of the BA.2.86 RBD binding with the ACE2 receptor and antibodies to rationalize the experimental data and provide atomistic basis for the proposed molecular mechanisms. To develop accurate and robust atomistic models of BA.2.86 RBD structure and conformational ensembles, we performed comparative structural prediction of the BA.2 and BA.2.86 RBD-ACE2 complexes using AF2 approach with shallow MSA depth. Despite a significant accumulation of unique BA.2.86 mutations in the ACE2 binding interface, we observed relatively moderate structural response of the BA.2.86 RBD that preserves the RBD fold and topology of the ACE2 binding interface. Our results showed that AF2 models with the shallow MSA depth are robust in accurately capturing the experimental structure and that pLDDT statistical assessment of the AF2 models can be reliably used to identify flexible RBD regions and predict conformational heterogeneity of these adaptable regions with atomistic accuracy. By combining atomistic MD simulations with systematic mutational scanning of the RBD-ACE2 interfaces, we found that R403K and reversed R493Q mutations contribute significantly to the binding affinity of the BA.2.86 RBD while other mutations (D339H, K356T, V445H, G446S, N450D, L452W, N460K, N481K and A484K) have only minor effect and may have emerged primarily to strengthen immune evasion potential rather than improving ACE2 binding. By estimating the binding free energy changes of mutations in the BA.2 and BA.2.86 backgrounds, we found that in the BA.2.86 background K403 and Q493 positions are far more significant for the ACE2 binding, revealing a potential epistatic effect of R403K and R493Q mutations in the BA.2.86 RBD-ACE2 complex which explains the experiments.^56^ The pairs of Omicron substitutions with strong epistasis tend to be spatially proximal as evident from most recent findings of epistatic interactions for the L455F/F456L flip that synergistically enhances the ACE2 binding both thermodynamically and kinetically.^128^ The results of our study suggest that evolution of latest Omicron variants fits into a perspective of balancing mechanisms between immune evasion and ACE2 binding. We argue that emergence of BA.2.86 sublineage may have been particularly driven by expanding the scope of mutations to boost immune resistance while employing a relatively focused group of Omicron mutational hotspots (including R403K, R493Q and F486P) to modulate the ACE2 binding affinity. According to our analysis, a much broader range of emerged BA.2.86 RBD mutations can induce robust and broad immune escape without significantly compromising the ACE2 receptor binding. Recent studies indicated that ACE2 binding can often be synergistically modulated and amplified via epistatic interactions of physically proximal binding hotspots, including Y501, R498, L455 and F456 residues.^128,142,143^ The results of forward and reversed mutational scanning indicated potential epistatic effects induced by R403K and R493Q mutations. We suggest that through epistatic interactions of several binding hotspots (R403K, R493K, R498, Y501) the latest Omicron variants (XBB and BA.2.86) may amplify their individual favorable effect on ACE2 binding without invoking “help” of other mutational sites that are primarily implicated in the immune evasion.

Structure-based computational modeling and mutational scanning of the RBD binding interfaces with different classes of RBD antibodies characterized the role of specific BA.2.86 mutations in eliciting broad resistance to neutralization against distinct epitope classes. The results showed that N460K and F486P are primary determinants of BA.2.86 resistance to class 1 and 2 RBD antibodies, while other BA.2.86 mutational positions (such as V445H, N450D, L452W and A484K) are recruited to block binding of class 3 RBD antibodies. Based on the correspondence between the computational results and biochemical experiments^51-56^, we infer that the primary role of most BA.2.86 mutations may be to ensure a broad resistance against different classes of RBD antibodies, while several important BA.2.86 mutations R403K and R493Q could confer the improved ACE2 binding affinity. The results of our study support a recently proposed hypothesis that the impact of the increased ACE2 binding affinity on viral fitness is more universal and is mediated through a group of common binding hotspots.^129^ In contrast, the effect of immune evasion could be more variant-dependent and modulated through recruitment of mutational sites in various adaptable RBD regions. This mechanism is supported by the evidence that high-frequency mutations in the Omicron lineages are more likely to confer a stronger resistance to the potent antibodies against the variant than to materially improve ACE2 binding affinity.^129^

## Conclusions

In this study, we have provided the first comprehensive computational analysis of the BA.2.86 S-RBD protein structure, dynamics and energetics from the perspective of immune evasion and ACE2 binding affinity properties. We combined AF2 structural modeling, atomistic MD simulations and systematic binding energetics analysis of the BA.2.86 S protein with ACE2 receptor and a panel of distinct classes of RBD antibodies to quantify molecular mechanisms underlying BA.2.86 binding and immune evasion. Several AF2-based structural modeling pipelines were adapted to predict structure and functionally relevant conformational ensemble of the BA.2.86 RBD-ACE2 complexes. AF2-based structural modeling with varied MSA depth produced robust conformational predictions of the BA.2.86 RBD-ACE2 complex revealing important functional variations are localized in two RBD loops (resides 444-452 and 475-481) that harbor important mutational sites. Atomistic MD simulations of the BA.2 and BA.2.86 RBD-ACE2 complexes revealed important common and variant-specific dynamic signatures exemplified by modulation of flexibility of functional RBD loops. Importantly, we found that AF2 predicted ensemble of functional conformations using pLDDT metric can accurately capture the main dynamics signatures of the RBD-ACE2 complexes obtained from microscond MD simulations. Using the conformational ensembles obtained from AF2 MSA depth predictions and MD simulations we performed a systematic mutational scanning of the BA.2 and BA.2.86 RBD residues in the RBD-ACE2 complexes revealing a group of conserved hydrophobic hotspots and critical variant-specific contributions of R403K, F486P and R493Q mutations. Our analysis also estimated the role of the BA.2 and BA.2.86 backgrounds on the magnitude of binding free energy changes induced by mutations which revealed potential epistatic effects of R403K and R403Q BA.2.86 substitutions. Our data suggested that BA.2.86 lineage may have evolved to outcompete other Omicron subvariants by improving immune suppression while balancing binding affinity with ACE2 via through compensatory effect of R493Q and F486P mutations. This interpretation is consistent with the notion that acquisition of functionally balanced substitutions to optimize multiple fitness tradeoffs between immune evasion, high ACE2 affinity and sufficient conformational adaptability may potentially be a common strategy of SARS-CoV-2 evolution executed by Omicron subvariants.

To examine immune evasion properties of BA.2.86 in atomistic detail and clarify role of the BA.2.86 mutations in immune escape against distinct classes of RBD-targeted antibodies, we performed systematic structure-based mutational scanning of the S protein binding interfaces with distinct classes of RBD antibodies that retained activity against BA.2 and XBB.1.5 variants but displayed significantly reduced or completely abolished neutralization against BA.2.86 variant. A comparison of the computationally predicted binding changes with the experimental data in which each of the BA.2.86 mutants was evaluated for antibody binding showed a robust quantitative agreement. We found that N460K and F486P are primary determinants of BA.2.86 resistance to class 1 and 2 RBD antibodies, while other BA.2.86 mutational positions (such as V445H, N450D, L452W and A484K) are recruited to block binding of class 3 RBD antibodies. These findings highlighted the significant role of BA.2.86 mutations that may have emerged to ensure broad resistance against different classes of RBD antibodies. Our study provided support to a mechanism in which acquisition of functionally balanced substitutions to optimize multiple fitness tradeoffs between immune evasion, ACE2 affinity and sufficient conformational adaptability may potentially be a common strategy of SARS-CoV-2 evolution executed by Omicron subvariants where the latest wave of mutations.

## Data Availability Statement

Data is fully contained within the article and Supplementary Materials. Crystal structures were obtained and downloaded from the Protein Data Bank (http://www.rcsb.org). All simulations were performed using the all-atom additive CHARMM36M protein force field that can be obtained from http://mackerell.umaryland.edu/charmm_ff.shtml. The rendering of protein structures was done with UCSF ChimeraX package (https://www.rbvi.ucsf.edu/chimerax/) and Pymol (https://pymol.org/2/). The software tools used in this study are freely available at GitHub sites: https://github.com/deepmind/alphafold; https://github.com/sokrypton/ColabFold/; https://github.com/RSvan/SPEACH_AF; https://www.github.com/HWaymentSteele/AFCluster; https://github.com/nextstrain; https://github.com/Amber-MD/cpptraj; https://github.com/smu-tao-group/protein-VAE.

All the data obtained in this work (including simulation trajectories, topology and parameter files, the software tools, and the in-house scripts are freely available at ZENODO website https://zenodo.org/records/10140418.

## Supporting Information

An overview of the phylogenetic analysis of BA.2.86 and BA.2 variants (Figure S1). The alignment of top five BA.2 RBD models with the crystallographic conformation of the BA.2 RBD (Figure S2). The alignment of top five BA.2.86 RBD models with the crystallographic conformation of the BA.2 RBD (Figure S3). The post-processing AF2 analysis of predictions for the BA.2.86 RBD-ACE2 complex includes the pLDDT per residue for the top five models and the predicted alignment error (PAE) for the top five models obtained from AF2 predictions.

(Figure S4). Structural analysis of conservation of the C488-C480 disulfide bond in BA.2 and BA.2.86 RBD-ACE2 complexes (Figure S5). Conformational dynamics profiles obtained MD simulations of the ACE2 residues in BA.2 RBD-ACE2 and BA.2.86 RBD-ACE2 complexes (Figure S6). The residue-based MM-GBSA binding energy contributions for the BA.2 and BA.2.86 RBD-ACE2 complexes (Figure S7). The distribution of the electrostatic potentials on the molecular surface of the S-RBD-ACE2 complexes for BA.1, BA.2, BA.2.75. BA.3, BA.4/BA.5 and XBB.1 variants (Figure S8). This material is available free of charge via the Internet at http://pubs.acs.org.

## Author Contributions

Conceptualization, G.V.; methodology, N.R., M.A., G.C., S.X., G.V. P.T.; software, N.R., S.X., M.A., G.G., G.V and P.T.; validation, N.R., G.V.; formal analysis, N.R., G.V., M.A., G.G., S.X., and P.T.; investigation, N.R., M.A., G.C., G.V. and P.T.; resources, N.R., G.V., M.A. S.X., and G.V.; data curation, N.R., M.A., G.C.,G.V.; writing— original draft preparation, N.R., M.A., G.V.; writing—review and editing, N.R. and G.V.; visualization, N.R., M.A., G.C., S.X., G.V. G.V.; supervision, G.V.; project administration, G.V.; funding acquisition, P.T. and G.V. All authors have read and agreed to the published version of the manuscript.

## Acknowledgments

G.V acknowledges support from Schmid College of Science and Technology at Chapman University for providing computing resources at the Keck Center for Science and Engineering.

## References

(1) Tai, W.; He, L.; Zhang, X.; Pu, J.; Voronin, D.; Jiang, S.; Zhou, Y.; Du, L. Characterization of the receptor-binding domain (RBD) of 2019 novel coronavirus: implication for development of RBD protein as a viral attachment inhibitor and vaccine. Cell. Mol. Immunol. 2020, 17, 613–620. doi: 10.1038/s41423-020-0400-4.

(2) Wang, Q.; Zhang, Y.; Wu, L.; Niu, S.; Song, C.; Zhang, Z.; Lu, G.; Qiao, C.; Hu, Y.; Yuen, K. Y.; Wang, Q.; Zhou, H.; Yan, J.; Qi, J. Structural and functional basis of SARS-CoV-2 entry by using human ACE2. Cell 2020, 181, 894–904.e9. doi: 10.1016/j.cell.2020.03.045.

(3) Walls, A. C.; Park, Y. J.; Tortorici, M. A.; Wall, A.; McGuire, A. T.; Veesler, D. Structure, Function, and Antigenicity of the SARS-CoV-2 Spike Glycoprotein. Cell 2020, 181, 281–292.e6. doi: 10.1016/j.cell.2020.02.058.

(4) Wrapp, D.; Wang, N.; Corbett, K. S.; Goldsmith, J. A.; Hsieh, C. L.; Abiona, O.; Graham, B. S.; McLellan, J. S. Cryo-EM structure of the 2019-nCoV spike in the prefusion conformation. Science 2020, 367, 1260–1263. doi: 10.1126/science.abb2507.

(5) Cai, Y.; Zhang, J.; Xiao, T.; Peng, H.; Sterling, S. M.; Walsh, R. M., Jr.; Rawson, S.; Rits-Volloch, S.; Chen, B. Distinct conformational states of SARS-CoV-2 spike protein. Science 2020, 369, 1586–1592. doi: 10.1126/science.abd4251.

(6) Hsieh, C. L.; Goldsmith, J. A.; Schaub, J. M.; DiVenere, A. M.; Kuo, H. C.; Javanmardi, K.; Le, K. C.; Wrapp, D.; Lee, A. G.; Liu, Y., Chou, C.W.; Byrne, P.O.; Hjorth, C.K.; Johnson, N.V.; Ludes-Meyers J.; Nguyen, A.W.; Park, J.; Wang, N.; Amengor, D.; Lavinder, J.J.; Ippolito, G.C.; Maynard, J.A.; Finkelstein, I.J.; McLellan, J.S. Structure-based design of prefusion-stabilized SARS-CoV-2 spikes. Science 2020, 369, 1501–1505. doi: 10.1126/science.abd0826.

(7) Henderson, R.; Edwards, R. J.; Mansouri, K.; Janowska, K.; Stalls, V.; Gobeil, S. M. C.; Kopp, M.; Li, D.; Parks, R.; Hsu, A. L., Borgnia, M.J.; Haynes, B.F.; Acharya, P. Controlling the SARS-CoV-2 spike glycoprotein conformation. Nat. Struct. Mol. Biol. 2020, 27, 925–933. doi: 10.1038/s41594-020-0479-4.

(8) McCallum, M.; Walls, A. C.; Bowen, J. E.; Corti, D.; Veesler, D. Structure-guided covalent stabilization of coronavirus spike glycoprotein trimers in the closed conformation. Nat. Struct. Mol. Biol. 2020, 27, 942–949. doi: 10.1038/s41594-020-0483-8.

(9) Xiong, X.; Qu, K.; Ciazynska, K. A.; Hosmillo, M.; Carter, A. P.; Ebrahimi, S.; Ke, Z.; Scheres, S. H. W.; Bergamaschi, L.; Grice, G. L., Zhang, Y.; CITIID-NIHR COVID-19 BioResource Collaboration, Nathan, J.A.; Baker, S.; James, L.C.; Baxendale, H.E.; Goodfellow, I.; Doffinger, R.; Briggs, J.A.G. A thermostable, closed SARS-CoV-2 spike protein trimer. Nat. Struct. Mol. Biol. 2020, 27, 934–941. doi: 10.1038/s41594-020-0478-5.

(10) Costello, S.M.; Shoemaker, S.R.; Hobbs, H.T.; Nguyen, A.W.; Hsieh, C.L.; Maynard, J.A.; McLellan, J.S.; Pak, J.E.; Marqusee, S. The SARS-CoV-2 spike reversibly samples an open-trimer conformation exposing novel epitopes. Nat. Struct. Mol. Biol. 2022, 27, 229–238. doi: 10.1038/s41594-022-00735-5.

(11) McCormick, K.D.; Jacobs, J.L.; Mellors, J.W. The emerging plasticity of SARS-CoV-2. Science 2021, 371, 1306–1308. doi: 10.1126/science.abg4493.

(12) Ghimire, D.; Han, Y.; Lu, M. Structural Plasticity and Immune Evasion of SARS-CoV-2 Spike Variants. Viruses 2022, 14, 1255. 10.3390/v14061255.

(13) Xu, C.; Wang, Y.; Liu, C.; Zhang, C.; Han, W.; Hong, X.; Wang, Y.; Hong, Q.; Wang, S.; Zhao, Q.; Wang, Y.; Yang, Y.; Chen, K.; Zheng, W.; Kong, L.; Wang, F.; Zuo, Q.; Huang, Z.; Cong, Y. Conformational dynamics of SARS-CoV-2 trimeric spike glycoprotein in complex with receptor ACE2 revealed by cryo-EM. Sci. Adv. 2021, 7, eabe5575. doi: 10.1126/sciadv.abe5575.

(14) Benton, D. J.; Wrobel, A. G.; Xu, P.; Roustan, C.; Martin, S. R.; Rosenthal, P. B.; Skehel, J. J.; Gamblin, S. J. Receptor binding and priming of the spike protein of SARS-CoV-2 for membrane fusion. Nature 2020, 588, 327–330. doi: 10.1038/s41586-020-2772-0.

(15) Turoňová, B.; Sikora, M.; Schuerman, C.; Hagen, W. J. H.; Welsch, S.; Blanc, F. E. C.; von Bülow, S.; Gecht, M.; Bagola, K.; Hörner, C.; van Zandbergen, G.; Landry, J.; de Azevedo, N. T. D.; Mosalaganti, S.; Schwarz, A.; Covino, R.; Mühlebach, M. D.; Hummer, G.; Krijnse Locker, J.; Beck, M. In situ structural analysis of SARS-CoV-2 spike reveals flexibility mediated by three hinges. Science 2020, 370, 203–208. doi: 10.1126/science.abd5223.

(16) Lu, M.; Uchil, P. D.; Li, W.; Zheng, D.; Terry, D. S.; Gorman, J.; Shi, W.; Zhang, B.; Zhou, T.; Ding, S.; Gasser, R.; Prevost, J.; Beaudoin-Bussieres, G.; Anand, S. P.; Laumaea, A.; Grover, J. R.; Lihong, L.; Ho, D. D.; Mascola, J.R.; Finzi, A.; Kwong, P. D.; Blanchard, S. C.; Mothes, W. Real-time conformational dynamics of SARS-CoV-2 spikes on virus particles. Cell Host Microbe. 2020, 28, 880–891.e8. doi: 10.1016/j.chom.2020.11.001.

(17) Yang, Z.; Han, Y.; Ding, S.; Shi, W.; Zhou, T.; Finzi, A.; Kwong, P.D.; Mothes, W.; Lu, M. SARS-CoV-2 Variants Increase Kinetic Stability of Open Spike Conformations as an Evolutionary Strategy. mBio 2022, 13, e0322721. doi: 10.1128/mbio.03227-21.

(18) Díaz-Salinas, M.A.; Li, Q.; Ejemel, M.; Yurkovetskiy, L.; Luban, J.; Shen, K.; Wang, Y.; Munro, J.B. Conformational dynamics and allosteric modulation of the SARS-CoV-2 spike. Elife 2022, 11, e75433. doi: 10.7554/eLife.75433.

(19) Han, P.; Li, L.; Liu, S.; Wang, Q.; Zhang, D.; Xu, Z.; Li, X.; Peng, Q.; Su, C.; Huang, B.; Li, D.; Zhang, R.; Tian, M.; Fu, L.; Gao, Y.; Zhao, X.; Liu, K.; Qi, J.; Gao, G. F.; Wang, P. Receptor binding and complex structures of human ACE2 to spike RBD from omicron and delta SARS-CoV-2. Cell 2022, doi: 10.1016/j.cell.2022.01.001.

(20) Saville, J.W.; Mannar, D.; Zhu, X.; Srivastava, S.S.; Berezuk, A.M.; Demers, J.P.; Zhou, S.; Tuttle, K.S.; Sekirov, I.; Kim A.; Li, W.; Dimitrov, D.S.; Subramaniam, S. Structural and biochemical rationale for enhanced spike protein fitness in delta and kappa SARS-CoV-2 variants. Nat. Commun. 2022, 13, 742. doi: 10.1038/s41467-022-28324-6.

(21) Wang, Y.; Liu, C.; Zhang, C.; Wang, Y.; Hong, Q.; Xu, S.; Li, Z.; Yang, Y.; Huang, Z.; Cong, Y. Structural basis for SARS-CoV-2 Delta variant recognition of ACE2 receptor and broadly neutralizing antibodies. Nat. Commun. 2022, 13, 871. doi: 10.1038/s41467-022-28528-w.

(22) Zhang, J.; Xiao, T.; Cai, Y.; Lavine, C.L.; Peng, H.; Zhu, H.; Anand, K.; Tong, P.; Gautam, A.; Mayer, M.L.; Walsh, R.M. Jr.; Rits-Volloch, S.; Wesemann, D.R.; Yang, W.; Seaman, M.S.; Lu, J.; Chen, B. Membrane fusion and immune evasion by the spike protein of SARS-CoV-2 Delta variant. Science 2021, 374, 1353–1360. doi: 10.1126/science.abl9463.

(23) Mannar, D.; Saville, J.W.; Zhu, X.; Srivastava, S.S.; Berezuk, A.M.; Tuttle, K.S.; Marquez, A.C.; Sekirov, I.; Subramaniam, S. SARS-CoV-2 Omicron variant: Antibody evasion and cryo-EM structure of spike protein-ACE2 complex. Science 2022, 375, 760–764. doi: 10.1126/science.abn7760.

(24) Hong, Q.; Han, W.; Li, J.; Xu, S.; Wang, Y.; Xu, C.; Li, Z.; Wang, Y.; Zhang, C.; Huang, Z.; Cong, Y. Molecular basis of receptor binding and antibody neutralization of Omicron. Nature 2022. doi: 10.1038/s41586-022-04581-9.

(25) McCallum, M.; Czudnochowski, N.; Rosen, L.E.; Zepeda, S.K.; Bowen, J.E.; Walls, A.C.; Hauser, K.; Joshi, A.; Stewart, C.; Dillen, J.R.; Powell, A.E.; Croll, T.I.; Nix, J.; Virgin, H.W.; Corti, D.; Snell, G.; Veesler, D. Structural basis of SARS-CoV-2 Omicron immune evasion and receptor engagement. Science 2022, 375, 864–868. doi: 10.1126/science.abn8652.

(26) Yin, W.; Xu, Y.; Xu, P.; Cao, X.; Wu, C.; Gu, C.; He, X.; Wang, X.; Huang, S.; Yuan, Q.; Wu, K.; Hu, W.; Huang, Z.; Liu, J.; Wang, Z.; Jia, F.; Xia, K.; Liu, P.; Wang, X.; Song, B.; Zheng, J.; Jiang, H.; Cheng, X.; Jiang, Y.; Deng, S.J.; Xu, H.E. Structures of the Omicron Spike trimer with ACE2 and an anti-Omicron antibody. Science 2022, 375, 1048–1053. doi: 10.1126/science.abn8863.

(27) Gobeil, S.M.; Henderson, R.; Stalls, V.; Janowska, K.; Huang, X.; May, A.; Speakman, M.; Beaudoin, E.; Manne, K.; Li, D.; Parks, R.; Barr, M.; Deyton, M.; Martin, M.; Mansouri, K.; Edwards, R.J.; Sempowski, G.D.; Saunders, K.O.; Wiehe, K.; Williams, W.; Korber, B.; Haynes, B.F.; Acharya, P. Structural diversity of the SARS-CoV- 2 Omicron spike. bioRxiv, 2022, doi: 10.1101/2022.01.25.477784.

(28) Cui, Z.; Liu, P.; Wang, N.; Wang, L.; Fan, K.; Zhu, Q.; Wang, K.; Chen, R.; Feng, R.; Jia, Z.; Yang, M.; Xu, G.; Zhu, B.; Fu, W.; Chu, T.; Feng, L.; Wang, Y.; Pei, X.; Yang, P.; Xie, X.S.; Cao, L.; Cao, Y.; Wang, X. Structural and functional characterizations of infectivity and immune evasion of SARS-CoV-2 Omicron. Cell 2022, 185, 860–871.e13. doi: 10.1016/j.cell.2022.01.019.

(29) Stalls, V.; Lindenberger, J.; Gobeil, S. M.-C.; Henderson, R.; Parks, R.; Barr, M.; Deyton, M.; Martin, M.; Janowska, K.; Huang, X.; May, A.; Speakman, M.; Beaudoin, E.; Kraft, B.; Lu, X.; Edwards, R. J.; Eaton, A.; Montefiori, D. C.; Williams, W. B.; Saunders, K. O.; Wiehe, K.; Haynes, B. F.; Acharya, P. Cryo-EM Structures of SARS-CoV-2 Omicron BA.2 Spike. Cell Rep. 2022, 39, 111009. doi: 10.1016/j.celrep.2022.111009.

(30) Li, L.; Liao, H.; Meng, Y.; Li, W.; Han, P.; Liu, K.; Wang, Q.; Li, D.; Zhang, Y.; Wang, L.; Fan, Z.; Zhang, Y.; Wang, Q.; Zhao, X.; Sun, Y.; Huang, N.; Qi, J.; Gao, G.F. Structural basis of human ACE2 higher binding affinity to currently circulating Omicron SARS-CoV-2 sub-variants BA.2 and BA.1.1. Cell 2022, 185, 2952–2960.e10. doi: 10.1016/j.cell.2022.06.023.

(31) Xu, Y.; Wu, C.; Cao, X.; Gu, C.; Liu, H.; Jiang, M.; Wang, X.; Yuan, Q.; Wu, K.; Liu, J.; Wang, D.; He, X.; Wang, X.; Deng, S.J.; Xu, H.E.; Yin, W. Structural and biochemical mechanism for increased infectivity and immune evasion of Omicron BA.2 variant compared to BA.1 and their possible mouse origins. Cell Res. 2022, 32, 609–620. doi: 10.1038/s41422-022-00672-4.

(32) Cao, Y.; Yisimayi, A.; Jian, F.; Song, W.; Xiao, T.; Wang, L.; Du, S.; Wang, J.; Li, Q.; Chen, X.; Yu, Y.; Wang, P.; Zhang, Z.; Liu, P.; An, R.; Hao, X.; Wang, Y.; Wang, J.; Feng, R.; Sun, H.; Zhao, L.; Zhang, W.; Zhao, D.; Zheng, J.; Yu, L.; Li, C.; Zhang, N.; Wang, R.; Niu, X.; Yang, S.; Song, X.; Chai, Y.; Hu, Y.; Shi, Y.; Zheng, L.; Li, Z.; Gu, Q.; Shao, F.; Huang, W.; Jin, R.; Shen, Z.; Wang, Y.; Wang, X.; Xiao, J.; Xie, X.S. BA.2.12.1, BA.4 and BA.5 escape antibodies elicited by Omicron infection. Nature 2022, 608, 593-602. doi: 10.1038/s41586-022-04980-y.

(33) Bowen, J.E.; Addetia, A.; Dang, H.V.; Stewart, C.; Brown, J.T.; Sharkey, W.K.; Sprouse, K.R.; Walls, A.C.; Mazzitelli, I.G.; Logue, J.K.; Franko, N.M.; Czudnochowski, N.; Powell, A.E.; Dellota, E. Jr.; Ahmed, K.; Ansari, A.S.; Cameroni, E.; Gori, A.; Bandera, A.; Posavad, C.M.; Dan, J.M.; Zhang, Z.; Weiskopf, D.; Sette, A.; Crotty, S.; Iqbal, N.T.; Corti, D.; Geffner, J.; Snell, G.; Grifantini, R.; Chu, H.Y.; Veesler, D. Omicron spike function and neutralizing activity elicited by a comprehensive panel of vaccines. Science 2022, 377, 890–894. doi: 10.1126/science.abq0203.

(34) Huo, J.; Dijokaite-Guraliuc, A.; Liu, C.; Zhou, D.; Ginn, H. M.; Das, R.; Supasa, P.; Selvaraj, M.; Nutalai, R.; Tuekprakhon, A.; Duyvesteyn, H. M. E.; Mentzer, A. J.; Skelly, D.; Ritter, T. G.; Amini, A.; Bibi, S.; Adele, S.; Johnson, S. A.; Paterson, N. G.; Williams, M. A.; Hall, D. R.; Plowright, M.; Newman, T. A. H.; Hornsby, H.; de Silva, T. I.; Temperton, N.; Klenerman, P.; Barnes, E.; Dunachie, S. J.; Pollard, A. J.; Lambe, T.; Goulder, P.; Fry, E. E.; Mongkolsapaya, J.; Ren, J.; Stuart, D. I.; Screaton, G. R. A Delicate Balance between Antibody Evasion and ACE2 Affinity for Omicron BA.2.75. Cell Rep. 2023, 42, 111903. doi: 10.1016/j.celrep.2022.111903.

(35) Cao, Y.; Song, W.; Wang, L.; Liu, P.; Yue, C.; Jian, F.; Yu, Y.; Yisimayi, A.; Wang, P.; Wang, Y.; Zhu, Q.; Deng, J.; Fu, W.; Yu, L.; Zhang, N.; Wang, J.; Xiao, T.; An, R.; Wang, J.; Liu, L.; Yang, S.; Niu, X.; Gu, Q.; Shao, F.; Hao, X.; Meng, B.; Gupta, R. K.; Jin, R.; Wang, Y.; Xie, X. S.; Wang, X. Characterization of the Enhanced Infectivity and Antibody Evasion of Omicron BA.2.75. Cell Host Microbe 2022, 30, 1527–1539.e5. doi: 10.1016/j.chom.2022.09.018.

(36) Saito, A.; Tamura, T.; Zahradnik, J.; Deguchi, S.; Tabata, K.; Anraku, Y.; Kimura, I.; Ito, J.; Yamasoba, D.; Nasser, H.; Toyoda, M.; Nagata, K.; Uriu, K.; Kosugi, Y.; Fujita, S.; Shofa, M.; Monira Begum, M.; Shimizu, R.; Oda, Y.; Suzuki, R.; Ito, H.; Nao, N.; Wang, L.; Tsuda, M.; Yoshimatsu, K.; Kuramochi, J.; Kita, S.; Sasaki-Tabata, K.; Fukuhara, H.; Maenaka, K.; Yamamoto, Y.; Nagamoto, T.; Asakura, H.; Nagashima, M.; Sadamasu, K.; Yoshimura, K.; Ueno, T.; Schreiber, G.; Takaori-Kondo, A.; Shirakawa, K.; Sawa, H.; Irie, T.; Hashiguchi, T.; Takayama, K.; Matsuno, K.; Tanaka, S.; Ikeda, T.; Fukuhara, T.; Sato, K. Virological Characteristics of the SARS-CoV-2 Omicron BA.2.75 Variant. Cell Host Microbe 2022, 30, 1540–1555.e15. doi: 10.1016/j.chom.2022.10.003.

(37) Barton, M.I.; MacGowan, S.A.; Kutuzov, M.A.; Dushek, O.; Barton, G.J.; van der Merwe, P.A. Effects of common mutations in the SARS-CoV-2 Spike RBD and its ligand, the human ACE2 receptor on binding affinity and kinetics. Elife 2021, 10, e70658. doi: 10.7554/eLife.70658.

(38) Cao, Y.; Wang, J.; Jian, F.; Xiao, T.; Song, W.; Yisimayi, A.; Huang, W.; Li, Q.; Wang, P.; An, R.; Wang, J.; Wang, Y.; Niu, X.; Yang, S.; Liang, H.; Sun, H.; Li, T.; Yu, Y.; Cui, Q.; Liu, S.; Yang, X.; Du, S.; Zhang, Z.; Hao, X.; Shao, F.; Jin, R.; Wang, X.; Xiao, J.; Wang, Y.; Xie, X.S. Omicron escapes the majority of existing SARS-CoV-2 neutralizing antibodies. Nature 2022, 602, 657–663. doi: 10.1038/s41586-021-04385-3.

(39) Liu, L.; Iketani, S.; Guo, Y.; Chan, J.F.; Wang, M.; Liu, L.; Luo, Y.; Chu, H.; Huang, Y.; Nair, M.S.; Yu, J.; Chik, K. K.; Yuen, T.T.; Yoon, C.; To, K.K.; Chen, H.; Yin, M.T.; Sobieszczyk, M.E.; Huang, Y.; Wang, H.H.; Sheng, Z.; Yuen, K.Y.; Ho, D.D. Striking antibody evasion manifested by the Omicron variant of SARS-CoV-2. Nature 2022, 602, 676–681. doi: 10.1038/s41586-021-04388-0.

40. A. Zhang, J.; Cai, Y.; Lavine, C.L.; Peng, H.; Zhu, H.; Anand, K.; Tong, P.; Gautam,; Mayer, M.L.; Rits-Volloch, S.; Wang, S.; Sliz, P.; Wesemann, D.R.; Yang, W.; Seaman, M.S.; Lu, J.; Xiao, T.; Chen, B. Structural and functional impact by SARS-CoV-2 Omicron spike mutations. Cell Rep. 2022, 39, 110729. doi: 10.1016/j.celrep.2022.110729.

(40) Tuekprakhon, A.; Nutalai, R.; Dijokaite-Guraliuc, A.; Zhou, D.; Ginn, H.M.; Selvaraj, M.; Liu, C.; Mentzer, A.J.; Supasa, P.; Duyvesteyn, H.M.E.; Das, R.; Skelly, D.; Ritter, T.G.; Amini, A.; Bibi, S.; Adele, S.; Johnson, S.A.; Constantinides, B.; Webster, H.; Temperton, N.; Klenerman, P.; Barnes, E.; Dunachie, S.J.; Crook, D.; Pollard, A.J.; Lambe, T.; Goulder, P.; Paterson, N.G.; Williams, M.A.; Hall DR; OPTIC Consortium; ISARIC4C Consortium; Fry, E.E.; Huo, J.; Mongkolsapaya, J.; Ren, J.; Stuart, D.I.; Screaton, G.R. Antibody escape of SARS-CoV-2 Omicron BA.4 and BA.5 from vaccine and BA.1 serum. Cell 2022, 185, 2422–2433.e13. doi: 10.1016/j.cell.2022.06.005.

(41) Kimura, I.; Yamasoba, D.; Tamura, T.; Nao, N.; Suzuki, T.; Oda, Y.; Mitoma, S.; Ito, J.; Nasser, H.; Zahradnik, J.; Uriu, K.; Fujita, S.; Kosugi, Y.; Wang, L.; Tsuda, M.; Kishimoto, M.; Ito, H.; Suzuki, R.; Shimizu, R.; Begum, M. M.; Yoshimatsu, K.; Kimura, K. T.; Sasaki, J.; Sasaki-Tabata, K.; Yamamoto, Y.; Nagamoto, T.; Kanamune, J.; Kobiyama, K.; Asakura, H.; Nagashima, M.; Sadamasu, K.; Yoshimura, K.; Shirakawa, K.; Takaori-Kondo, A.; Kuramochi, J.; Schreiber, G.; Ishii, K. J.; Hashiguchi, T.; Ikeda, T.; Saito, A.; Fukuhara, T.; Tanaka, S.; Matsuno, K.; Sato, K. Virological Characteristics of the SARS-CoV-2 Omicron BA.2 Subvariants, Including BA.4 and BA.5. Cell 2022, 185, 3992–4007.e16. doi: 10.1016/j.cell.2022.09.018.

(42) Qu, P.; Evans, J. P.; Zheng, Y.-M.; Carlin, C.; Saif, L. J.; Oltz, E. M.; Xu, K.; Gumina, R. J.; Liu, S.-L. Evasion of Neutralizing Antibody Responses by the SARS-CoV-2 BA.2.75 Variant. Cell Host Microbe 2022, 30, 1518–1526.e4. doi: 10.1016/j.chom.2022.09.015.

(43) Wang, Q.; Guo, Y.; Iketani, S.; Nair, M.S.; Li, Z.; Mohri, H.; Wang, M.; Yu, J.; Bowen, A.D.; Chang, J.Y.; et al. Antibody evasion by SARS-CoV-2 Omicron subvariants BA.2.12.1, BA.4 and BA.5. Nature 2022, 608, 603–608. 10.1038/s41586-022-05053-w.

(44) Callaway E. Coronavirus variant XBB.1.5 rises in the United States - is it a global threat? Nature 2023, 613, 222–223. doi: 10.1038/d41586-023-00014-3.

(45) Wang, Q.; Iketani, S.; Li, Z.; Liu, L.; Guo, Y.; Huang, Y.; Bowen, A. D.; Liu, M.; Wang, M.; Yu, J.; Valdez, R.; Lauring, A. S.; Sheng, Z.; Wang, H. H.; Gordon, A.; Liu, L.; Ho, D. D. Alarming Antibody Evasion Properties of Rising SARS-CoV-2 BQ and XBB Subvariants. Cell 2023, 186, 279–286.e8. 10.1016/j.cell.2022.12.018.

(46) Yue, C.; Song, W.; Wang, L.; Jian, F.; Chen, X.; Gao, F.; Shen, Z.; Wang, Y.; Wang, X.; Cao, Y. ACE2 Binding and Antibody Evasion in Enhanced Transmissibility of XBB.1.5. Lancet Infect. Dis. 2023, 23, 278–280. 10.1016/s1473-3099(23)00010-5.

(47) Hoffmann, M.; Arora, P.; Nehlmeier, I.; Kempf, A.; Cossmann, A.; Schulz, S. R.; Morillas Ramos, G.; Manthey, L. A.; Jäck, H.-M.; Behrens, G. M. N.; Pöhlmann, S. Profound Neutralization Evasion and Augmented Host Cell Entry Are Hallmarks of the Fast-Spreading SARS-CoV-2 Lineage XBB.1.5. Cell Mol Immunol. 2023, 1–4. doi: 10.1038/s41423-023-00988-0.

(48) Tamura, T.; Ito, J.; Uriu, K.; Zahradnik, J.; Kida, I.; Anraku, Y.; Nasser, H.; Shofa, M.; Oda, Y.; Lytras, S.; Nao, N.; Itakura, Y.; Deguchi, S.; Suzuki, R.; Wang, L.; Begum, M. M.; Kita, S.; Yajima, H.; Sasaki, J.; Sasaki-Tabata, K.; Shimizu, R.; Tsuda, M.; Kosugi, Y.; Fujita, S.; Pan, L.; Sauter, D.; Yoshimatsu, K.; Suzuki, S.; Asakura, H.; Nagashima, M.; Sadamasu, K.; Yoshimura, K.; Yamamoto, Y.; Nagamoto, T.; Schreiber, G.; Maenaka, K.; Ito, H.; Misawa, N.; Kimura, I.; Suganami, M.; Chiba, M.; Yoshimura, R.; Yasuda, K.; Iida, K.; Ohsumi, N.; Strange, A. P.; Takahashi, O.; Ichihara, K.; Shibatani, Y.; Nishiuchi, T.; Kato, M.; Ferdous, Z.; Mouri, H.; Shishido, K.; Sawa, H.; Hashimoto, R.; Watanabe, Y.; Sakamoto, A.; Yasuhara, N.; Suzuki, T.; Kimura, K.; Nakajima, Y.; Nakagawa, S.; Wu, J.; Shirakawa, K.; Takaori-Kondo, A.; Nagata, K.; Kazuma, Y.; Nomura, R.; Horisawa, Y.; Tashiro, Y.; Kawai, Y.; Irie, T.; Kawabata, R.; Motozono, C.; Toyoda, M.; Ueno, T.; Hashiguchi, T.; Ikeda, T.; Fukuhara, T.; Saito, A.; Tanaka, S.; Matsuno, K.; Takayama, K.; Sato, K. Virological Characteristics of the SARS-CoV-2 XBB Variant Derived from Recombination of Two Omicron Subvariants. Nat Commun. 2023, 14, 2800. doi: 10.1038/s41467-023-38435-3.

(49) Hadfield, J.; Megill, C.; Bell, S. M.; Huddleston, J.; Potter, B.; Callender, C.; Sagulenko, P.; Bedford, T.; Neher, R. A. Nextstrain: Real-Time Tracking of Pathogen Evolution. Bioinformatics 2018, 34, 4121–4123. doi: 10.1093/bioinformatics/bty407.

(50) Roemer, C.; Sheward, D. J.; Hisner, R.; Gueli, F.; Sakaguchi, H.; Frohberg, N.; Schoenmakers, J.; Sato, K.; O’Toole, Á.; Rambaut, A.; Pybus, O. G.; Ruis, C.; Murrell, B.; Peacock, T. P. SARS-CoV-2 Evolution in the Omicron Era. Nat. Microbiol. 2023, 8, 1952–1959. 10.1038/s41564-023-01504-w.

(51) Wang, Q.; Guo, Y.; Liu, L.; Schwanz, L. T.; Li, Z.; Nair, M. S.; Ho, J.; Zhang, R. M.; Iketani, S.; Yu, J.; Huang, Y.; Qu, Y.; Valdez, R.; Lauring, A. S.; Huang, Y.; Gordon, A.; Wang, H. H.; Liu, L.; Ho, D. D. Antigenicity and Receptor Affinity of SARS-CoV-2 BA.2.86 Spike. Nature 2023. 10.1038/s41586-023-06750-w.

(52) Yang, S.; Yu, Y.; Jian, F.; Song, W.; Yisimayi, A.; Chen, X.; Xu, Y.; Wang, P.; Wang, J.; Yu, L.; Niu, X.; Wang, J.; Xiao, T.; An, R.; Wang, Y.; Gu, Q.; Shao, F.; Jin, R.; Shen, Z.; Wang, Y.; Cao, Y. Antigenicity and Infectivity Characterization of SARS-CoV-2 BA.2.86. Lancet Infect Dis. 2023, 23, e457–e459. doi: 10.1016/S1473-3099(23)00573-X.

(53) Lasrado, N.; Collier, A. Y.; Hachmann, N. P.; Miller, J.; Rowe, M.; Schonberg, E. D.; Rodrigues, S. L.; LaPiana, A.; Patio, R. C.; Anand, T.; Fisher, J.; Mazurek, C. R.; Guan, R.; Wagh, K.; Theiler, J.; Korber, B. T.; Barouch, D. H. Neutralization Escape by SARS-CoV-2 Omicron Subvariant BA.2.86. Vaccine 2023, 41, 6904–6909. doi: 10.1016/j.vaccine.2023.10.051.

(54) Qu, P.; Xu, K.; Faraone, J. N.; Goodarzi, N.; Zheng, Y.-M.; Carlin, C.; Bednash, J. S.; Horowitz, J. C.; Mallampalli, R. K.; Saif, L. J.; Oltz, E. M.; Jones, D.; Gumina, R. J.; Liu, S.-L. Immune Evasion, Infectivity, and Fusogenicity of SARS-CoV-2 Omicron BA.2.86 and FLip Variants, bioRxiv 2023. doi: 10.1101/2023.09.11.557206.

(55) Uriu, K.; Ito, J.; Kosugi, Y.; Tanaka, Y. L.; Mugita, Y.; Guo, Z.; Hinay, A. A., Jr.; Putri, O.; Kim, Y.; Shimizu, R.; Begum, M. M.; Jonathan, M.; Saito, A.; Ikeda, T.; Sato, K. Transmissibility, Infectivity, and Immune Resistance of the SARS-CoV-2 BA.2.86 Variant. bioRxiv 2023. doi 10.1101/2023.09.07.556636

(56) Tamura, T.; Mizuma, K.; Nasser, H.; Deguchi, S.; Padilla-Blanco, M.; Uriu, K.; Tolentino, J. E. M.; Tsujino, S.; Suzuki, R.; Kojima, I.; Nao, N.; Shimizu, R.; Jonathan, M.; Kosugi, Y.; Guo, Z.; Hinay, A. A., Jr.; Putri, O.; Kim, Y.; Tanaka, Y. L.; Asakura, H.; Nagashima, M.; Sadamasu, K.; Yoshimura, K.; Saito, A.; Ito, J.; Irie, T.; Zahradnik, J.; Ikeda, T.; Takayama, K.; Matsuno, K.; Fukuhara, T.; Sato, K. Virological Characteristics of the SARS-CoV-2 BA.2.86 Variant. bioRxiv 2023. doi: 10.1101/2023.11.02.565304.

(57) Casalino, L.; Gaieb, Z.; Goldsmith, J.A.; Hjorth, C.K.; Dommer, A. C.; Harbison, A. M.; Fogarty, C. A.; Barros, E. P.; Taylor, B. C.; McLellan, J.S., et al. Beyond shielding: The roles of glycans in the SARS-CoV-2 spike potein. ACS Cent. Sci. 2020, 6, 1722–1734.

(58) Barros, E. P.; Casalino, L.; Gaieb, Z.; Dommer, A. C.; Wang, Y.; Fallon, L.; Raguette, L.; Belfon, K.; Simmerling, C.; Amaro, R. E. The Flexibility of ACE2 in the Context of SARS-CoV-2 Infection. Biophys J. 2021, 120, 1072–1084. doi: 10.1016/j.bpj.2020.10.036.

(59) Mehdipour, A. R.; Hummer, G. Dual Nature of Human ACE2 Glycosylation in Binding to SARS-CoV-2 Spike. Proc. Natl. Acad. Sci. U. S. A. 2021, 118, e2100425118. doi: 10.1073/pnas.2100425118

(60) Sztain, T.; Ahn, S.H.; Bogetti, A.T.; Casalino, L.; Goldsmith, J.A.; Seitz, E.; McCool, R.S.; Kearns, F.L.; Acosta-Reyes, F.; Maji, S.; Mashayekhi, G.; McCammon, J.A.; Ourmazd, A.; Frank, J.; McLellan, J.S.; Chong, L.T.; Amaro, R.E. A glycan gate controls the opening of the SARS-CoV-2 spike protein. Nat. Chem. 2021, doi: 10.1038/s41557-021-00758-3.

(61) Pang, Y. T.; Acharya, A.; Lynch, D. L.; Pavlova, A.; Gumbart, J. C. SARS-CoV-2 Spike Opening Dynamics and Energetics Reveal the Individual Roles of Glycans and Their Collective Impact. *Commun*. Biol. 2022, 5, 1170. doi: 10.1038/s42003-022-04138-6.

(62) Zimmerman, M. I.; Porter, J. R.; Ward, M. D.; Singh, S.; Vithani, N.; Meller, A.; Mallimadugula, U. L.; Kuhn, C. E.; Borowsky, J. H.; Wiewiora, R. P., Hurley, M.F.D.; Harbison, A.M.; Fogarty, C.A.; Coffland, J.E.; Fadda, E.; Voelz, V.A.; Chodera, J.D.; Bowman, G.R. SARS-CoV-2 simulations go exascale to predict dramatic spike opening and cryptic pockets across the proteome. Nat. Chem. 2021, 13, 651–659. doi: 10.1038/s41557-021-00707-0.

(63) Mori, T.; Jung, J.; Kobayashi, C.; Dokainish, H.M.; Re, S.; Sugita, Y. Elucidation of interactions regulating conformational stability and dynamics of SARS-CoV-2 S-protein. Biophys. J. 2021, 120, 1060–1071. doi: 10.1016/j.bpj.2021.01.012.

(64) Dokainish, H.M.; Re, S.; Mori, T.; Kobayashi, C.; Jung, J.; Sugita, Y. The inherent flexibility of receptor binding domains in SARS-CoV-2 spike protein. Elife 2022, 11, e75720. doi: 10.7554/eLife.75720.

(65) Verkhivker, G.M.; Di Paola, L. Integrated Biophysical Modeling of the SARS-CoV-2 Spike Protein Binding and Allosteric Interactions with Antibodies. J. Phys. Chem. B. 2021, 125, 4596–4619. doi: 10.1021/acs.jpcb.1c00395.

(66) Verkhivker, G.M.; Agajanian, S.; Oztas, D.Y.; Gupta, G. Comparative Perturbation-Based Modeling of the SARS-CoV-2 Spike Protein Binding with Host Receptor and Neutralizing Antibodies: Structurally Adaptable Allosteric Communication Hotspots Define Spike Sites Targeted by Global Circulating Mutations. Biochemistry 2021, 60, 1459–1484. doi: 10.1021/acs.biochem.1c00139.

(67) Verkhivker, G.M.; Agajanian, S.; Oztas, D.Y.; Gupta, G. Dynamic Profiling of Binding and Allosteric Propensities of the SARS-CoV-2 Spike Protein with Different Classes of Antibodies: Mutational and Perturbation-Based Scanning Reveals the Allosteric Duality of Functionally Adaptable Hotspots. J. Chem. Theory Comput. 2021, 17, 4578–4598. doi: 10.1021/acs.jctc.1c00372.

(68) Verkhivker, G.M.; Agajanian, S.; Oztas, D.Y.; Gupta, G. Allosteric Control of Structural Mimicry and Mutational Escape in the SARS-CoV-2 Spike Protein Complexes with the ACE2 Decoys and Miniprotein Inhibitors: A Network-Based Approach for Mutational Profiling of Binding and Signaling. J. Chem. Inf. Model. 2021, 61, 5172–5191. doi: 10.1021/acs.jcim.1c00766.

(69) Verkhivker, G.; Alshahrani, M.; Gupta, G. Balancing Functional Tradeoffs between Protein Stability and ACE2 Binding in the SARS-CoV-2 Omicron BA.2, BA.2.75 and XBB Lineages: Dynamics-Based Network Models Reveal Epistatic Effects Modulating Compensatory Dynamic and Energetic Changes. Viruses 2023, 15, 1143. 10.3390/v15051143.

(70) Verkhivker, G.; Alshahrani, M.; Gupta, G.; Xiao, S.; Tao, P. Probing Conformational Landscapes of Binding and Allostery in the SARS-CoV-2 Omicron Variant Complexes Using Microsecond Atomistic Simulations and Perturbation-Based Profiling Approaches: Hidden Role of Omicron Mutations as Modulators of Allosteric Signaling and Epistatic Relationships. Phys. Chem. Chem. Phys. 2023, 25, 21245–21266. doi: 10.1039/d3cp02042h.

(71) Xiao, S.; Alshahrani, M.; Gupta, G.; Tao, P.; Verkhivker, G. Markov State Models and Perturbation-Based Approaches Reveal Distinct Dynamic Signatures and Hidden Allosteric Pockets in the Emerging SARS-Cov-2 Spike Omicron Variant Complexes with the Host Receptor: The Interplay of Dynamics and Convergent Evolution Modulates Allostery and Functional Mechanisms. J. Chem. Inf. Model. 2023, 63, 5272–5296. doi: 10.1021/acs.jcim.3c00778.

(72) Gan, H.H. Twaddle, A.; Marchand, B.; Gunsalus, K.C. Structural Modeling of the SARS-CoV-2 Spike/Human ACE2 Complex Interface can Identify High-Affinity Variants Associated with Increased Transmissibility. J. Mol. Biol. 2021, 433, 167051. doi: 10.1016/j.jmb.2021.167051.

(73) Gan, H. H.; Zinno, J.; Piano, F.; Gunsalus, K. C. Omicron Spike Protein Has a Positive Electrostatic Surface That Promotes ACE2 Recognition and Antibody Escape. Front. Virol. 2022, 2. 10.3389/fviro.2022.894531.

(74) Barroso da Silva, F. L.; Giron, C. C.; Laaksonen, A. Electrostatic Features for the Receptor Binding Domain of SARS-COV-2 Wildtype and Its Variants. Compass to the Severity of the Future Variants with the Charge-Rule. J. Phys. Chem. B. 2022, 126, 6835–6852. doi: 10.1021/acs.jpcb.2c04225.

(75) Hristova, S. H.; Zhivkov, A. M. Omicron Coronavirus: pH-Dependent Electrostatic Potential and Energy of Association of Spike Protein to ACE2 Receptor. Viruses 2023, 15, 1752. doi: 10.3390/v15081752.

(76) Verkhivker, G.; Alshahrani, M.; Gupta, G. Exploring Conformational Landscapes and Cryptic Binding Pockets in Distinct Functional States of the SARS-CoV-2 Omicron BA.1 and BA.2 Trimers: Mutation-Induced Modulation of Protein Dynamics and Network-Guided Prediction of Variant-Specific Allosteric Binding Sites. Viruses 2023, 15, 2009. doi: 10.3390/v15102009.

(77) Alshahrani, M.; Gupta, G.; Xiao, S.; Tao, P.; Verkhivker, G. Comparative Analysis of Conformational Dynamics and Systematic Characterization of Cryptic Pockets in the SARS-CoV-2 Omicron BA.2, BA.2.75 and XBB.1 Spike Complexes with the ACE2 Host Receptor: Confluence of Binding and Structural Plasticity in Mediating Networks of Conserved Allosteric Sites. Viruses 2023, 15, 2073. doi: 10.3390/v15102073.

(78) Jumper, J.; Evans, R.; Pritzel, A.; Green, T.; Figurnov, M.; Ronneberger, O.; Tunyasuvunakool, K.; Bates, R.; Žídek, A.; Potapenko, A.; Bridgland, A.; Meyer, C.; Kohl, S. A. A.; Ballard, A. J.; Cowie, A.; Romera-Paredes, B.; Nikolov, S.; Jain, R.; Adler, J.; Back, T.; Petersen, S.; Reiman, D.; Clancy, E.; Zielinski, M.; Steinegger, M.; Pacholska, M.; Berghammer, T.; Bodenstein, S.; Silver, D.; Vinyals, O.; Senior, A. W.; Kavukcuoglu, K.; Kohli, P.; Hassabis, D. Highly Accurate Protein Structure Prediction with AlphaFold. Nature 2021, 596, 583–589. doi: 10.1038/s41586-021-03819-2.

(79) Tunyasuvunakool, K.; Adler, J.; Wu, Z.; Green, T.; Zielinski, M.; Žídek, A.; Bridgland, A.; Cowie, A.; Meyer, C.; Laydon, A.; Velankar, S.; Kleywegt, G. J.; Bateman, A.; Evans, R.; Pritzel, A.; Figurnov, M.; Ronneberger, O.; Bates, R.; Kohl, S. A. A.; Potapenko, A.; Ballard, A. J.; Romera-Paredes, B.; Nikolov, S.; Jain, R.; Clancy, E.; Reiman, D.; Petersen, S.; Senior, A. W.; Kavukcuoglu, K.; Birney, E.; Kohli, P.; Jumper, J.; Hassabis, D. Highly Accurate Protein Structure Prediction for the Human Proteome. Nature 2021, 596, 590–596. doi: 10.1038/s41586-021-03828-1.

(80) Lin, Z.; Akin, H.; Rao, R.; Hie, B.; Zhu, Z.; Lu, W.; Smetanin, N.; Verkuil, R.; Kabeli, O.; Shmueli, Y.; dos Santos Costa, A.; Fazel-Zarandi, M.; Sercu, T.; Candido, S.; Rives, A. Evolutionary-Scale Prediction of Atomic-Level Protein Structure with a Language Model. Science 2023, 379, 1123–1130. doi: 10.1126/science.ade2574.

(81) Del Alamo, D.; Sala, D.; Mchaourab, H.S.; Meiler, J. Sampling alternative conformational states of transporters and receptors with AlphaFold2. Elife 2022, 11, e75751. doi: 10.7554/eLife.75751.

(82) Stein, R. A.; Mchaourab, H. S. SPEACH_AF: Sampling Protein Ensembles and Conformational Heterogeneity with Alphafold2. PLoS Comput Biol. 2022, 18, e1010483. doi: 10.1371/journal.pcbi.1010483.

(83) Chakravarty, D.; Porter, L. L. AlphaFold2 fails to predict protein fold switching. Protein Sci. 2022, 31, e4353. doi: 10.1002/pro.4353.

(84) Wayment-Steele, H. K.; Ovchinnikov, S.; Colwell, L.; Kern, D. Prediction of Multiple Conformational States by Combining Sequence Clustering with AlphaFold2. bioRxiv 2022. doi: 10.1101/2022.10.17.512570.

(85) Ester, M.; Kriegel, H.-P., Sander, J.; Xu, X.. A density-based algorithm for discovering clusters in large spatial databases with noise. International Conference on Knowledge Discovery in Databases and Data Mining (KDD-96), 1996, 226–231.

(86) Mirdita, M.; Schütze, K.; Moriwaki, Y.; Heo, L.; Ovchinnikov, S.; Steinegger, M. ColabFold: Making Protein Folding Accessible to All. Nat. Methods 2022, 19, 679–682. doi: 10.1038/s41592-022-01488-1.

(87) Zhang, Y. TM-Align: A Protein Structure Alignment Algorithm Based on the TM-Score. Nucleic Acids Res. 2005, 33, 2302–2309. doi: 10.1093/nar/gki524.

(88) Zemla, A. LGA: A Method for Finding 3D Similarities in Protein Structures. Nucleic Acids Res. 2003, 31, 3370–3374. doi: 10.1093/nar/gkg571.

(89) Hekkelman, M.L.; Te Beek, T.A.; Pettifer, S.R.; Thorne, D.; Attwood, T.K.; Vriend, G. WIWS: A protein structure bioinformatics web service collection. Nucleic Acids Res. 2010, 38, W719–W723. 10.1093/nar/gkq453.

(90) Fernandez-Fuentes, N.; Zhai, J.; Fiser, A. ArchPRED: A template based loop structure prediction server. Nucleic Acids Res. 2006, 34, W173–W176. 10.1093/nar/gkl113.

(91) Krivov, G.G.; Shapovalov, M.V.; Dunbrack, R.L., Jr. Improved prediction of protein side-chain conformations with SCWRL4. Proteins 2009, 77, 778–795. 10.1002/prot.22488.

(92) Søndergaard C. R.; Olsson M. H.; Rostkowski M.; Jensen J. H. Improved treatment of ligands and coupling effects in empirical calculation and rationalization of pKa values. J. Chem. Theory Comput. 2011, 7, 2284–2295. 10.1021/ct200133y.

(93) Olsson M. H.; Søndergaard C. R.; Rostkowski M.; Jensen J. H. PROPKA3: consistent treatment of internal and surface residues in empirical pKa predictions. J. Chem. Theory Comput. 2011, 7, 525–537. 10.1021/ct100578z.

(94) Bhattacharya, D.; Cheng, J. 3Drefine: Consistent Protein Structure Refinement by Optimizing Hydrogen Bonding Network and Atomic-Level Energy Minimization. Proteins 2013, 81, 119–131. doi: 10.1002/prot.24167.

(95) Bhattacharya, D.; Nowotny, J.; Cao, R.; Cheng, J. 3Drefine: An Interactive Web Server for Efficient Protein Structure Refinement. Nucleic Acids Res. 2016, 44, W406–W409. doi: 10.1093/nar/gkw336.

(96) Phillips, J.C.; Hardy, D.J.; Maia, J.D.C.; Stone, J.E.; Ribeiro, J.V.; Bernardi, R.C.; Buch, R.; Fiorin, G.; Hénin, J.; Jiang, W.;, et al. Scalable Molecular Dynamics on CPU and GPU Architectures with NAMD. J. Chem. Phys. 2020, 153, 044130. 10.1063/5.0014475.

(97) Huang, J.; Rauscher, S.; Nawrocki, G.; Ran, T.; Feig, M.; de Groot, B.L.; Grubmüller, H.; MacKerell, A.D., Jr. CHARMM36m: An improved force field for folded and intrinsically disordered proteins. Nat. Methods 2017, 14, 71–73. 10.1038/nmeth.4067.

(98) Fernandes, H.S.; Sousa, S.F.; Cerqueira, N.M.F.S.A. VMD Store-A VMD Plugin to Browse, Discover, and Install VMD Extensions. J. Chem. Inf. Model. 2019, 59, 4519–4523. doi: 10.1021/acs.jcim.9b00739.

(99) Jo, S.; Kim, T.; Iyer, V. G.; Im, W. CHARMMLJGUI: A WebLJbased Graphical User Interface for CHARMM. J Comput Chem. 2008, 29, 1859–1865. doi: 10.1002/jcc.20945.

(100) Lee, J.; Cheng, X.; Swails, J. M.; Yeom, M. S.; Eastman, P. K.; Lemkul, J. A.; Wei, S.; Buckner, J.; Jeong, J. C.; Qi, Y.; Jo, S.; Pande, V. S.; Case, D. A.; Brooks, C. L., III; MacKerell, A. D., Jr.; Klauda, J. B.; Im, W. CHARMM-GUI Input Generator for NAMD, GROMACS, AMBER, OpenMM, and CHARMM/OpenMM Simulations Using the CHARMM36 Additive Force Field. J Chem Theory Comput. 2016, 12, 405–413. doi: 10.1021/acs.jctc.5b00935.

(101) Jorgensen, W.L.; Chandrasekhar, J.; Madura, J.D.; Impey, R.W.; Klein, M.L. Comparison of Simple Potential Functions for Simulating Liquid Water. J. Chem. Phys. 1983, 79, 926–935. 10.1063/1.445869.

(102) Ross, G.A.; Rustenburg, A.S.; Grinaway, P.B.; Fass, J.; Chodera, J.D. Biomolecular Simulations under Realistic Macroscopic Salt Conditions. J. Phys. Chem. B 2018, 122, 5466– 5486. 10.1021/acs.jpcb.7b11734.

(103) Di Pierro, M.; Elber, R.; Leimkuhler, B. A Stochastic Algorithm for the Isobaric-Isothermal Ensemble with Ewald Summations for All Long Range Forces. J. Chem. Theory Comput. 2015, 11, 5624–5637. 10.1021/acs.jctc.5b00648.

(104) Martyna, G.J.; Tobias, D.J.; Klein, M.L. Constant pressure molecular dynamics algorithms. J. Chem. Phys. 1994, 101, 4177–4189. 10.1063/1.467468.

(105) Feller, S.E.; Zhang, Y.; Pastor, R.W.; Brooks, B.R. Constant pressure molecular dynamics simulation: The Langevin piston method. J. Chem. Phys. 1995, 103, 4613–4621. 10.1063/1.470648.

(106) Davidchack, R.L.; Handel, R.; Tretyakov, M.V. Langevin thermostat for rigid body dynamics. J. Chem. Phys. 2009, 130, 234101. 10.1063/1.3149788.

(107) Sacquin-Mora, S.; Lavery, R., Investigating the local flexibility of functional residues in hemoproteins. Biophys. J. 2006, 90, 2706–2717

(108) Sacquin-Mora, S.; Laforet, E.; Lavery, R., Locating the active sites of enzymes using mechanical properties. Proteins 2007, 67, 350–359.

(109) Hou, T.; Wang, J.; Li, Y.; Wang, W. Assessing the Performance of the MM/PBSA and MM/GBSA Methods. 1. The Accuracy of Binding Free Energy Calculations Based on Molecular Dynamics Simulations. J. Chem. Inf. Model. 2011, 51, 69–82. 10.1021/ci100275a.

(110) Sun, H.; Li, Y.; Tian, S.; Xu, L.; Hou, T. Assessing the Performance of MM/PBSA and MM/GBSA Methods. 4. Accuracies of MM/PBSA and MM/GBSA Methodologies Evaluated by Various Simulation Protocols Using PDBbind Data Set. Phys. Chem. Chem. Phys. 2014, 16, 16719–16729. 10.1039/c4cp01388c.

(111) Dehouck, Y.; Kwasigroch, J. M.; Rooman, M.; Gilis, D. BeAtMuSiC: Prediction of changes in protein-protein binding affinity on mutations. Nucleic Acids Res. 2013, 41, W333–W339. doi: 10.1093/nar/gkt450.

(112) Dehouck, Y.; Gilis, D.; Rooman, M. A new generation of statistical potentials for proteins. Biophys. J. 2006, 90, 4010–4017. doi: 10.1529/biophysj.105.079434.

(113) Dehouck, Y.; Grosfils, A.; Folch, B.; Gilis, D.; Bogaerts, P.; Rooman, M. Fast and accurate predictions of protein stability changes upon mutations using statistical potentials and neural networks: PoPMuSiC-2.0. Bioinformatics 2009, 25, 2537–2543. doi: 10.1093/bioinformatics/btp445.

(114) Baker N. A.; Sept D.; Joseph S.; Holst M. J.; McCammon J. A. Electrostatics of Nanosystems: Application to Microtubules and the Ribosome. Proc. Natl. Acad. Sci. U.S.A. 2001, 98, 10037–10041. doi:10.1073/pnas.181342398.

(115) Dolinsky T. J.; Nielsen J. E.; McCammon J. A.; Baker N. A. PDB2PQR: An Automated Pipeline for the Setup, Execution, and Analysis of Poisson-Boltzmann Electrostatics Calculations. Nucleic Acids Res. 2004, 32, W665–W667. doi:10.1093/nar/gkh381.

(116) Jurrus, E.; Engel, D.; Star, K.; Monson, K.; Brandi, J.; Felberg, L.E.; Brookes, D.H.; Wilson, L.; Chen, J.; Liles, K.; Chun, M.; Li, P.; Gohara, D.W.; Dolinsky, T.; Konecny, R.; Koes, D.R.; Nielsen, J.E.; Head-Gordon, T.; Geng, W.; Krasny, R.; Wei, G-W.; Holst, M.J.; McCammon, J.A.; Baker, N.A. Improvements to the APBS biomolecular solvation software suite. Protein Sci. 2018, 27, 112–128.

(117) Costello, S. M.; Shoemaker, S. R.; Hobbs, H. T.; Nguyen, A. W.; Hsieh, C.-L.; Maynard, J. A.; McLellan, J. S.; Pak, J. E.; Marqusee, S. The SARS-CoV-2 Spike Reversibly Samples an Open-Trimer Conformation Exposing Novel Epitopes. Nat Struct Mol Biol. 2022, 29, 229–238. doi: 10.1038/s41594-022-00735-5.

(118) Calvaresi, V.; Wrobel, A. G.; Toporowska, J.; Hammerschmid, D.; Doores, K. J.; Bradshaw, R. T.; Parsons, R. B.; Benton, D. J.; Roustan, C.; Reading, E.; Malim, M. H.; Gamblin, S. J.; Politis, A. Structural Dynamics in the Evolution of SARS-CoV-2 Spike Glycoprotein. Nat Commun. 2023, 14, 1421. doi: 10.1038/s41467-023-36745-0.

(119) Braet, S. M.; Buckley, T. S.; Venkatakrishnan, V.; Dam, K.-M. A.; Bjorkman, P. J.; Anand, G. S. Timeline of Changes in Spike Conformational Dynamics in Emergent SARS-CoV- 2 Variants Reveal Progressive Stabilization of Trimer Stalk with Altered NTD Dynamics. Elife 2023, 12, e82584. doi: 10.7554/eLife.82584.

(120) Starr, T. N.; Greaney, A. J.; Hilton, S. K.; Ellis, D.; Crawford, K. H. D.; Dingens, A. S.; Navarro, M. J.; Bowen, J. E.; Tortorici, M. A.; Walls, A. C.; King, N. P.; Veesler, D.; Bloom, J. D. Deep Mutational Scanning of SARS-CoV-2 Receptor Binding Domain Reveals Constraints on Folding and ACE2 Binding. Cell 2020, 182, 1295–1310.e20. doi: 10.1016/j.cell.2020.08.012.

(121) Starr, T. N.; Greaney, A. J.; Stewart, C. M.; Walls, A. C.; Hannon, W. W.; Veesler, D.; Bloom, J. D. Deep Mutational Scans for ACE2 Binding, RBD Expression, and Antibody Escape in the SARS-CoV-2 Omicron BA.1 and BA.2 Receptor-Binding Domains. PLoS Pathog. 2022, 18, e1010951. doi: 10.1371/journal.ppat.1010951.

(122) Dadonaite, B.; Crawford, K. H. D.; Radford, C. E.; Farrell, A. G.; Yu, T. C.; Hannon, W. W.; Zhou, P.; Andrabi, R.; Burton, D. R.; Liu, L.; Ho, D. D.; Chu, H. Y.; Neher, R. A.; Bloom, J. D. A Pseudovirus System Enables Deep Mutational Scanning of the Full SARS-CoV-2 Spike. Cell 2023, 186, 1263–1278.e20. doi: 10.1016/j.cell.2023.02.001.

(123) Williams, J. K.; Wang, B.; Sam, A.; Hoop, C. L.; Case, D. A.; Baum, J. Molecular Dynamics Analysis of a Flexible Loop at the Binding Interface of the SARS-CoV-2 spike protein receptor-binding domain. Proteins 2022, 90, 1044–1053. doi: 10.1002/prot.26208.

(124) Focosi, D.; Quiroga, R.; McConnell, S.; Johnson, M.C.; Casadevall, A. Convergent Evolution in SARS-CoV-2 Spike Creates a Variant Soup from Which New COVID-19 Waves Emerge. Int. J. Mol. Sci. 2023, 24, 2264. 10.3390/ijms24032264.

(125) Cao, Y.; Jian, F.; Wang, J.; Yu, Y.; Song, W.; Yisimayi, A.; Wang, J.; An, R.; Chen, X.; Zhang, N.; Wang, Y.; Wang, P.; Zhao, L.; Sun, H.; Yu, L.; Yang, S.; Niu, X.; Xiao, T.; Gu, Q.; Shao, F.; Hao, X.; Xu, Y.; Jin, R.; Shen, Z.; Wang, Y.; Xie, X. S. Imprinted SARS-CoV-2 Humoral Immunity Induces Convergent Omicron RBD Evolution. Nature 2023, 614, 521–529. doi: 10.1038/s41586-022-05644-7.

(126) Li, Y.; Ren, C.; Shen, Y.; Zhang, Y.; Chen, J.; Zheng, J.; Tian, R.; Cao, L.; Yan, R. Cryo-EM Structures of SARS-CoV-2 BA.2-Derived Subvariants Spike in Complex with ACE2 Receptor. Cell Discov. 2023, 9, 108. doi: 10.1038/s41421-023-00607-2.

(127) Philip, A. M.; Ahmed, W. S.; Biswas, K. H. Reversal of the Unique Q493R Mutation Increases the Affinity of Omicron S1-RBD for ACE2. Comput Struct Biotechnol J. 2023, 21, 1966–1977. doi: 10.1016/j.csbj.2023.02.019.

(128) Jian, F.; Feng, L.; Yang, S.; Yu, Y.; Wang, L.; Song, W.; Yisimayi, A.; Chen, X.; Xu, Y.; Wang, P.; Yu, L.; Wang, J.; Liu, L.; Niu, X.; Wang, J.; Xiao, T.; An, R.; Wang, Y.; Gu, Q.; Shao, F.; Jin, R.; Shen, Z.; Wang, Y.; Wang, X.; Cao, Y. Convergent Evolution of SARS-CoV-2 XBB Lineages on Receptor-Binding Domain 455-456 Synergistically Enhances Antibody Evasion and ACE2 Binding, bioRxiv 2023. doi: 10.1101/2023.08.30.555211.

(129) Ma, W.; Fu, H.; Jian, F.; Cao, Y.; Li, M. Immune Evasion and ACE2 Binding Affinity Contribute to SARS-CoV-2 Evolution. Nat Ecol Evol. 2023, 7, 1457–1466. doi: 10.1038/s41559-023-02123-8.

(130) Neverov, A.D.; Fedonin, G.; Popova, A.; Bykova, D.; Bazykin, G. Coordinated Evolution at Amino Acid Sites of SARS-CoV-2 Spike. Elife 2023, 12, e82516. 10.7554/eLife.82516.

(131) Park, Y.-J.; De Marco, A.; Starr, T. N.; Liu, Z.; Pinto, D.; Walls, A. C.; Zatta, F.; Zepeda, S. K.; Bowen, J. E.; Sprouse, K. R.; Joshi, A.; Giurdanella, M.; Guarino, B.; Noack, J.; Abdelnabi, R.; Foo, S.-Y. C.; Rosen, L. E.; Lempp, F. A.; Benigni, F.; Snell, G.; Neyts, J.; Whelan, S. P. J.; Virgin, H. W.; Bloom, J. D.; Corti, D.; Pizzuto, M. S.; Veesler, D. Antibody-Mediated Broad Sarbecovirus Neutralization through ACE2 Molecular Mimicry. Science 2022, 375, 449–454. 10.1126/science.abm8143.

(132) Nutalai, R.; Zhou, D.; Tuekprakhon, A.; Ginn, H. M.; Supasa, P.; Liu, C.; Huo, J.; Mentzer, A. J.; Duyvesteyn, H. M. E.; Dijokaite-Guraliuc, A.; Skelly, D.; Ritter, T. G.; Amini, A.; Bibi, S.; Adele, S.; Johnson, S. A.; Constantinides, B.; Webster, H.; Temperton, N.; Klenerman, P.; Barnes, E.; Dunachie, S. J.; Crook, D.; Pollard, A. J.; Lambe, T.; Goulder, P.; Paterson, N. G.; Williams, M. A.; Hall, D. R.; Mongkolsapaya, J.; Fry, E. E.; Dejnirattisai, W.; Ren, J.; Stuart, D. I.; Screaton, G. R. Potent Cross-Reactive Antibodies Following Omicron Breakthrough in Vaccinees. Cell 2022, 185, 2116–2131.e18. doi: 10.1016/j.cell.2022.05.014.

(133) Cao, Y.; Yisimayi, A.; Bai, Y.; Huang, W.; Li, X.; Zhang, Z.; Yuan, T.; An, R.; Wang, J.; Xiao, T.; Du, S.; Ma, W.; Song, L.; Li, Y.; Li, X.; Song, W.; Wu, J.; Liu, S.; Li, X.; Zhang, Y.; Su, B.; Guo, X.; Wei, Y.; Gao, C.; Zhang, N.; Zhang, Y.; Dou, Y.; Xu, X.; Shi, R.; Lu, B.; Jin, R.; Ma, Y.; Qin, C.; Wang, Y.; Feng, Y.; Xiao, J.; Xie, X. S. Humoral Immune Response to Circulating SARS-CoV-2 Variants Elicited by Inactivated and RBD-Subunit Vaccines. Cell Res. 2021, 31, 732–741. doi: 10.1038/s41422-021-00514-9.

(134) Parzych, E. M.; Du, J.; Ali, A. R.; Schultheis, K.; Frase, D.; Smith, T. R. F.; Cui, J.; Chokkalingam, N.; Tursi, N. J.; Andrade, V. M.; Warner, B. M.; Gary, E. N.; Li, Y.; Choi, J.; Eisenhauer, J.; Maricic, I.; Kulkarni, A.; Chu, J. D.; Villafana, G.; Rosenthal, K.; Ren, K.; Francica, J. R.; Wootton, S. K.; Tebas, P.; Kobasa, D.; Broderick, K. E.; Boyer, J. D.; Esser, M. T.; Pallesen, J.; Kulp, D. W.; Patel, A.; Weiner, D. B. DNA-Delivered Antibody Cocktail Exhibits Improved Pharmacokinetics and Confers Prophylactic Protection against SARS-CoV-2. Nat Commun. 2022, 13, 5886. doi: 10.1038/s41467-022-33309-6.

(135) Wang, K.; Jia, Z.; Bao, L.; Wang, L.; Cao, L.; Chi, H.; Hu, Y.; Li, Q.; Zhou, Y.; Jiang, Y.; Zhu, Q.; Deng, Y.; Liu, P.; Wang, N.; Wang, L.; Liu, M.; Li, Y.; Zhu, B.; Fan, K.; Fu, W.; Yang, P.; Pei, X.; Cui, Z.; Qin, L.; Ge, P.; Wu, J.; Liu, S.; Chen, Y.; Huang, W.; Wang, Q.; Qin, C.-F.; Wang, Y.; Qin, C.; Wang, X. Memory B Cell Repertoire from Triple Vaccinees against Diverse SARS-CoV-2 Variants. Nature 2022, 603, 919–925. doi: 10.1038/s41586-022-04466-x.

(136) Wang, L.; Fu, W.; Bao, L.; Jia, Z.; Zhang, Y.; Zhou, Y.; Wu, W.; Wu, J.; Zhang, Q.; Gao, Y.; Wang, K.; Wang, Q.; Qin, C.; Wang, X. Selection and Structural Bases of Potent Broadly Neutralizing Antibodies from 3-Dose Vaccinees That Are Highly Effective against Diverse SARS-CoV-2 Variants, Including Omicron Sublineages. Cell Res. 2022, 32, 691–694. doi: 10.1038/s41422-022-00677-z.

(137) Zhou, B.; Zhou, R.; Tang, B.; Chan, J. F.-W.; Luo, M.; Peng, Q.; Yuan, S.; Liu, H.; Mok, B. W.-Y.; Chen, B.; Wang, P.; Poon, V. K.-M.; Chu, H.; Chan, C. C.-S.; Tsang, J. O.-L.; Chan, C. C.-Y.; Au, K.-K.; Man, H.-O.; Lu, L.; To, K. K.-W.; Chen, H.; Yuen, K.-Y.; Dang, S.; Chen, Z. A Broadly Neutralizing Antibody Protects Syrian Hamsters against SARS-CoV-2 Omicron Challenge. Nat Commun. 2022, 13, 3589. doi: 10.1038/s41467-022-31259-7.

(138) Zhou, T.; Wang, L.; Misasi, J.; Pegu, A.; Zhang, Y.; Harris, D.R.; Olia, A.S.; Talana, C.A.; Yang, E.S.; Chen, M.; Choe, M.; Shi, W.; Teng, I.T.; Creanga, A.; Jenkins, C.; Leung, K.; Liu, T.; Stancofski, E.D.; Stephens, T.; Zhang, B.; Tsybovsky, Y.; Graham, B.S.; Mascola, J.R.; Sullivan, N.J.; Kwong, P.D. Structural basis for potent antibody neutralization of SARS-CoV-2 variants including B.1.1.529. Science 2022, 376, eabn8897. doi: 10.1126/science.abn8897.

(139) Zhao, Z.; Zhou, J.; Tian, M.; Huang, M.; Liu, S.; Xie, Y.; Han, P.; Bai, C.; Han, P.; Zheng, A.; Fu, L.; Gao, Y.; Peng, Q.; Li, Y.; Chai, Y.; Zhang, Z.; Zhao, X.; Song, H.; Qi, J.; Wang, Q.; Wang, P.; Gao, G. F. Omicron SARS-CoV-2 Mutations Stabilize Spike up-RBD Conformation and Lead to a Non-RBM-Binding Monoclonal Antibody Escape. Nat Commun. 2022, 13, 4958. doi: 10.1038/s41467-022-32665-7.

(140) Westendorf, K.; Žentelis, S.; Wang, L.; Foster, D.; Vaillancourt, P.; Wiggin, M.; Lovett, E.; van der Lee, R.; Hendle, J.; Pustilnik, A.; Sauder, J. M.; Kraft, L.; Hwang, Y.; Siegel, R. W.; Chen, J.; Heinz, B. A.; Higgs, R. E.; Kallewaard, N. L.; Jepson, K.; Goya, R.; Smith, M. A.; Collins, D. W.; Pellacani, D.; Xiang, P.; de Puyraimond, V.; Ricicova, M.; Devorkin, L.; Pritchard, C.; O’Neill, A.; Dalal, K.; Panwar, P.; Dhupar, H.; Garces, F. A.; Cohen, C. A.; Dye, J. M.; Huie, K. E.; Badger, C. V.; Kobasa, D.; Audet, J.; Freitas, J. J.; Hassanali, S.; Hughes, I.; Munoz, L.; Palma, H. C.; Ramamurthy, B.; Cross, R. W.; Geisbert, T. W.; Menachery, V.; Lokugamage, K.; Borisevich, V.; Lanz, I.; Anderson, L.; Sipahimalani, P.; Corbett, K. S.; Yang, E. S.; Zhang, Y.; Shi, W.; Zhou, T.; Choe, M.; Misasi, J.; Kwong, P. D.; Sullivan, N. J.; Graham, B. S.; Fernandez, T. L.; Hansen, C. L.; Falconer, E.; Mascola, J. R.; Jones, B. E.; Barnhart, B. C. LY-CoV1404 (Bebtelovimab) Potently Neutralizes SARS-CoV-2 Variants. Cell Rep. 2022, 39, 110812. doi: 10.1016/j.celrep.2022.110812.

(141) Liu, C.; Zhou, D.; Nutalai, R.; Duyvesteyn, H. M. E.; Tuekprakhon, A.; Ginn, H. M.; Dejnirattisai, W.; Supasa, P.; Mentzer, A. J.; Wang, B.; Case, J. B.; Zhao, Y.; Skelly, D. T.; Chen, R. E.; Johnson, S. A.; Ritter, T. G.; Mason, C.; Malik, T.; Temperton, N.; Paterson, N. G.; Williams, M. A.; Hall, D. R.; Clare, D. K.; Howe, A.; Goulder, P. J. R.; Fry, E. E.; Diamond, M. S.; Mongkolsapaya, J.; Ren, J.; Stuart, D. I.; Screaton, G. R. The Antibody Response to SARS-CoV-2 Beta Underscores the Antigenic Distance to Other Variants. Cell Host Microbe. 2022, 30, 53–68.e12. doi: 10.1016/j.chom.2021.11.013.

(142) Starr, T.N.; Greaney, A.J.; Hannon, W.W.; Loes, A.N.; Hauser, K.; Dillen, J.R.; Ferri, E.; Farrell, A.G.; Dadonaite, B.; McCallum, M.; Matreyek, K.A.; Corti, D.; Veesler, D.; Snell, G.; Bloom, J.D. Shifting mutational constraints in the SARS-CoV-2 receptor-binding domain during viral evolution. Science 2022, 377, 420–424. doi: 10.1126/science.abo7896.

(143) Moulana, A.; Dupic, T.; Phillips, A. M.; Chang, J.; Nieves, S.; Roffler, A. A.; Greaney, A. J.; Starr, T. N.; Bloom, J. D.; Desai, M. M. Compensatory Epistasis Maintains ACE2 Affinity in SARS-CoV-2 Omicron BA.1. Nat Commun. 2022, 13, 7011. doi: 10.1038/s41467-022-34506-z.

