## Supplemental Figures S1-S8, Table S1 for "Accurate Characterization of Conformational Ensembles and Binding Mechanisms of the SARS-CoV-2 Omicron BA.2 and BA.2.86 Spike Protein with the Host Receptor and Distinct Classes of Antibodies Using AlphaFold2-Augmented Integrative Computational Modeling"

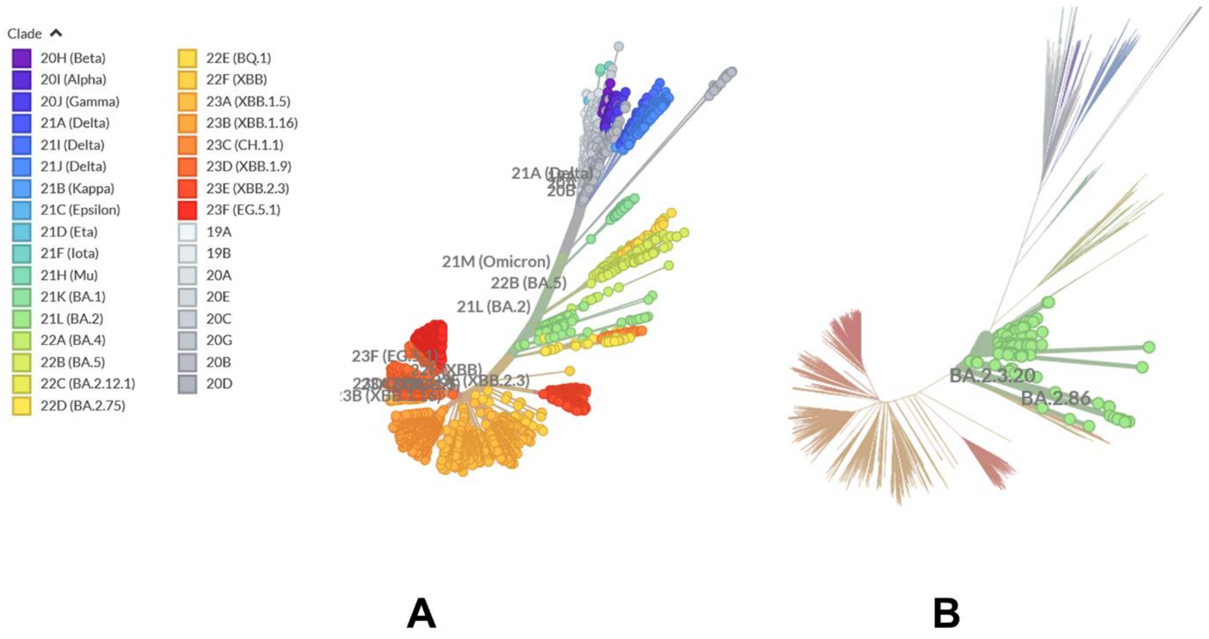


**Figure S1.** (A) An overview of the phylogenetic analysis of BA.2.86 and BA.2 variants. (B) Divergence of BA.2.86 spike sequence from major SARS-CoV-2 variants. The depicted phylogeny reflected genomic epidemiology of SARS-CoV-2 with subsampling focused globally over the past 6 months, showing 3639 of 3639 genomes sampled between Dec 2019 and Nov 2023. The graphs are generated using Nextstrain, an open-source project for real time tracking of evolving pathogen populations (https://nextstrain.org/).[50]

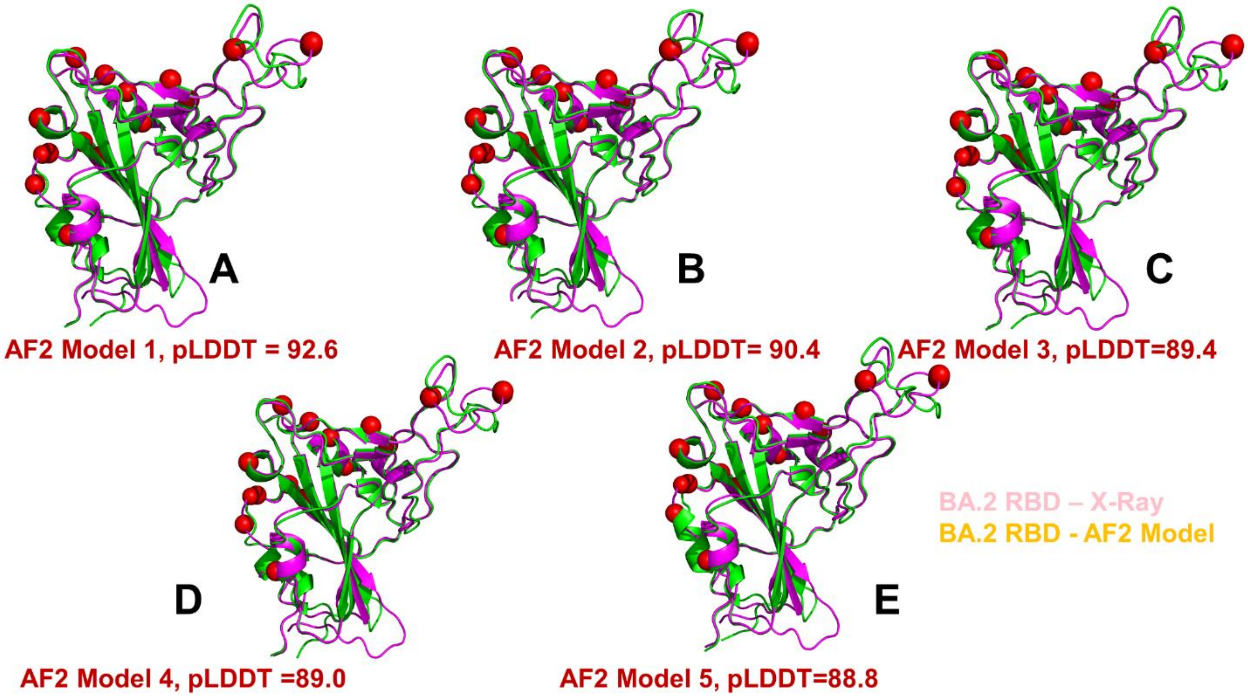


**Figure S2.** The alignment of top five BA.2 RBD models with the crystallographic conformation of the BA.2 RBD. The top ranked model (A), second ranked model (B), third ranked model (C), fourth ranked model (D) and fifth ranked model (E). The crystallographic BA.2 RBD conformation (pdb id 7XB0) is shown in magenta ribbons. The AF2 predicted BA.2 RBD models are shown in green ribbons. The pLDDT values associated with each of the top five modes are shown on the respective panels (A-E). The positions of BA.2 RBD mutations are shown in red spheres.

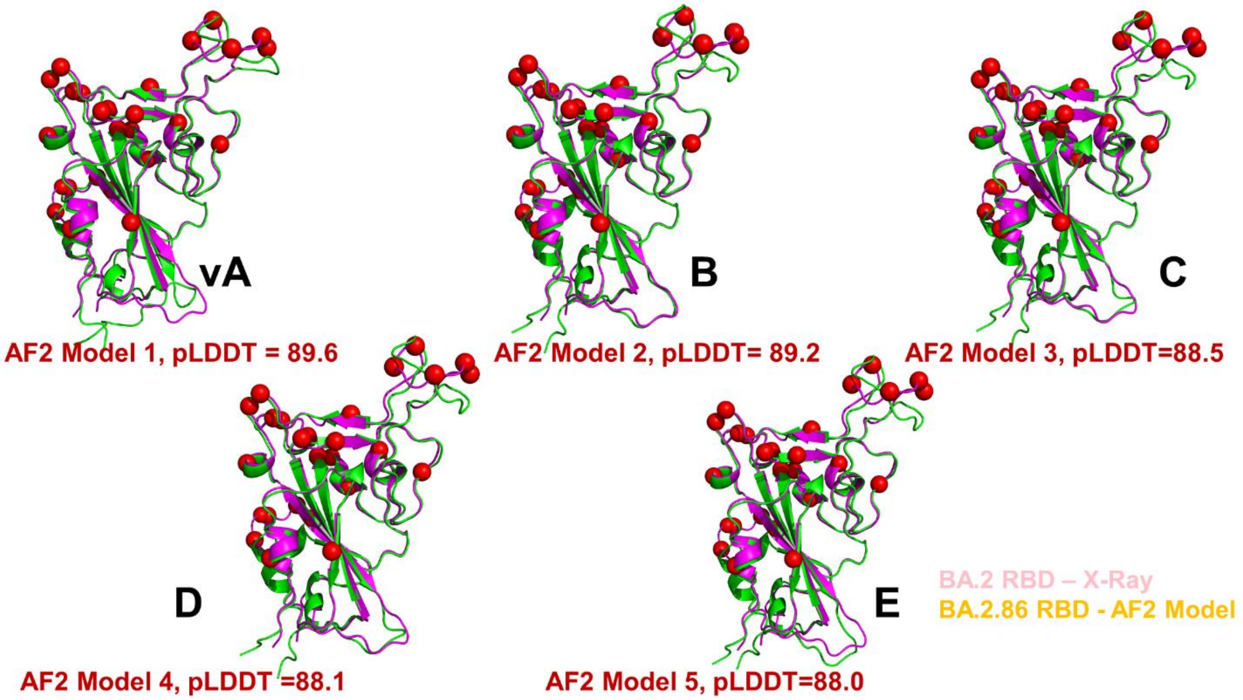


**Figure S3.** The alignment of top five BA.2.86 RBD models with the crystallographic conformation of the BA.2 RBD. The top ranked model (A), second ranked model (B), third ranked model (C), fourth ranked model (D) and fifth ranked model (E). The crystallographic BA.2 RBD conformation (pdb id 7XB0) is shown in magenta ribbons. The AF2 predicted BA.2.86 RBD models are shown in green ribbons. The pLDDT values associated with each of the top five modes are shown on the respective panels (A-E). The positions of BA.2.86 RBD mutations are shown in red spheres.

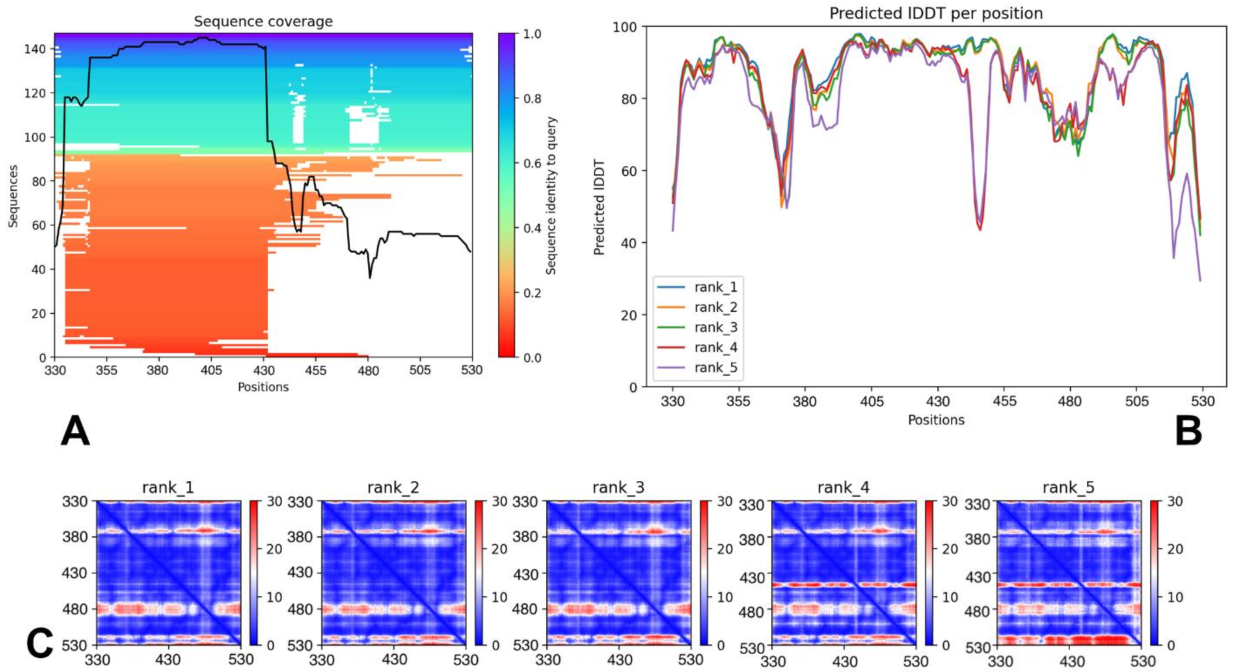


**Figure S4.** The post-processing AF2 analysis of predictions for the BA.2.86 RBD-ACE2 complex. (A) A heatmap representation of the MSA indicates all sequences mapped to the input sequences. The color scale points to the identity score, and sequences are ordered from top (largest identity) to bottom (lowest identity). White regions are not covered, which occurs with sub-sequence entries in the database. The black line qualifies the relative coverage of the sequence with respect to the total number of aligned sequences. (B) The pLDDT per residue for the top five models obtained from AF2 predictions of the BA.2.86 RBD-ACE2 complex. (C) The predicted alignment error (PAE) for the top five models obtained from AF2 predictions. These heat maps are provided for each final model and show the PAE between each residue in the model. The color scale contains three colors to highlight the contrast between the high confidence regions and the low confidence regions.

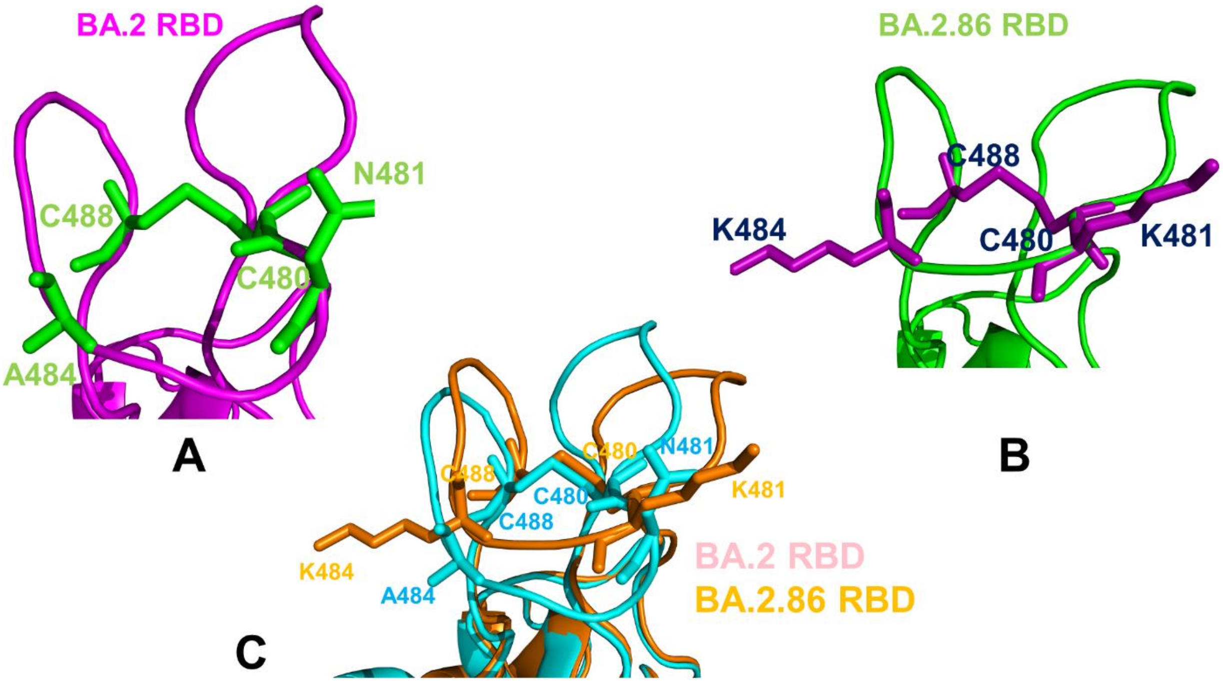


**Figure S5.** (A) A closeup of the disulfide bond in the RBD region of the BA.2 RBD-ACE2 complex. The C488-C480 disulfide bond and positions of N481 and A484 residues are shown in green sticks. The BA.2 RBD is in magenta-colored ribbons. (B) A closeup of the disulfide bond in the RBD region of the BA.2.86 RBD-ACE2 complex. The C488-C480 disulfide bond and positions of N481K and A484K residues are shown in purple sticks. The BA.2.86 RBD is in green ribbons. (C) Structural alignment of the C480-C488 disulfide bond in the BA.2 RBD (in cyan) and BA.2.86 (in orange). The positions of overlayed N481/K481 and A484/K484 are shown in respective cyan-colored (BA.2) and orange-colored sticks (BA.2.86).

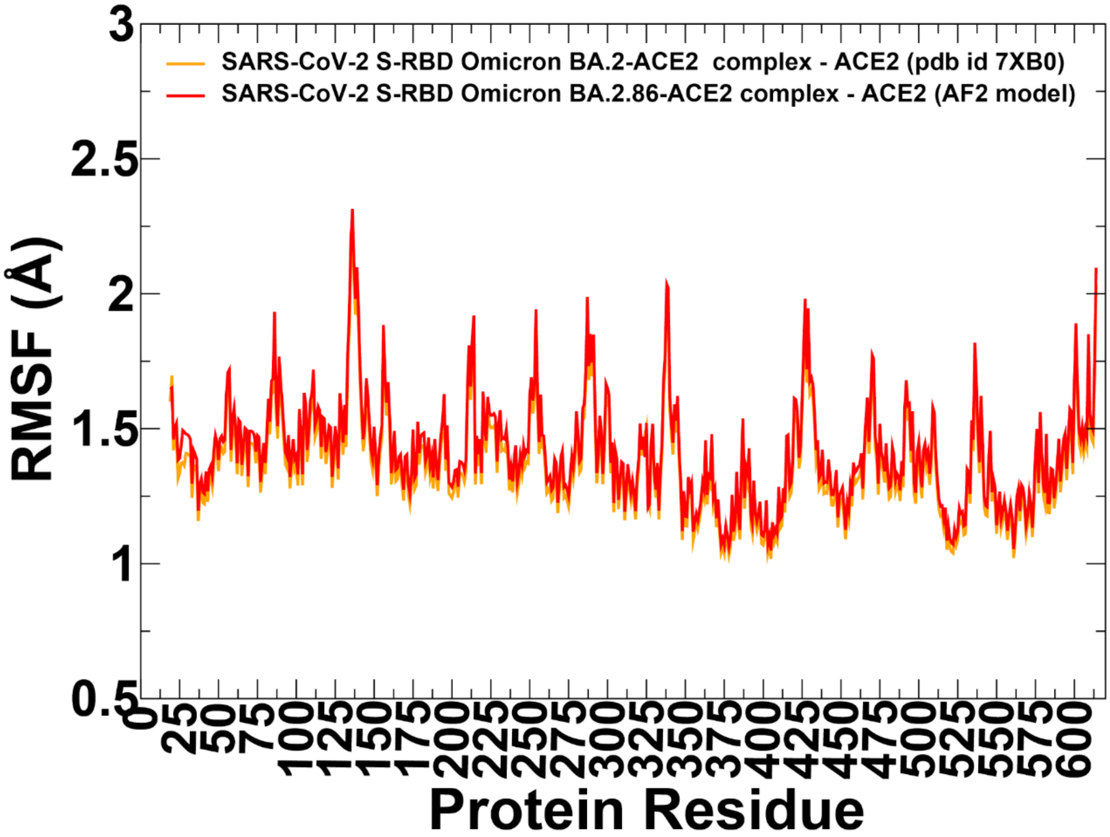


**Figure S6.** Conformational dynamics profiles obtained MD simulations of the BA.2 RBD-ACE2 and BA.2.86 RBD-ACE2 complexes. (A) The RMSF profiles for the ACE2 residues obtained from MD simulations of the BA.2 RBD-ACE2 complex, pdb id 7XB0 (in green lines), and BA.2.86 RB-ACE2 complex (in red lines).

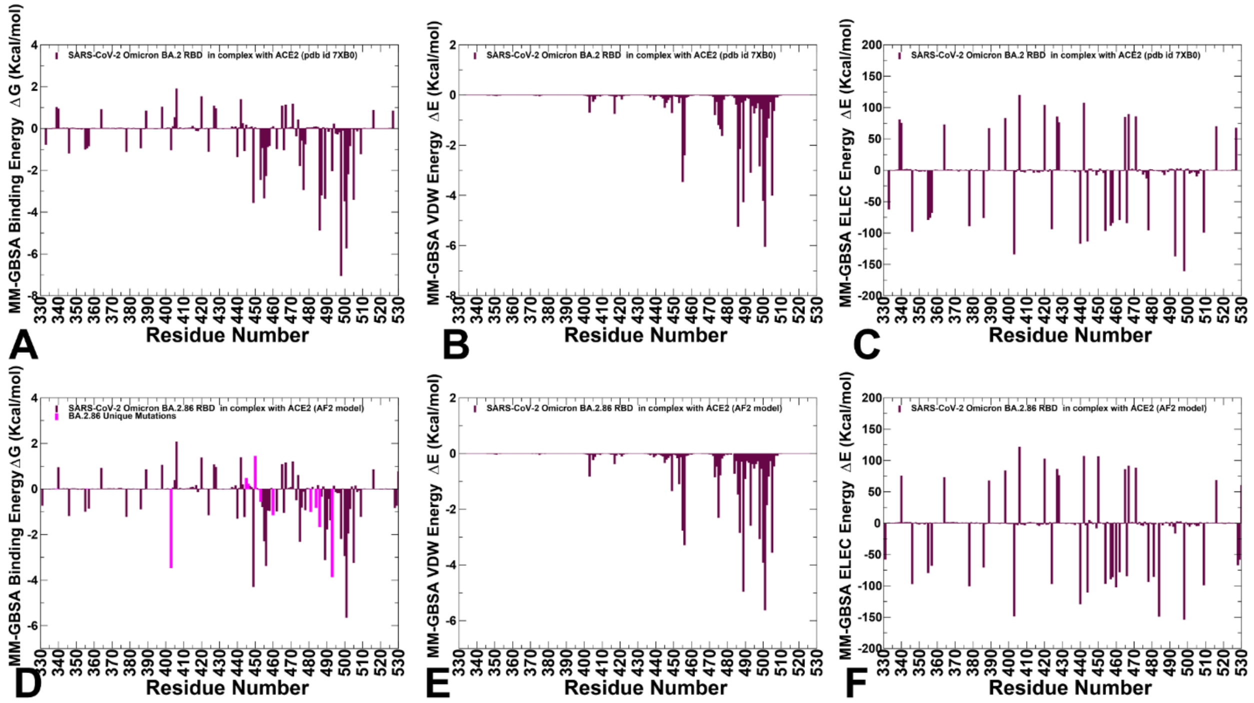


**Figure S7.** (A) The residue-based total MM-GBSA binding energy ΔG contribution for the BA.2 RBD-ACE2 complex. (B) The van der Waals contribution of the total binding energy for the BA.2 RBD-ACE2 complex. (C) The electrostatic contribution of the total binding energy for the BA.2 RBD-ACE2 complex. The residue-based total MM-GBSA binding energy ΔG contribution for the BA.2.86 RBD-ACE2 complex (D) , the van der Waals contribution of the BA.2.86 total binding energy (E) and the electrostatic contribution of the BA.2.86 total binding energy (F). Th MM-GBSA contributions are evaluated using 1,000 samples from the equilibrium MD simulations of the BA.2 and BA.2.86 RBD-ACE2 complexes. It is assumed that the entropy contributions for binding are relatively similar and are not considered in the analysis. The statistical errors was estimated on the basis of the deviation between block average and are in the range of 0.2-1.6 kcal/mol.

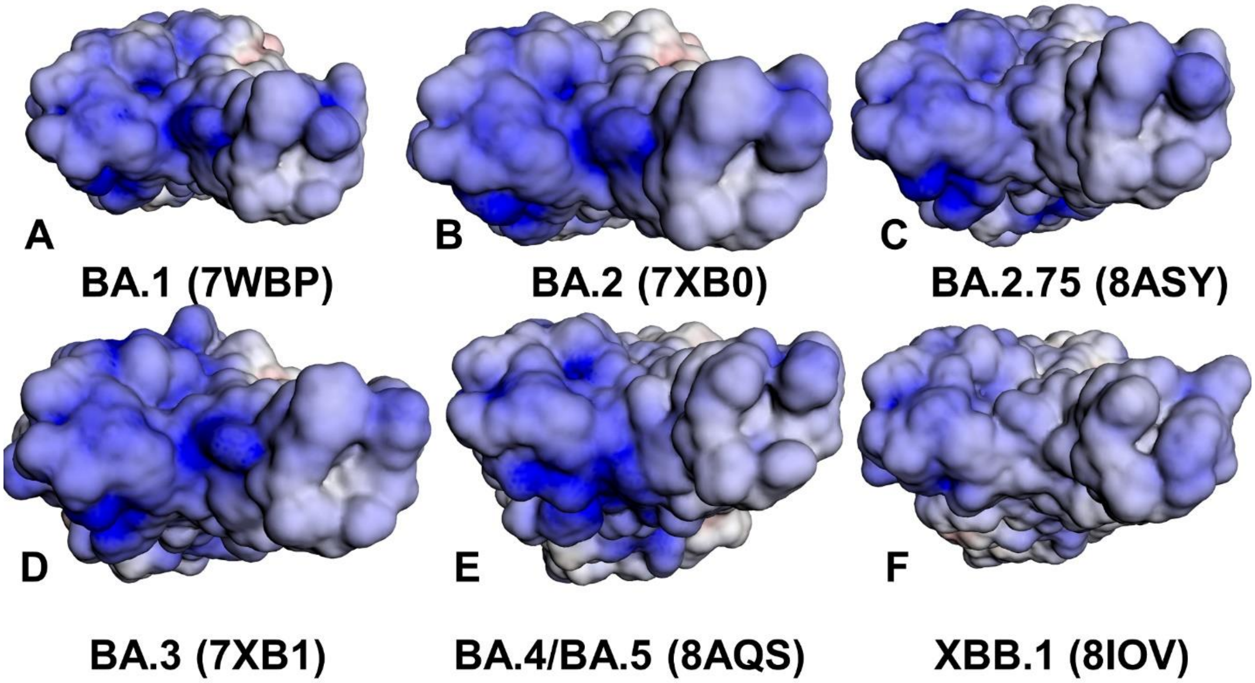


**Figure S8.** The distribution of the electrostatic potentials on the molecular surface of the S-RBD-ACE2 complexes for BA.1 RBD, pdb id 7WBP (A), BA.2 RBD, pdb id 7XB0 (B), BA.2.75 RBD, pb id 8ASY (C), BA.3 RBD, pdb id 7XB1 (D), BA.4/BA.5 RBD, pdb id 8AQS (E), and XBB.1 RBD, pdb id 8IOV (F). The crystal structures of the Omicron RBD-ACE2 complexes are used in computations of the electrostatic potentials. The color scale of the electrostatic potential surface is in units of kT/e at T = 37°C. Electro-positively and electronegatively charged areas are colored in blue and red, respectively. Neutral residues are shown in white.

**Table S1.** MM-GBSA Binding Energies for the BA.2 and BA.2.86 RBD-ACE2 Complexes.

| **System** | $E_{vdW}$ | $E_{elec}$ | $E\_GB$ | $E\_SA$ | $\Delta G_{bind(kcal/mol)}$ | Experimental binding |
|---|---|---|---|---|---|---|
| BA.2.86 RBD-ACE2 | -107.7 | -1923.8 | 1954.0 | -13.27 | -90.7 | 0.6 nM |
| BA.2 RBD-ACE2 | -113.8 | -1474.1 | 1514.1 | -14.8 | -88.6 | 1.68 nM |
